# The impact of life stage and pigment source on the evolution of novel warning signal traits

**DOI:** 10.1101/2020.10.20.346700

**Authors:** Carita Lindstedt, Robin Bagley, Sara Calhim, Mackenzie Jones, Catherine Linnen

## Abstract

Our understanding of how novel color traits evolve in aposematic taxa is based largely on studies of reproductive stages and organisms with endogenously produced pigmentation. In these systems, genetic drift is often required for novel alleles to overcome strong purifying selection stemming from frequency-dependent predation and positive assortative mating. Here we show that the importance of these mechanisms can differ if selective processes are considered in larval stage instead. By integrating population genomic data, predation experiments and phenotypic measurements of larvae and their host plants, we show that novel white alleles in *Neodiprion lecontei* (pine sawfly) larvae spread via selection rather than drift. The cost of being rare was not offset by an enhanced aposematic display or immune function. Instead, bottom-up selection via host plants may drive divergence among populations as white larvae were disproportionately abundant on a pine species with a reduced carotenoid content relative to other pine hosts.

## INTRODUCTION

Determining how novel traits originate and become abundant is a core goal of evolutionary biology. To this end, vibrant colors that animals use to communicate their unprofitability to predators have made aposematic species attractive model systems (1, 2). Aposematism is a defensive strategy based on the shared costs of predator education: predators learn to associate conspicuous coloration with the unprofitability of the prey and avoid attacking prey individuals with a similar appearance in future encounters (3). As a result, the fitness of a particular aposematic color morph is expected to be dependent on its local abundance and evolve under positive frequency-dependent selection (2, 4). This frequency-dependence should reduce genetic and phenotypic variation in signal design because rare or novel color patterns suffer increased predation risk (5–7). However, intraspecific variation in warning color is surprisingly common both within populations (polymorphism) and among populations (polytypism) of aposematic species (2). Such variation offers valuable opportunities for characterizing the evolutionary and ecological processes that generate novel warning color traits and signal divergence (8).

Three scenarios could explain the origin and spread of novel warning color alleles that produce color differences among aposematic populations. First, under some circumstances, novel color alleles may evolve neutrally. For example, when predators focus only on a specific element or combination of elements in an aposematic signal, variation in other signal elements may have little to no impact on predation rates (9, 10). Under this scenario, among-population differentiation at color loci evolves under migration-drift balance. Second, even when novel color alleles evolve under negative (purifying) selection, stochastic shifts in color allele frequencies that occur when a small number of individuals colonize a new area or survive a population bottleneck could enable novel color alleles to reach threshold frequencies at which they are common enough to be protected (11–14). For both of these first two scenarios (neutrality and purifying selection), novel color alleles increase in frequency via genetic drift. The main difference between these scenarios is in the magnitude of drift needed to explain differentiation at color loci: the stronger the selection against a novel color allele, the stronger the drift (i.e., larger reduction in *N_e_*) required to enable the allele to spread.

A third explanation for polytypic warning color is that novel warning color alleles are favored in some populations. Because a multitude of selection pressures can act on color alleles, the net selection coefficient for a novel color allele can be positive, even when the color morph it produces experiences an elevated predation risk. For example, warning signal efficacy may trade off with enhanced thermoregulation (15, 16) or improved defense against pathogens (17). Less efficient warning signal forms can also be favored under certain dietary conditions (18–20) or maintained via genetic correlations with other traits through physical linkage or pleiotropy (21). Importantly, these three scenarios (neutrality, purifying selection, and positive selection) are not mutually exclusive and the origin and spread of novel color alleles could be facilitated by a combination of genetic drift under relaxed selection, followed by positive selection (12–14).

Ultimately, determining the relative contribution of drift and selection and the relative importance of different selection pressures to the evolution of warning color requires integrating genetic, demographic, and ecological analyses. Despite several promising study systems (e.g. (22–24)), this level of integration remains rare with one notable exception: Müllerian mimics in the genus *Heliconius* (e.g. (25, 26)). These iconic butterflies have contributed substantially to our understanding of warning color evolution based on endogenously produced pigmentation. However, the extent to which lessons learned from *Heliconius* apply to other aposematic species remains unclear.

For example, key factors such as the life stage that expresses warning signals (larval vs. adult) (27, 28), could have profound—and possibly predictable—impacts on color evolution that are not typically considered in current models for warning color evolution. Specifically, adult coloration of many aposematic taxa is often subject to positive frequency-dependent selection via not only predator learning, but also positive assortative mating by color (29–32). Color-based mating preferences in the adults can impact warning coloration evolution in two ways. First, when reproductive adults mate assortatively by color and color varies among populations, gene flow will be reduced and genome-wide differentiation can accumulate more readily between populations via genetic drift and divergent natural selection. Second, positive assortative mating by warning color may make it even more difficult for shifts in coloration to occur because rare color morphs will not only experience increased predation risk, but also reduced mating success (30, 33). As a result, this can further strengthen selection against novel warning signal forms (30, 32). By contrast, because coloration tends to be decoupled across ontogeny (28, 34, 35) but see (21), aposematic larvae are not usually subject to direct sexual selection. Thus, all else equal, we hypothesize that the cost of being a rare aposematic color allele may be lessened in the larval stage, making the evolutionary shift to a novel color morph comparatively easier for larvae. Currently we have very little data on how novel warning signals evolve and become abundant in immature stages (27), which prevents comparisons with the mechanisms found to be important in the adult stage.

Similarly, the source of color pigments - whether they are produced endogenously or acquired from the diet - can influence the relative importance of top-down and bottom-up selective agents acting on warning color variation (36–38). Bottom-up selection may have an especially strong impact in herbivorous species, for which variation in host-plant use across space can lead to differences in access to defensive compounds acquired from the host (39), the visual background against which coloration is displayed (40), and access to diet-derived pigments (41). While host shifts may also impact the availability of nutrients needed to produce pigments endogenously, warning signals based on endogenous pigments seem to be more robust and less variable under nutritional stress (19). As a consequence, spatial variation in diet quality and nutrient availability is hypothesized to have a much stronger impact on diet-derived warning signal pigmentation than on endogenously produced pigments (36). Yet, few studies have evaluated the contribution of diet quality to geographic variation in warning color pigmentation derived from the diet.

As a starting point to test these hypotheses and to address key gaps in currently available study systems, we investigated diet-based warning color evolution in redheaded pine sawfly (*Neodiprion lecontei*) larvae. *N. lecontei* are specialist herbivores of pines and occur over a wide geographical and climatic range in eastern North America (Fig 1). They are semi-social hymenopterans, meaning that larvae feed in large groups until the final instar, at which point they disperse from the feeding site to spin cocoons and pupate (42). Based on their bright coloration, *N. lecontei* larvae are assumed to be aposematic. They defend against predators and parasitoids collectively using a synchronized display in which they raise their heads and regurgitate resinous droplets of sticky fluid sequestered from the host plant (43–45).

**Figure 1.**
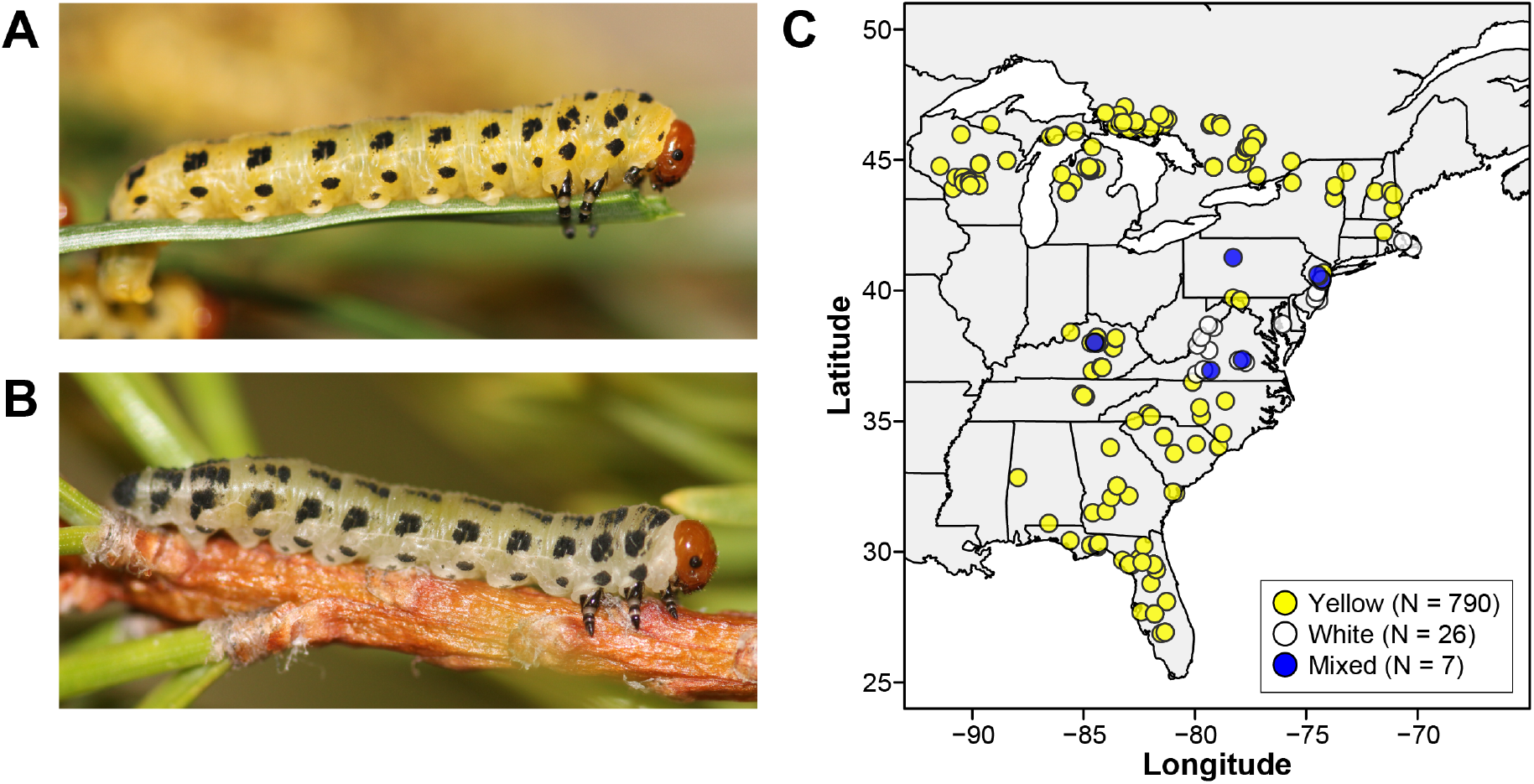
Appearance and distribution of white and yellow color morphs of *N. lecontei* larvae. Photographs depict representative yellow-bodied (**A**) and white-bodied (**B**) larvae (both photos by R. Bagley). (**C**) Approximate collecting locations and recorded color of 823 *N. lecontei* larval colonies collected between 2001-2004 and 2009-2016.

Throughout most of *N. lecontei*’s range, larvae have a bright, carotenoid-based yellow body color overlaid with several rows of melanic black spots (Fig. 1) (46). Like most insects, *N. lecontei* cannot synthesize carotenoids, which must therefore be obtained from the host plant. In addition, carotenoids are also thought to be involved in protecting the insect’s own tissues from defensive toxins (scavenging free-radicals) and in supporting immune defense (36, 47–50). Thus, diet quality impacts *N. lecontei* fitness in multiple ways, including the availability of defensive terpene compounds (39) as well as acting as a direct source of carotenoids for conspicuous warning coloration (46) and immune function. By contrast, *N. lecontei* adults are not aposematic, and adult and larval pigmentation is decoupled in pine sawflies (51). Therefore, larval color is not likely to be subject to sexual selection.

Although most *N. lecontei* populations have yellow larvae, field surveys have revealed the presence of white-bodied larvae in the Mid-Atlantic/northeastern United States (Fig. 1). Previous demographic and genetic mapping analyses provide insight into the origin of this larval color variation. First, a range-wide demographic analysis indicates that the white-body phenotype is the derived state for *N. lecontei* larval pigmentation (52). Second, a QTL mapping analysis of *N. lecontei* larval color suggests that the loss of yellow pigmentation in white-bodied populations is attributable to small number of large-effect loci that reduce or halt the transport of carotenoids from the gut to the integument (46). Although these studies have shed light on the origin of white-bodied larvae, additional data are needed to explain how novel white-bodied alleles spread in some *N. lecontei* populations.

To explain spatial variation in diet-based larval warning color, we combined population genomic analyses with multiple experiments to address five questions: (1) did a bottleneck facilitate a shift from yellow to white pigmentation, (2) is among-population differentiation at color loci attributable to drift or selection, (3) does the loss of yellow pigmentation impact warning signal efficacy, (4) does the loss of yellow coloration impact other defensive traits (immune and chemical defense), and (5) does larval color correlate with host-plant characteristics? Together, the answers to these questions enabled us to make inferences regarding the relative importance of drift and selection, as well as the relative importance of top-down and bottom-up selection pressures. When contrasted with study systems involving aposematic adults that use endogenous warning pigments, our results support the hypothesis that life stage and pigment source can have predictable impacts on warning color evolution.

## RESULTS

### 1. A reduction in population size did not facilitate the spread of novel white-body alleles

If novel color alleles are disfavored due to frequency-dependent predation, a novel deleterious allele could only spread if the effective population size (*N_e_*) is small enough for drift to overwhelm selection (specifically, when *s* < 1/*N_e_*). If a reduction in *N_e_* (e.g., during a bottleneck) facilitated the spread of white-body alleles in some *N. lecontei* populations, we should see genomic evidence of (1) historical isolation between white-bodied and yellow-bodied populations and (2) if there was isolation, a historical bottleneck in the white-bodied populations. To evaluate these predictions, we sampled 65 *N. lecontei* larvae from 29 locations throughout the United States. This sample consisted of 19 white-bodied larvae, 2 larvae from mixed-color colonies, and 44 yellow-bodied larvae (Table S1, Supplementary information). Our geographic sampling strategy was informed by a previous range-wide analysis that identified three main genetic and geographic clusters of *N. lecontei* populations (52)). These clusters—which were dubbed North, Central, and South—most likely formed as a consequence of isolation in different pine refugia during the Pleistocene (52). Because white-bodied populations are restricted to the Central cluster, we focused on this region only to avoid confounding color-associated divergence with refugia-associated divergence. Also, because larvae feed in colonies that typically consist of siblings, each larva came from a different colony.

To generate genome-wide single-nucleotide polymorphism (i.e., SNP) data for the *N. lecontei* samples, we performed double-digest restriction associated DNA (ddRAD) sequencing (53). We aligned reads (1.84 ± 1.14 million reads per individual) to a high-coverage, linkage-group anchored *N. lecontei* genome assembly (version 1.1; GenBank assembly Accession no. GCA_001263575.2; (46, 54)). After alignment, paralog filtering, and removal of putative PCR duplicates, 0.97 ± 0.54 million alignments remained. We then used STACKS (v1.46; Catchen *et al*. 2013) to construct RAD loci from these filtered alignments. After filtering on depth (>10x), missingness (≤30%), and minor allele frequency (>0.01), we obtained 17,444 ± 4,090 RAD loci (average coverage: 44.05 ± 22.37x) containing 70,297 SNPs. To minimize linkage disequilibrium between markers for population structuring analyses, we randomly sampled one SNP per locus. Finally, to remove paralogously mapping loci, we excluded markers displaying significant heterozygote excess (*P* < 0.01). After filtering, the number of SNPs for each of our two population structure datasets (one SNP per locus) was: 11,603 (all individuals) and 11,431 (“East” individuals only, see below). For our F_ST_ outlier analysis, we created a third dataset of 38,852 SNPs (same filters as our population structure datasets but excluding all unambiguously “West” individuals and retaining all SNPs for each RAD locus).

Using our genome-wide SNP data, we evaluated the relationship between color and population structure in three ways. The first two analyses included a model-based clustering analysis (admixture v1.23; (56) and a non-model based clustering analysis (discriminant analysis of principal components (DAPC) implemented in the adegenet R package (v1.3- 9.2;(57)). Both approaches recovered *K* = 2 as the optimal number of clusters (Fig. S1). Further investigation of assignment patterns under *K* = 2 revealed that this structure corresponded to geographical regions, not color. Specifically, individuals west of the Appalachian Mountains (“West”; i.e., those in Kentucky and Tennessee) belonged to one cluster and individuals east of the Appalachian Mountains assigned to the other (“East”) (Fig. 2a). Individual assignments were stable across all 100 admixture runs, and similar for most individuals between the admixture and DAPC methods (Fig. 2b).

**Figure 2.**
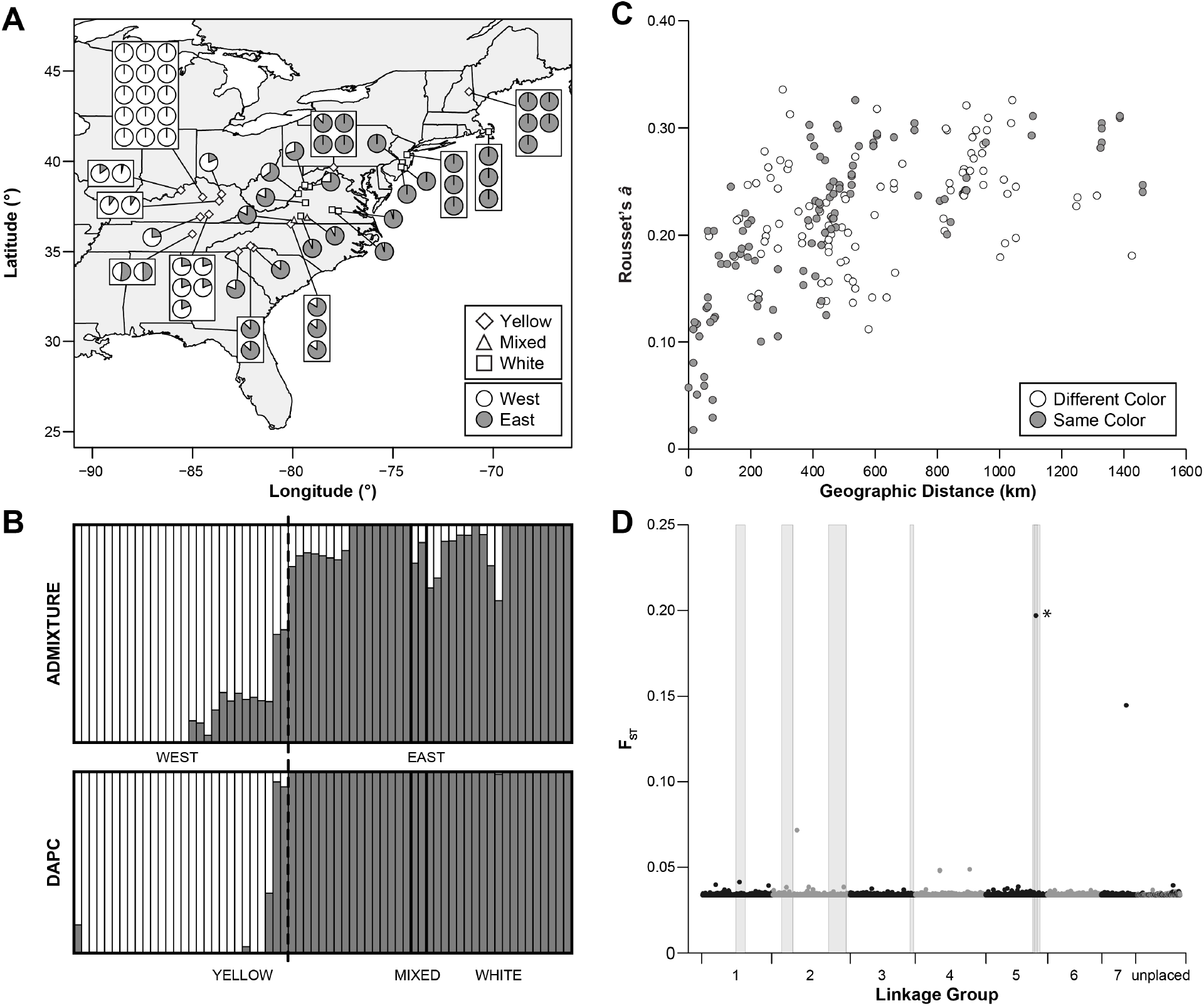
Genome-wide markers reveal no evidence of color-associated population structure in *Neodiprion lecontei*. **(A)** Color, location, and population assignment for 65 individuals included in the population structure analyses. The color of the collected colony is indicated by the shape overlaid on the collection location. At each site, each pie chart depicts the proportion of a single larva’s ancestry stemming from eastern (grey) and western (white) genetic clusters, both of which belong to the “Central” genetic cluster (Bagley et al. 2017). Ancestry proportions were inferred from admixture analysis under a model of *K* = 2. **(B)** Inferred ancestry for each individual sorted by color (Yellow, Mixed, or White) under a model of *K* = 2 based on the program admixture (top) and *adegenet* (bottom). Within each color, individuals are sorted in geographical order. While the western genetic cluster contains only yellow larvae, the eastern genetic cluster contains yellow, white, and mixed-color colonies. **(C)** Relationship between geographical distance (km) and genetic distance (as measured by Rousett’s a) for pairs of individuals sampled from the eastern genetic cluster. Filled circles indicate the pair had the same body color. **(D)** Per-site differentiation (F_ST_) between white-bodied and yellow-bodied individuals of *N. lecontei* across seven linkage groups and remaining unplaced scaffolds. Shaded grey boxes denote the locations of six QTL associated with larval body color in a cross between white-bodied and yellow-bodied populations (Linnen et al. 2018). The asterisk indicates the only significant F_ST_ outlier, which falls within two major-effect QTL intervals.

Our third population structure analysis asked whether color had a subtler impact on population structure that might have been missed in the first two clustering approaches. Specifically, we used a partial Mantel test (58–60) to ask whether different-colored colonies were more genetically dissimilar than same-colored colonies (i.e., “isolation-by-color”), after controlling for geographic distance (isolation-by-distance). To avoid pseudo-replication, we included only one randomly selected individual per site. For this analysis, we also excluded samples from mixed-color colonies and, on the basis of results from the first two population structure analyses, samples collected west of the Appalachian Mountains. Consistent with previous isolation-by-distance analyses (52), we found a significant correlation between genetic and geographical distance (*r* = 0.561; *p* = 0.0001). However, there was no correlation between body color and genetic differentiation when controlling for geographical distance (*r* = 0.023, *p* = 0.733; Fig. 2c). Overall, there was no evidence of any form of population structure corresponding to larval body color. Thus, there is no indication that white-bodied alleles rose to a high frequency in an isolated, bottlenecked population.

### 2. Geographic variation in color was consistent with selection, not drift

If body-color alleles evolve neutrally, among-population differentiation at color loci should reflect a balance between genetic drift and migration. Under this scenario, differentiation at color loci should be similar to neutral differentiation across the rest of the genome. Alternatively, if natural selection maintains geographic variation in larval color in the face of gene flow, differentiation at color loci should exceed genome-wide levels of differentiation. To determine whether any SNPs exhibited elevated differentiation between white-bodied and yellow-bodied samples, we conducted an F_ST_ outlier analysis with BayeScan v2.1 (61). This analysis identified a single SNP on chromosome 5 (position 21793198) with significantly elevated differentiation between white-bodied and yellow-bodied larvae (F_ST_ = 0.197, q-value = 0.012; Fig. 2d). Notably, this SNP fell within the intervals of two QTL peaks previously identified for larval body color in *N. lecontei*, including one QTL that explains ~52% of the difference in color between white- and yellow-bodied populations (46). This finding suggests that differences in pigmentation among *N. lecontei* populations are currently maintained by selection in the face of gene flow.

### 3. White and yellow larvae have equally effective aposematic displays

We next evaluated the role of predation as a source of selection shaping color variation in *N. lecontei*. If *N. lecontei* larvae are aposematic (as has been assumed (44)), they should be highly conspicuous to potential predators when viewed against a natural background (pine branches). Therefore, we first tested how conspicuous *N. lecontei* larvae are against their natural pine backgrounds in terms of their color and luminance. We quantified the conspicuousness of white- and yellow-bodied larvae against the three most common pine hosts for Central *N. lecontei*: *Pinus virginiana* (VA pine)*, P. echinata* (shortleaf pine), and *P. rigida* (pitch pine) by using mathematical models that simulate the vision of a insectivorous avian predator (blue tit, *Cyanistes caeruleus*) (62, 63). We found that regardless of the larval body color, body region, host species, and host tissue type, *N. lecontei* larvae were highly conspicuous against a pine background in terms of color contrast (JND (‘just noticeable differences’) values ranged between 10 and 50; Fig. 3a). Most larval/host combinations were also highly conspicuous in terms of luminance, but some individual larvae had JND <5 for some host background types (Fig. 3b). Nevertheless, these results support the hypothesis that *N. lecontei* larvae are conspicuous to their predators.

**Figure 3.**
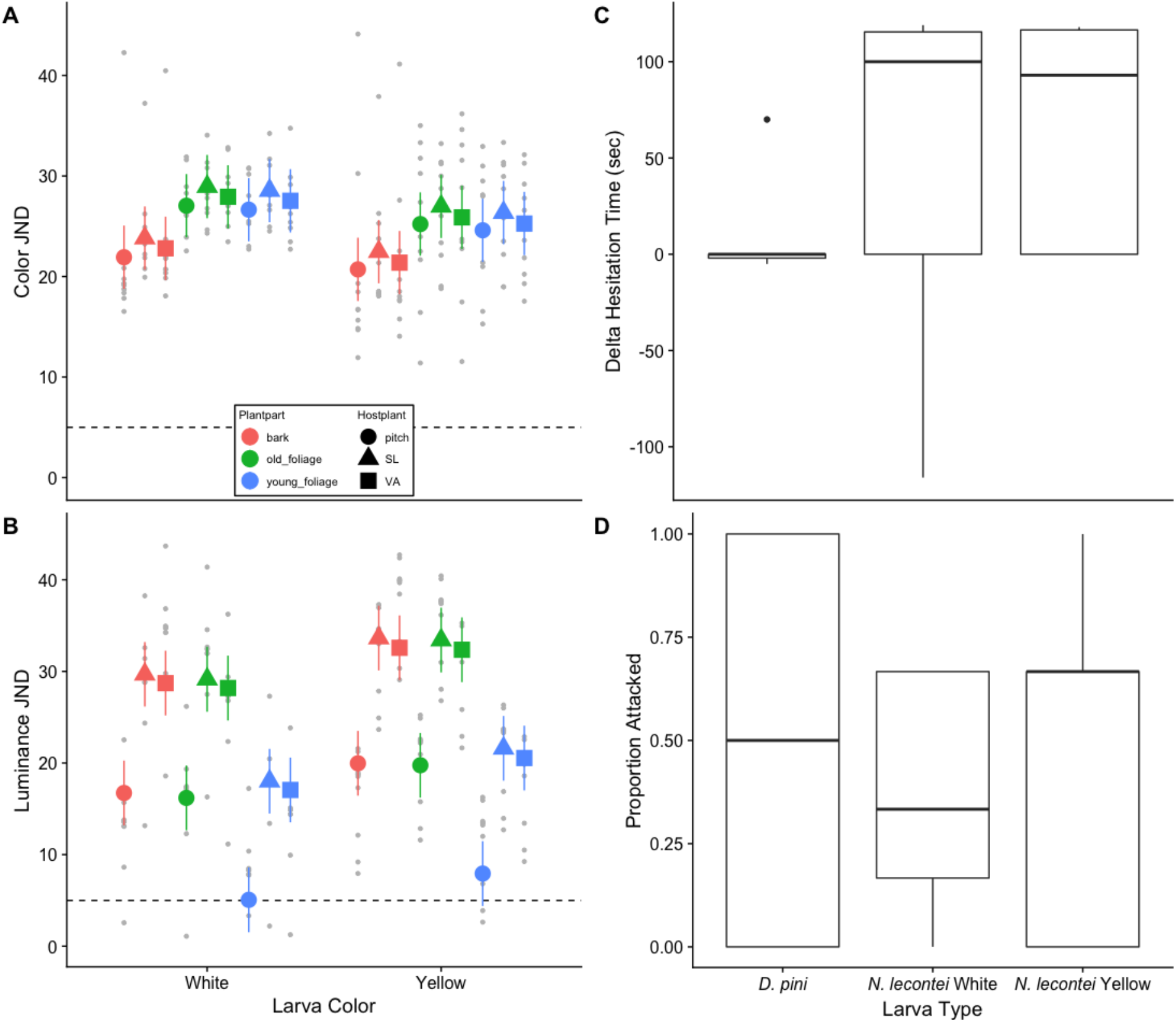
Warning signal efficacy did not differ depending on pigmentation in *N. lecontei*. Color **(A)** and luminance **(B)** contrasts against different host plants and host plant parts. For reference, JND values for prey/background combinations that are <1 are indistinguishable, values between <1 and 3 are hard to distinguish unless under optimal conditions, and values > 5 are easy to tell apart under most conditions (62). Dashed lines show the threshold value for JND=5 above which objects should appear clearly conspicuous for blue tits (*Cyanistes caeruleus*). Changes in attack latency **(C)** and attack risk **(D)** between the first and last trial in avoidance learning assays.

We next used our JND estimates to ask whether the conspicuousness of yellow and white larvae differed. For example, if white larvae are more conspicuous than yellow larvae, this could have facilitated the spread of white alleles due to increased signal efficacy (64, 65). However, we found no evidence to suggest that white and yellow larvae differ in conspicuousness, nor did we find any evidence that shifts in host plant would affect the conspicuousness of white and yellow larvae differently. First, white and yellow-bodied larvae did not differ significantly in terms of color or luminance (Table 1, Fig. 3a-b, Table S2 in Supplementary information). Similarly, white and yellow larvae appeared equally conspicuous against different host plants and plant parts (Table 1). Overall, conspicuousness of both larval color types varied significantly depending on the host plant species in terms of luminance, but not color (Table 1). In addition, conspicuousness of larvae varied depending on the plant part for both color and luminance (Table 1, Fig. 3a-b).

**Table 1.** ANOVA-table for color (A) and luminance (B) contrasts with respect to larval color, host plant, and plant part. Wald tests were performed on the model output object of a linear mixed model fit by REML.

|  | numDF | denDF | df | F | P |
| --- | --- | --- | --- | --- | --- |
| **A: Color contrast** |  |  |  |  |  |
| Intercept | 1 | 152 | 1, 152 | 698.8717 | <.0001 |
| Larval color | 1 | 18 | 1, 18 | 0.9021 | 0.3548 |
| Host plant | 2 | 152 | 2, 152 | 2.9706 | 0.0543 |
| Plant part | 2 | 152 | 2, 152 | 24.5574 | <0.0001 |
| Larval color * Host plant | 2 | 152 | 2, 152 | 0.0103 | 0.9897 |
| Larval color * Plant part | 2 | 152 | 2, 152 | 0.1658 | 0.8474 |
| **B: Luminance contrast** |  |  |  |  |  |
| Intercept | 1 | 152 | 1, 152 | 373.6643 | <0.0001 |
| Larval color | 1 | 18 | 1, 18 | 2.4059 | 0.1383 |
| Host plant | 2 | 152 | 2, 152 | 316.6230 | <0.001 |
| Plant part | 2 | 152 | 2, 152 | 261.5544 | <0.001 |
| Larval color *<br>Host plant | 2 | 152 | 2, 152 | 0.2123 | 0.8090 |
| Larval color *<br>Plant part | 2 | 152 | 2, 152 | 0.1724 | 0.8418 |

If *N. lecontei* larvae are aposematic, they should also be unpalatable to potential predators. To evaluate this prediction, we offered *N. lecontei* larvae to free-living house sparrows (*Passer domesticus*) in their natural habitat. These trials revealed that *N. lecontei* larvae are highly unprofitable prey to sparrows: whereas all of the mealworms offered were always attacked, only 2 / 20 (10 %) of the white *N. lecontei* larvae and 1/15 (6%) of the yellow *N. lecontei* larvae were attacked. Additionally, the similarity in attack rates between white and yellow larvae suggests that they were equally unpalatable.

A final piece of evidence needed to demonstrate aposematism is that the combination of conspicuousness and unpalatability facilitates predator avoidance learning. In support of the hypothesis that *N. lecontei* larvae are aposematic, we found that captive great tits learned to avoid brightly colored white and yellow *N. lecontei* larvae more effectively in comparison to light green and chemically defended *D. pini* sawfly larvae. Specifically, the change in the mean attack latency from the first to third trial was significantly larger for *N. lecontei* larvae than for *D. pini* larvae (Table 2, Fig. 3c). Hunger level of birds also significantly affected attack latencies; hungrier birds had shorter attack latencies (Table 2). However, changes in attack latencies between the first and third trials of white and yellow *N. lecontei* morphs did not differ significantly (Table 2, Fig. 3c). Attack rates (whether the great tit attacked or not) did not differ significantly between *D. pini* and *N. lecontei* larvae or between white and yellow *N. lecontei* larvae (Table 2, Fig. 3d). Again, hungrier birds were more likely to attack the prey (Table 2). Thus, our experiments suggest that there are neither costs nor benefits to being a white-bodied larva in terms of baseline signal efficacy.

**Table 2.** Predator response with respect to larval type. Output tables from (A) a linear model (y = change in attack latency, gaussian errors, t statistic) and (B) generalized linear mixed model (y = attack risk, binomial family, z statistic, random effect = larval ID) using planned contrasts (*D. pini* vs. pooled *N. lecontei*; white vs. yellow *N. lecontei*).

| **A: Change in attack latency** | **Estimate** | **Std. Error** | **T or z** | **P** |
| --- | --- | --- | --- | --- |
| Intercept | 11.428 | 22.778 | 0.502 | 0.6200 |
| *D. pini* vs. *N. lecontei* | 56.175 | 23.972 | 2.348 | 0.0265 |
| White *N. lecontei* vs. Yellow *N. lecontei* | -4.356 | 25.668 | -0.170 | 0.8665 |
| Hunger (mg mealworms eaten) | -14.043 | 33.213 | -0.423 | 0.6758 |
| **B: Attack risk** |  |  |  |  |
| Intercept | -1.3814 | 0.6693 | -2.064 | 0.03904 |
| *D. pini* vs. *N. lecontei* | -0.5644 | 0.6766 | -0.834 | 0.40414 |
| White *N. lecontei* vs. Yellow *N. lecontei* | -0.1464 | 0.6888 | -0.213 | 0.83165 |
| Hunger (mg mealworms eaten) | 3.3283 | 1.0703 | 3.110 | 0.00187 |

### 4. The loss of yellow coloration did not enhance other defensive traits

Because dietary carotenoids serve several important functions in insects, there may be trade-offs between the use of carotenoids for warning coloration and allocation to essential functions such as immunity and protection of an insect’s own tissues from the toxic compounds used in chemical defense (36, 47–49). If so, the spread of rare white-body alleles may have been facilitated by enhanced immune function and/or chemical defense as a higher proportion of carotenoids could be allocated to these non-color functions. This hypothesis predicts that there will be genetic correlations between larval body color and other carotenoid-related traits. To test this prediction, we evaluated trait correlations in recombinant F2 progeny derived from a cross between white-bodied and yellow-bodied *N. lecontei* (as in (46)).

Overall, our analysis of over 200 recombinant haploid males revealed that there was no correlation between body color (carotenoid content) and any of the defensive traits (behavioral, chemical, immune) measured. The volume of defense fluid was not significantly correlated with carotenoid pigmentation (S1B) (Table 3). Similarly, variation in the concentration of monoterpene compounds or total amount of terpene compounds was not explained by carotenoid pigmentation (Table 3). Carotenoid pigmentation did not differ significantly between defending (82.2%) and non-defending (16.7%) individuals. Encapsulation response and carotenoid pigmentation were also non-significantly correlated (Table 3). Together, these results do not support the hypothesis that the loss of yellow pigmentation was favored by natural selection because it enabled white-bodied larvae to allocate more resources to defense and immune functions (but see “Discussion”).

**Table 3.** Correlation between carotenoid concentration (S1B) and defensive behavior, quality and quantity of chemical defense against predators, and encapsulation response (defense against parasitoids). t statistic for Gaussian error LMMs (B & C & E) and z statistic for GLMMs (A & D).

|  | Estimate ± s.e. | t or z | P |
| --- | --- | --- | --- |
| A: Defensive fluid volume |  |  |  |
| Intercept | 0.603 ± 0.251 | 2.40 | 0.0163 |
| Body length | -0.072 ± 0.094 | -0.76 | 0.445 |
| S1B | 1.506 ± 1.930 | 0.78 | 0.435 |
| B: Monoterpene (per µL of fluid) |  |  |  |
| Intercept | 0.094 ± 0.013 | 7.08 | <0.0001 |
| S1B | 0.154 ± 0.1.6 | 1.46 | 0.153 |
| C: Total terpene (per µL of fluid) |  |  |  |
| Intercept | 0.118 ± 0.017 | 7.14 | <0.0001 |
| S1B | 0.177 ± 0.130 | 1.36 | 0.180 |
| D: Prob defense (deploys fluid or not) |  |  |  |
| Intercept | 1.94 ± 0.45 | 4.34 | <0.0001 |
| S1B | 3.99 ± 4.62 | 0.864 | 0.388 |
| E: Encapsulation response |  |  |  |
| Intercept | 48.14 ± 2.47 | 19.47 | <0.0001 |
| S1B | -13.39 ± 36.42 | -0.368 | 0.714 |

### 5. White-bodied larvae are associated with low-carotenoid hosts

Because the yellow body color of *N. lecontei* larvae is derived from carotenoids acquired from the host plants (46), a reduction in carotenoid availability in the larval diet (adults are non-feeding) could potentially favor the loss of yellow body color in some populations. To evaluate this possibility, we first asked whether the prevalence of white-bodied colonies differed among host plant species. To do so, we compiled larval colony color and host records from 12 years of collecting data, during which we obtained thousands of *N. lecontei* larva (Table S4).

These records confirmed that all-white colonies are restricted to the east of the Appalachians in the mid-Atlantic states and coastal regions of New England in the U.S. (Fig. 1c). Although some states had records of both all-white and all-yellow colonies, these were never collected at the same site. Mixed-color colonies were rare and 4/7 were collected on non-native pines in disturbed areas (e.g. parking lots). Overall, these results reveal that much of the variation in larval color is distributed among rather than within populations.

We then narrowed our focus to colonies that were collected on the three primary pine hosts in the Central region (*P. echinata*, shortleaf pine [N = 63]; *P. rigida,* pitch pine [N = 50]; and *P. virginiana*, Virginia pine [N = 167]). We found that the proportion of colonies that were white-bodied differed significantly across host plants (FET P < 1 x 10^−9^), with white-bodied colonies significantly more common on *P. rigida* than on either *P. echinata* (*P* = 9.6 x 10^−6^) or *P. virginiana* (*P* = 2.4 x 10^−9^) (Fig. 4a). Prevalence of white-bodied colonies did not differ between *P. virginiana* and *P. echinata* (*P* = 0.66).

**Figure 4.**
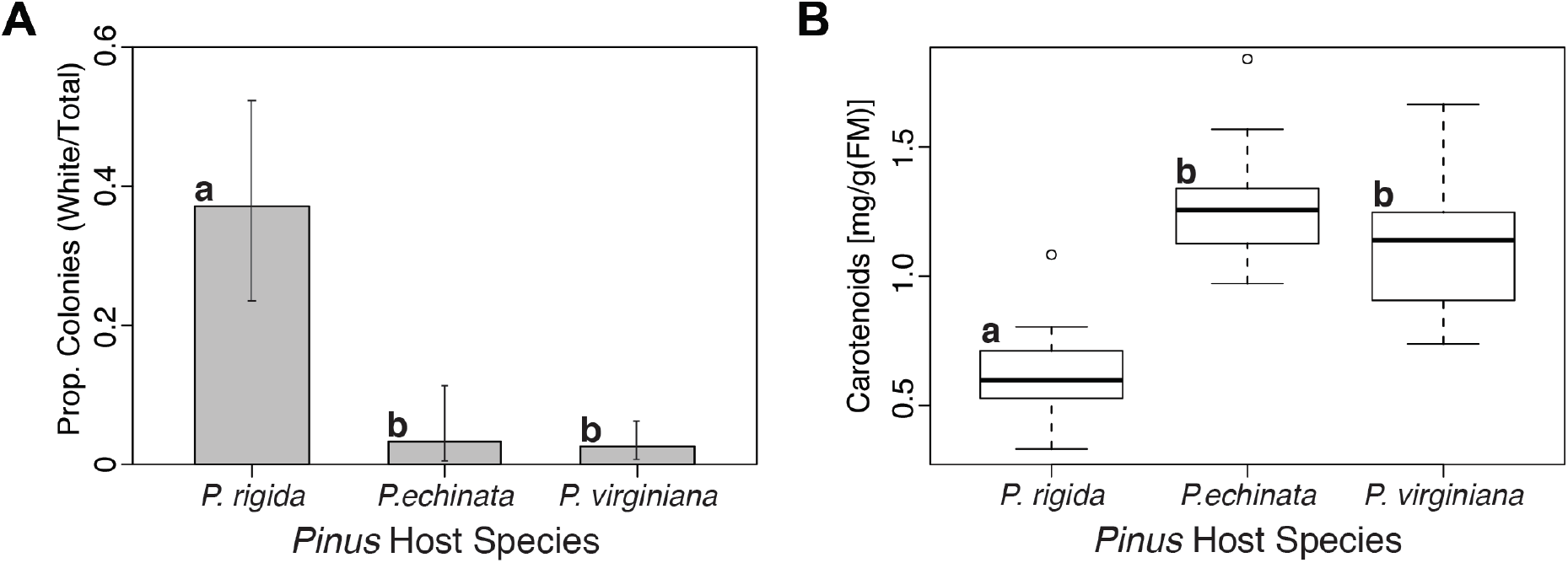
Relationship between host use and larval body color in *N. lecontei*. **(A)** Compared to *Pinus echinata* and *P. virginiana, P. rigida* had a significantly higher proportion of white-bodied larval colonies. Error bars are Clopper-Pearson 95% confidence intervals. **(B)** Compared to *P. echinata* and *P. virginiana, P. rigida* foliage had significantly lower carotenoid content. In all panels, letters indicate significant pairwise differences between colony color or host plants at *P* < 0.05.

Because the prevalence of white-bodied colonies differed among the three pine hosts, we next asked whether the carotenoid content of these hosts differed. In particular, if low carotenoid content in the host favored the loss of carotenoid-based body pigmentation, we would expect the lowest carotenoid content in the host species with the highest prevalence of white-bodied colonies. To test this prediction, we quantified carotenoid content for each host species following (66). We found that the three hosts differed significantly in carotenoid content (F2 = 20.12, P = 0.0022), with *P. rigida* having the lowest carotenoid content of all three hosts (Fig. 4b; *P. rigida* vs. *P. echinata:* z = 6.108, P = 3.02 x 10^−9^; *P. rigida* vs. *P. virginiana:* z = 4.538, P =1.13 x 10^−5^; *P. echinata* vs. *P. virginiana:* z = −1.570, P = 0.116). Notably, while *P. rigida* is widely distributed throughout the eastern United States, it is most abundant in the Atlantic Coastal plain (67, 68), where white-bodied colonies are predominantly found (Fig. 1c). Overall, these results support the hypothesis that an increased tendency to use a low-carotenoid host could have favored the spread of novel white-body alleles in the mid-Atlantic United States.

## DISCUSSION

Here, we investigated the evolutionary processes and selective pressures shaping geographic variation in body color in conspicuously colored *N. lecontei* larvae. We first confirmed that what we consider to be conspicuous coloration does, in fact, confer protection against visual predators of *N. lecontei* larvae (Fig. 3a-d). Therefore, we can expect *N. lecontei* larval coloration to evolve under positive-frequency dependent selection that disfavors novel color morphs (e.g. (7, 69)). Indeed, as expected under purifying selection, local populations of *N. lecontei* larvae tend to be monomorphic for color (Fig. 1c). In contrast to many studies on aposematic adult coloration (29–31), a lack of any form of color-associated population structure (Fig. 2) suggests that the spread of white alleles was not facilitated by genetic drift in isolated populations. Instead, our finding that a single color locus exceeded genome-wide levels of genetic differentiation between white- and yellow-bodied larvae (Fig. 2) suggests that geographical differences in color are attributable to divergent natural selection rather than migration-drift balance. Together, these results suggest that for aposematic larvae, genetic drift need not be invoked to explain shifts in warning color.

Having determined that white-body alleles likely evolved under positive selection, we next examined sources of selection that could have potentially offset (or exacerbated) the cost of being rare. Perhaps unexpectedly, we did not find any evidence that the loss of yellow pigmentation impacts defense against natural enemies. Body color had no impact on the efficacy of the warning signal (Fig. 3). We also found no evidence that white-body alleles have indirect effects on chemical defense or immune function via pleiotropy or physical linkage (Table 3). Instead, we found that white-bodied larvae were disproportionately abundant on a pine species with low carotenoid content (Fig. 4). Together these findings suggest that bottom-up selection via host-plant quality may play an important role in driving warning color polytypism in immature stages that rely on diet-derived pigmentation. Overall, these results are consistent with our hypothesis that life stage and pigment source have predictable impacts on the evolutionary processes and selective pressures shaping warning color variation. Here, we contrast our results with other aposematic study systems and highlight priorities for future work on this and other aposematic systems.

The lack of color-associated population structure in *N. lecontei* contrasts with several other aposematic study systems that implicate geographic isolation and genetic drift as key facilitators of polytypic warning signals. For example, in *Heliconius* butterflies, genetic drift is thought to promote the establishment of novel wing pattern variants in accordance with Wright’s shifting balance model (11, 14, 70) (but see (13, 71, 72)). In support of this hypothesis, multiple *Heliconius* species have pronounced population genetic structure indicative of subdivision and drift (73). Color-associated population structure has also been detected in other aposematic species, such as the strawberry poison frog, *Dendrobates pumilio* (74) and the polytypic dyeing dart poison frog (*Dendrobates tinctorius*) (22). One important difference between *N. lecontei* and many other aposematic study systems is the role of warning coloration in mate choice. While larval color variation in *Neodiprion* sawflies is unlikely to have any impact on mate preferences in the non-aposematic adults, positive assortative mating based on the color of aposematic adults has been demonstrated experimentally in both *Heliconius* (30, 33, 75) and *D. pumilio* (32). The absence of this added barrier to evolving novel color morph may explain, in part, why a shift in *N. lecontei* larval coloration did not require a drastic decrease in effective population size.

In addition to a lack of sexual selection, several other features of larval defensive displays in *N. lecontei* (and other taxa) have the potential to reduce the strength of purifying selection against novel warning color alleles, thereby facilitating their spread in the absence of isolation and genetic drift. First, *N. lecontei* larvae present their defensive chemicals externally (39, 44), a strategy that has been shown to reduce attack rates against novel conspicuous prey as predators can estimate a prey individual’s defensive capacity before tasting it (76). Second, both yellow and white *N. lecontei* larvae have melanin-based black spots across their body, which predators could use as an additional signal of unprofitability together with the bright pigmentation (but see (77). This similarity in one aspect of the aposematic pattern could have decreased initial costs of rarity for a white larval form, thereby facilitating the shift from yellow to white pigmentation (6). Third, and perhaps most importantly, the costs of being a low frequency color morph could be mitigated by gregariousness in chemically defended prey. *N. lecontei* larvae feed in large aggregations, which have been shown to increase avoidance learning efficacy and initial wariness of predators towards both conspicuous and cryptic unprofitable prey (78–80). Gregariousness could have therefore facilitated shifts in warning coloration among *N. lecontei* populations. Finally, we cannot exclude the possibility that predators could also generalize avoidance learning across white to yellow color variants (22), but see (10). Therefore, additional experiments are needed to determine the interactive effects of predator’s generalization, larval group size, and color. To assess if predation risk for yellow and white-bodied larvae varies among *N. lecontei* populations, future research should also test for differences in the survival of white and yellow-bodied larvae across their geographical range and across different visual backgrounds (81, 82).

In terms of possible benefits that could have facilitated the spread of novel white-body alleles via positive selection, our results suggest that geographic variation in the abundance of a low-carotenoid host (pitch pine) may have favored the loss of carotenoid-based larval coloration in some *N. lecontei* populations. Thus, rather than strong top-down selection by predators and pathogens, bottom-up selection via host-plant characteristics may have driven shifts in warning color among *N. lecontei* populations. This finding contrasts with strong top-down selection pressures that may explain endogenously produced polymorphic and polytypic warning color in *Arctia plantaginis* (82) and polytypic warning color in *Heliconius melpomene*. Historical demographic analyses suggest that *H. melpomene* radiated into areas already occupied by another, more abundant aposematic species, *H. erato* (e.g. (12, 13, 73, 83)). Whereas warning color polytypisms were thought to have evolved via drift in *H. erato*, later radiating *H. melpomene* are thought to have evolved under positive selection to match locally abundant *H. erato* color phenotypes (73, 83). Consistent with this “advergence” hypothesis, early-radiating *erato*-clade species exhibit much more pronounced population substructure than *melpomene*-clade species (73). Like *N. lecontei, H. melpomene* exhibits geographic variation in host use (84, 85). But unlike *N. lecontei, Heliconius* butterflies primarily rely on endogenous pigments for warning coloration (reviewed in (86)). Although these differences are intriguing, comparative analysis of color evolution in diverse taxa is needed to determine whether pigment source reliably predicts primary selective agents shaping pigment variation.

In the *N. lecontei* system, additional experimental work is also needed to clarify causal links between differences in host use and differences in fitness between *N. lecontei* color morphs. Our observation that *P. rigida*, a low-carotenoid host, harbors disproportionately more white-bodied larvae (Fig. 4) is consistent with the hypothesis that white alleles were favored due to carotenoid limitation. However, pine species differ in other characteristics besides carotenoid content that could potentially favor different carotenoid allocation strategies in the larvae. For example, differences in pine species color result in differences in larval luminance contrast values (Table 1, Fig. 3). Nevertheless, larvae are conspicuous on all three hosts and in the absence of a host-by-larvae interaction, host color seems unlikely to drive color divergence among larval populations*. P. rigida* could also differ from the other pine species in nutritional profile, secondary compounds (terpene content), and/or abiotic environments in ways that increase the demand for antioxidant functions of carotenoid compounds (87). However, based on previous studies, *N. lecontei* seem relatively insensitive to variation in the host-plant terpene content (39).

If carotenoid limitation is the primary selection pressure favoring white body color, there are two non-mutually exclusive mechanisms through which host carotenoid content could differentially impact the fitness of yellow and white *N. lecontei* larvae. First, production of conspicuous yellow coloration could be constrained under a carotenoid-limited diet, potentially weakening the signal efficacy (saturation and brightness) of yellow genotypes that develop on a low-carotenoid host. Second, since carotenoids are also involved in physiological processes such as immune defense (36, 47–50), trade-offs between warning signaling and these other functions could result in divergent selection on pigmentation on different host plants. These trade-offs and constraints should be most evident when the availability of pigments or their precursors are limited in the diet (19). However, such trade-offs should also generate genetic correlations between levels of carotenoid-based pigmentation and other fitness traits (especially immune or antioxidant function), which we did not find in this study (Table 3, see also (88)). That said, an important limitation of our data is that we did not evaluate conspicuousness or genetic correlations (trade-offs) in larvae that developed on the low carotenoid host, *P. rigida*. To rigorously test the signaling efficacy and carotenoid trade-off hypotheses, experimental manipulations of dietary carotenoid content are needed.

To conclude, our integrated analysis of color variation in *Neodiprion* sawflies provides a valuable point of comparison for other classic aposematic systems, such as *Heliconius* butterflies. While both taxa exhibit striking among-population variation in color attributable to major-effect loci, our data indicate that the observed polytypisms likely evolved under very different evolutionary scenarios and selection pressures. We hypothesize that these differences ultimately stem from differences in the source of pigments and the role of color in mate choice. To evaluate these hypotheses more rigorously, comparable analyses of diverse aposematic taxa that vary in pigment sources (dietary vs. endogenous), life stages (immature vs. adult), and behavior (mobility and gregariousness in larval stage vs. adult stage) are needed (27, 35, 46, 89). More generally, this kind of integrative study approach will give us a more comprehensive understanding of how organisms adapt to changes in their environment (90) and how ecological selection during immature life-stages impacts adaptation and speciation.

## MATERIALS AND METHODS

### DNA Extraction and Library Preparation

To extract DNA and generate ddRAD sequencing libraries we followed protocols outlined in (52), but used a new set of sequencing adapters and PCR primers. Briefly, we digested DNA with the restriction enzymes *EcoRI* and *NlaIII*. For adapter ligation, we assigned each sample one of 48 unique 5-10bp variable length barcodes (91) (Table S5). Following pooling and size selection (379 +/- 76bp) on a PippinPrep (SageScience, Beverly, MA, USA), we amplified libraries using high-fidelity polymerase (Phusion; NEB, Ipswich, MA) with primers that incorporated an Illumina multiplexing index (Table S6), as well as a string of four degenerate bases for PCR duplicate detection. We then sequenced these libraries with 150-bp reads on two replicate lanes of an Illumina HiSeq 4000 housed at the University of Illinois Roy J. Carver Biotechnology Center.

### Genotyping

Our sequencing yielded a total of 1.86 ± 1.15 (SD; standard deviation) million single-ended reads per individual. To quality-filter and demultiplex raw sequencing reads, we used the default settings of the *process_radtags* module in stacks (v1.46; (55)). We then aligned the resulting reads (1.84 ± 1.14 million reads per individual) to a high-coverage, linkage-group anchored *N. lecontei* genome assembly (version 1.1; GenBank assembly Accession no. GCA_001263575.2; (46, 54) using the “very sensitive” end-to-end alignment mode in bowtie2 (v2.3.1; (92). We then used SAMTOOLS (v1.3; (93) to retain only uniquely-mapping reads with MAPQ scores ≥30. Putative PCR duplicates were identified based on the sequence of the 4 degenerate bases in the index read (provided as a second fastq file) and removed using a custom python script (defRemove_ddRAD_PCRduplicates.py, Supplementary files). We then constructed RAD loci from the filtered alignments in stacks’ *ref_map.pl* pipeline (v1.46; Catchen *et al*. 2013). To ensure high-confidence genotype calls (53, 94), we required a minimum stack depth of 10 (*-m* 10). We then used stacks’ *populations* module to call SNPs present in ≥70% of individuals (*-r 0.7*) and with a minimum minor allele frequency of 0.01 (*--min_maf 0.01*). After alignment, paralog filtering, and removal of putative PCR duplicates, 0.97 ± 0.54 million alignments remained, which were assembled into 17,444 ± 4,090 RAD loci (average coverage: 44.05 ± 22.37x) containing 70,297 SNPs. For population structuring analyses (see below), we also randomly sampled one SNP per locus (*--write_random_snp*) to minimize linkage disequilibrium between markers. Finally, as an additional step to remove paralogously mapping loci, we performed exact tests of Hardy-Weinberg equilibrium in vcftools (v0.1.15; (95)) and excluded markers displaying significant heterozygote excess (*P* < 0.01).

### Inference of Population Structure

We evaluated the relationship between color and population structure in three ways. First, we used the model-based clustering approach implemented in the program admixture (v1.23; (56) to assign the proportion of ancestry for each individual from *K* ancestral populations without prior designation. We performed 100 independent runs for *K* = 1 through *K* = 10. The optimal *K* was selected by comparing 5-fold cross-validation (CV) error values for each value of *K* as suggested in the admixture manual. We then used the main pipeline of clumpak (v1.1; (96)) to summarize and evaluate stability of assignment solutions across the 100 replicates of each *K*.

Second, we used a non-model-based discriminant analysis of principal components (DAPC) implemented using the *dapc* function in the adegenet R package (v1.3-9.2;(57)), a multivariate approach that transforms genotypes using principal components analysis (PCA) to maximize differentiation between and minimize variation within groups (97). We identified the optimal number of clusters from *K* = 1 through *K* = 10 using a *K*-means clustering algorithm. To avoid overfitting of discriminant functions, following α-score optimization (Fig. S3), we performed this analysis using 1 principal component (PC) that explained ~7% of total variation (Fig. S4). The clustering solutions were then compared using Bayesian Information Criterion (BIC), following (97).

Third, to evaluate whether color has a subtler impact on population structure, we asked whether different-colored colonies were more genetically dissimilar than same-colored colonies (i.e., “isolation-by-color”), after controlling for geographic distance (isolation-by-distance). To do so, we used a partial Mantel test (58–60). To avoid pseudo-replication, we included only one randomly selected individual per site. We also excluded samples from mixed-color colonies and, on the basis of population structure results (Table S7), samples collected west of the Appalachian Mountains (in TN and KY). For remaining individuals, we generated pairwise distance matrices for genetic distance, geographical distance, and “color” distance. For genetic differentiation, we estimated Rousett’s a, an individual measure analogous to the F_ST_/(1-F_ST_) ratio ((98), using spagedi (v1.5a; (99)). We also calculated pairwise linear geographical distance between individuals using spagedi. For the “color” distance matrix, we coded pairs with the same body color (i.e., both individuals are yellow or both individuals are white) as having a distance of 0, while pairs with different body colors (one white individual and one yellow individual) were coded as a distance of 1. We assessed “isolation by color” using a partial Mantel test that examined the correlation between genetic distance and color distance, while controlling for the impact of geographical distance. We performed this partial Mantel test using passage2 (v2.0.11.6; (100)), and significance was determined using 10,000 permutations.

To determine whether any SNPs were significantly differentiated between white-bodied and yellow-bodied samples, we conducted an F_ST_ outlier analysis with BayeScan v2.1 (61). We converted the original VCF file to GESTE/BayeScan format using PGDSpider (v2.1.1.5; (101)). Our pilot analysis consisted of 20 runs with 5,000 iterations each; and our Markov chain was run for a total of 100,000 iterations, with a burnin of 50,000 generations, a thinning interval of 10, and a prior odds ratio of 100.

### *Conspicuousness of* N. lecontei *larvae against different host plants*

We quantified the conspicuousness of white- and yellow-bodied larvae against the three most common pine hosts for Central *N. lecontei: Pinus virginiana* (VA pine)*, P. echinata* (shortleaf pine), and *P. rigida* (pitch pine). Larvae used for these analyses were derived from a laboratory colony that was established from a single mixed-color colony collected on an introduced host, *Pinus sylvestris*, in Piscataway, New Jersey (40° 32′58.4″N, 74° 25′50.9″W) in August 2013. For host material, we collected five clippings from each of three individual trees. All clippings were collected in July 2018 from the University of Kentucky Arboretum in Lexington, KY.

To quantify color of *N. lecontei* larvae and their host plants, we recorded reflectance spectra from each larval and host sample with a USB2000 spectrophotometer (Ocean Optics, Largo, FL). For larvae, we recorded nine reflectance spectra across the dorsal, lateral, and ventral surfaces (three spectra per surface) of 20 individuals (10 white and 10 yellow) chosen at random from our laboratory population. For host plants, we recorded 15 spectra across the new foliage (current year’s growth), old foliage (previous years’ growth), and bark (5 spectra per region) from each of 45 clippings (15 fresh clippings from each *Pinus* species). After binning and trimming the raw spectra to 1-nm intervals between 300 and 750 nm, we averaged each bin for each sample to obtain a single summary spectrum for the dorsolateral (dorsal + lateral) and ventral surfaces of each larva and for the foliage and bark portions of each plant clipping. These averages were used in vision model analyses.

To predict whether a typical avian predator (blue tit, *Cyanistes caeruleus*) could discriminate between *N. lecontei* larvae and their natural pine backgrounds in terms of their color and luminance, we used a discrimination threshold model that assumes that noise in the receptors limits discrimination ability (62, 63). The model utilizes information about the visual system, such as the sensitivity and relative abundance of different receptor types, and estimates of noise that arise in the photoreceptors. An average spectrum per each stimulus type was modeled for a blue tit’s photon catch values for the single and double cones with a standard D65 irradiance spectrum. Color vision in birds stems from the 4 single cone types, whereas luminance-based tasks apparently stem from the double cones (Osorio and Vorobyev 2005). For the color model, we therefore used the 4 single cones, whereas the luminance model was based on the double cones (Siddiqi et al. 2004). We used a Weber fraction of 0.05 for the discrimination model for the most abundant cone type and the relative proportion of cone types in the blue tit retina (long wave = 1.00, medium wave = 0.99, short wave = 0.71 and ultraviolet (UV) sensitive = 0.37).

Using the same approach as in (102), we calculated “just noticeable difference” (JND) values for every combination of larval body region (dorsolateral and ventral) and host background (foliage and bark for *P. virginiana*, *P. echinata*, and *P. rigida*) for each larva. For reference, JND values for prey/background combinations that are <1 are indistinguishable, values between <1 and 3 are hard to distinguish unless under optimal conditions, and values > 5 are easy to tell apart under most conditions (62).

We also used our JND estimates to ask whether the conspicuousness of yellow and white larvae differed. To evaluate this possibility, we used linear mixed models (‘lme’ function of the ‘nlme’ package in R; (103)), with JND contrast as the dependent variable (color and luminosity values were analyzed separately). The fixed effect structure consisted of the additive effects of larval coloration (yellow or white), host plant (VA pine, shortleaf pine, or pitch pine), plant part (bark, old foliage or young foliage), and the two-way interactions between larval coloration and each of the other two predictors.

### Unpalatability of white and yellow larvae to avian predators (nest box experiments)

We offered *N. lecontei* larvae to free-living house sparrows (*Passer domesticus*) from a population that has been continuously monitored since 1992 at the University of Kentucky’s Agricultural Experimental Research Station in Lexington, KY (38°06′16.3″N 84°29′13.3″W) (104). Larvae for this experiment were sampled from the same mixed-color laboratory stock used in the conspicuousness analyses. After sorting by color (white or yellow), we froze live larvae at −80°C until needed. We also used spectrophotometric measurements to verify that these human-sorted larvae reflected true underlying differences in color (Table S3) and that color differences between frozen larvae recapitulated differences between living larvae (Table S3).

For the predation experiments, we installed feeding platforms (described in (105)) adjacent to house sparrow (*Passer domesticus*) nest boxes located at the University of Kentucky’s Agricultural Experimental Research Station in Lexington, KY (38°06′16.3″N 84°29′13.3″W) (104). Upon each platform, we placed a plastic, 150-mm petri dish with a color image of *Pinus* foliage taped to the base (hereafter, feeding tray). Starting approximately one week prior to experimental trials, we trained birds to use the feeding trays by offering seeds and dead mealworms (killed by freezing) ad libitum. Once eggs had hatched and birds from a particular nest box consistently removed mealworms, we started the experiment for that nest box.

Prior to the start of each experiment, we thawed frozen sawfly larvae (see above) for 30 minutes and installed a Panasonic SDR-S70 camcorder into the recording box. Then, at the start of each experiment, we placed a mealworm on the petri dish to ensure that birds were motivated to forage. After the mealworm was consumed, we offered individual dead sawfly larvae on the feeding tray in five consecutive trials per nest box (sawfly larvae for a particular nest box were either all white or all yellow). Each trial lasted a maximum of 20 minutes if the sawfly larva was not attacked at all after a bird had detected it. Birds were observed from a vehicle at a distance of approximately 10 m with binoculars; later, observations were confirmed by examination of accompanying video recordings. For each trial, we recorded whether a bird attacked the prey (i.e., touched it with the beak) or did not attack the prey (i.e., clearly observed the prey, but did not touch it). To ensure birds were motivated to forage, we also offered mealworms at the end of each experiment.

In total, we attempted 14 experiments on different nest boxes, 7 of which failed because the nestlings died and/or the sparrows were not motivated to feed (i.e., did not remove the mealworm at both the start and end of the trial). We successfully tested 4 nest boxes (8 sparrows) for white larvae and 3 nest boxes (6 sparrows) for yellow larvae.

### Signal efficacy of white and yellow larvae (predator learning experiments)

The avoidance learning experiment was conducted in plywood cages (plywood 50 x 50 x 70 cm, w x d x h) at Konnevesi Research Station (described in e.g. (65)). Prey were offered through a hatch behind a visual barrier, which enabled us to record the exact time of prey detection because the bird had to go around the barrier or fly on top of it to see the prey. After birds were acclimated to the cage, *N. lecontei* larvae were offered one at a time and dorsal side up on a white dish (i.e. all the prey items were easy to detect for birds) in three consecutive trials similar to (65). At the beginning of each experiment, a bird’s motivation to feed was confirmed by offering them a mealworm. To measure learning rate, we used attack latency (the time from when the bird noticed the prey to when it touched/attacked the prey with its beak (65, 106)). We also recorded the total number of sawfly larvae attacked per bird over three trials. Because hunger can affect a predator’s readiness to attack defended prey (107), we also quantified the hunger level as the mass of mealworms eaten after the experimental trials.

11 great tits were tested with white *N. lecontei* larvae, 11 great tits with yellow *N. lecontei* larvae, and 10 great tits with light green *D. pini* larvae. *N. lecontei* larvae were obtained from the same mixed-color colony described above, and *D. pini* larvae were obtained from laboratory insect culture described in (108). All larvae were killed by freezing and allowed to thaw 30 minutes before the experiments with birds.

We used two measures of predator response: delta hesitation time to attack (trial 3 – trial 1 hesitation times) and probability of attack. For the first measure, we used a linear model (‘lm’ function from the base R package), since each bird had only one data point. Model diagnostics were performed on the residuals. For the second measure, we used a generalized linear mixed effects model, with bird identity as the random effect (since each had three trials) and binomial family (‘glmer’ function from the ‘lme4’ R package; (109)). For both models, we used planned contrasts so that the fixed effects test for a difference (i) between *D. pini* and (the average) *N. lecontei* and (ii) between white and yellow *N. lecontei*. Moreover, both models include a measure of hunger as a covariate (amount of meal worms eaten after [each/combining all] trial[s]). Results tables were obtained using the ‘summary’ function.

### Genetic correlations between color and defensive traits

We evaluated trait correlations in recombinant F2 progeny produced via crossing *N. lecontei* females from a laboratory line derived from a white-bodied population (Valley View, VA; 37°54’47”N, 79°53’46”W) to males from a laboratory line derived from a yellow-bodied population (Bitley, MI; 43°47’46”N, 85°44’24”W) (as in (46)). Because *Neodiprion* are haplodiploid, these crosses generated hybrid, diploid F_1_ females and non-hybrid haploid males. To generate haploid F2 males, we reared the offspring of virgin F_1_ females.

To estimate carotenoid content for each larva, we took five measurements with the spectrophotometer from each of three body regions: the dorsum, lateral side, and ventrum. We then used the program CLR: Colour Analysis Programs v1.05 (110) to process raw spectra and compute color saturation (S1B) values for each larva, which correlate negatively with carotenoid content in multiple taxa (111). To obtain a single S1B value per larva, we averaged across the 15 reflectance spectra. To evaluate larval chemical defense, we gently poked each larva with a capillary tube twice between the front legs on the ventral side and recorded presence/absence of a defensive regurgitant. If a defensive regurgitant was produced, we collected it in a 5-μL capillary tube. To estimate defense fluid quantity, we measured the length of the regurgitant in the capillary tube with digital calipers. To control for the effect of body size on the amount of defense fluid produced, we also measured larval body length. To evaluate defense fluid quality, we estimated the concentration of mono- and other terpene compounds in the samples using a gas chromatograph similarly to (108). Finally, to evaluate larval immune defense, we measured encapsulation response as described in (112). In total, we measured color (carotenoid content) and defensive traits (presence, quantity, and quality of defensive regurgitant; encapsulation response) as described in the methods in N = 212 male larvae from 10 F_1_ mothers.

Correlations between color (carotenoid content), chemical defense (defensive behavior, quantity and terpene concentration of defense fluid), and immune defense (encapsulation response) were evaluated in R (version 3.6.2 and 4.4.0; (R Core Team 2019)). We used generalized linear mixed models using the function ‘glmer’ from the package ‘lme4’ (version 1.1-23; (109)) and linear mixed models using the function ‘lme’ from the package ‘nlme’ (v. 3.1-147; (103)). For all models, colony identity was used as the random effect and the continuous predictors were centered, so that the intercept term estimates the outcome variable at the average predictor levels.

### Chemical defense of recombinant F2 males

Larval defense fluid samples in capillary tubes were placed in 1.5-mL microcentrifuge tubes containing 500 μl n-hexane and stored in a −20 °C freezer until analysis. To determine monoterpenes and other terpene compounds in the defensive regurgitates, a Shimadzu, GC-2010 Plus (Shimadzu Corp., Kyoto, Japan) gas chromatograph fitted with a flame ionization detector (FID) was used. Samples dissolved in hexane were injected (splitless injection of 1 μl sample, inlet temperature 290 °C) into a Zebron ZB-5MSi capillary column (length 30 m, inner diameter 0.25 mm, film thickness 0.25 μm; Phenomenex Inc., Torrance, CA, USA). Helium was used as the carrier gas at a flow rate of 1 ml min^−1^ (83.6 kPa). The temperature program started at 50 °C (1.5 min hold) and the column oven was heated with a rate of 10 °C min^−1^ to 180 °C, and then at a rate of 2.5 °C min^−1^ to 290 °C (10 min hold). The temperature of FID was 290 °C. We used L-fenchone as an internal standard. We identified the compounds with separate runs using a GC equipped with a mass spectrophotometer (Shimadzu 2010 GC/MS; Shimadzu Corp., Kyoto, Japan) and otherwise similar conditions as in the GC/FID analyses. For the identification of terpenes we compared the mass spectra (electron ionization 70 eV) of the obtained peaks to those of the library. We used the retention times of terpene peaks in GC/MS chromatograms to assign the compounds in GC/FID chromatograms. Masses (μg) of different compounds in the defense fluid samples were calculated by: dividing the area of a compound’s peak by area of the internal standard’s peak, and then multiplying it with the concentration of the internal standard (20 ppm). Concentration (%) of monoterpene compounds (total mass of all the monoterpene compounds/sample) and other terpene compounds (total mass of all the other terpene compounds/sample) in the defense fluid sample was calculated with the following formula: (HPLC-value in mg/defense fluid sample size (μl)) x 100.

Chemical concentration variables were modeled using total volume as an offset term and Gaussian errors. Encapsulation response was also modeled using Gaussian errors. To account for large skew and excess zeros in defense fluid quantity, this trait was transformed (0.00001 added to all data points) and modeled using Gamma family. As a binary outcome variable, the probability of defense was modeled using binomial family.

### Encapsulation response of recombinant F2 males

To evaluate immune defense, we measured encapsulation response as described in (112) after chemical defense measurements. Larvae were first anaesthetized with CO2 (108). A sterilized hypodermic needle was then used to puncture the skin on the dorsal part of the individual and a nylon implant (length 3 mm, diameter 0.11 mm) was inserted into the resulting hole. After 24 h, the implant was removed, dried, and photographed under a Zeiss DiscoveryV8 microscope equipped with an Axiocam 105 camera and ZEN lite 2012 software (Carl Zeiss Microscopy, LLC, Thornwood, NY). Every implant was photographed three times from three different angles (Rantala *et al*. 2000). Image visualization software ImageJ 1.47v (National Institutes of Health, USA) was then used to quantify the grey value of the implant, which was used as a measure of the encapsulation reaction. The grey value of the background was subtracted from the grey value of the implant in order to correct for potential variation in light source. The darker the implant, the stronger the immune response.

### Carotenoid content of host plants

To measure carotenoid content in *P. rigida*, *P. echinata*, and *P. virginiana*, we sampled ~10-20 needles from each of the 45 clippings (5 clippings from three individual trees per species) that were used to measure host reflectance spectra (see “*Conspicuousness of N. lecontei larvae against different host plants”*). We then chopped the needles into 2-5 mm segments and stored them in 1.7-mL microcentrifuge tubes in the −20°C freezer until pigment extraction. To extract pigments and quantify carotenoid content, we followed protocols outlined in (66). First, we placed 15.0 mg (±0.3mg) of chopped needle into a 2.0-mL tube, to which we added 1.5 mL of 200-proof ethanol. Next, samples were vortexed and stored in the dark at 65°C for 24 hours. After cooling to room temperature, samples were vortexed for approx. 1 minute at medium speed and centrifuged at 13500g (rcf) for 5 minutes. For each sample, we then transferred 200 μl of the supernatant to each of three wells (replicates, to reduce measurement error) in a clear, flat-bottomed 96-well plate. Using a Synergy HT plate reader (Biotek Instruments, Winooski, VT) and Gen5 software, we read the absorbance of each sample at 664 nm, 649nm, and 470nm. We then used these values and an equation for ethanol solvent (66, 114) to compute carotenoid concentration. We used the *lme4* (v1.1-21), *lmertest* (v3.1-1) and *multcomp* (v1.4-12) R (v3.6.2) packages to fit a linear mixed model to the data (with host species and individual tree as fixed and random effects, respectively) and since the host effect was significant, to compare pairs of hosts with post-hoc Tukey contrasts.

## Ethics statement

Predation experiments were conducted in accordance with Finnish, U.S., and institutional guidelines for the care and use of animals. U.S. experiments with house sparrows were conducted with the approval of the University of Kentucky (IACUC protocol # 2012-0948). The experiments with wild great tits at Konnevesi Research Station (Jyväskylä, Finland) were carried out with permission from the Central Finland Centre for Economic Development, Transport and Environment (KESELY/1017/07.01/2010) and license from the National Animal Experiment Board (ESAVI-2010-087517Ym-23).

## Data accessibility statement

When the paper will be accepted, all the data and R-codes used in these analyses will be included in open access database.

## Acknowledgements

We are grateful to: David Westneat (University of Kentucky) and Johanna Mappes (University of Jyväskylä) for offering facilities, equipment, and technical help for the avian experiments and bird maintenance; Jimi Kirvesoja and Hannu Pakkanen for the help with chemical analyses; Danielle Herrig for custom python scripts; and members of the Linnen lab for rearing and collecting sawflies (especially Kim Vertacnik and Adam Leonberger). This study was funded by the National Science Foundation (site-based Research Experience for Undergraduates summer program, DBI-1062890; DEB-1257739 to CRL; DEB-CAREER-1750946 to CRL), Academy of Finland via Centre of Excellence in Biological interactions and individual grant (#257581 to CL) for CL. SC was funded by an Academy of Finland Fellowship (#314219).

## Notes

### Competing Interest Statement

The authors have declared no competing interest.

