## Supplementary information for "The impact of life stage and pigment source on the evolution of novel warning signal traits"

### The impact of life stage and pigment source on warning color evolution: an integrative analysis of color variation in aposematic pine sawfly larvae

Carita Lindstedt, Robin Bagley, Sara Calhim, Mackenzie Jones, and Catherine Linnen

**Table S1. Sampling locations for all individuals included in population genomic analyses.** Specimens noted with a dagger (†) were adult females; all other specimens were larvae of unknown sex, but confirmed to be diploid via heterozygosity estimates. Full sequences for the adapter-ligated barcodes and Illumina indexes are listed in Tables S3 and S4, respectively.

| ID | Location | Latitude | Longitude | Host | Color | Index | Barcode |
|---|---|---|---|---|---|---|---|
| LL004_01† | Lexington, KY | 38.014 | -84.504 | *P. echinata* | Yellow | ATCACGAT | TGGCACAGA |
| LL005_01† | Lexington, KY | 38.014 | -84.504 | *P. rigida* | Yellow | TGACCAAT | TGGCAACAGA |
| LL010 | Bronston, KY | 36.9279 | -84.6193 | *P. virginiana* | Yellow | GTGAAACG | AACTGG |
| LL011 | London, KY | 37.07133 | -84.2114 | *P. echinata* | Yellow | AGTCAACA | CTAAGCA |
| LL013 | London, KY | 37.07133 | -84.2114 | *P. echinata* | Yellow | TGACCAAT | ACAACT |
| LL014 | London, KY | 37.07133 | -84.2114 | *P. echinata* | Yellow | CCGTCCCG | ACTGCGAT |
| LL015_01 | London, KY | 37.07133 | -84.2114 | *P. echinata* | Yellow | ATCACGAT | TCAGAGAT |
| LL042 | Ford, VA | 37.24904 | -77.7253 | *P. echinata* | White | TTAGGCAT | GCAAGCCAT |
| LL058_2 | Lexington, KY | 38.014 | -84.504 | *P. virginiana* | Yellow | AGTCAACA | ACGGTACT |
| LL062_1 | Lexington, KY | 38.014 | -84.504 | *P. echinata* | Yellow | CGATGTAT | TGACGCCA |
| LL064 | Lexington, KY | 38.014 | -84.504 | *P. rigida* | Yellow | TTAGGCAT | ACGGTACT |
| LL070_1 | Lexington, KY | 38.014 | -84.504 | *P. rigida* | Yellow | CGATGTAT | ATATCGCCA |
| LL073_2R | Lexington, KY | 38.014 | -84.504 | *P. echinata* | Yellow | CGATGTAT | AAGACGCT |
| LL074_5 | Lexington, KY | 38.014 | -84.504 | *P. echinata* | Yellow | ATCACGAT | ACAACCAACT |
| LL084 | West Yarmouth, MA | 41.64545 | -70.2302 | *P. rigida* | White | ATCACGAT | GAGCGACAT |
| LL084_02 | West Yarmouth, MA | 41.64545 | -70.2302 | *P. rigida* | White | CCGTCCCG | GAGCGACAT |
| LL084_03 | West Yarmouth, MA | 41.64545 | -70.2302 | *P. rigida* | White | GTGAAACG | CTCGCGG |
| LL102 | Lexington, KY | 38.014 | -84.504 | *P. rigida* | Yellow | TTAGGCAT | AACGCACATT |
| LL137 | Lexington, KY | 38.014 | -84.504 | *P. virginiana* | Yellow | CGATGTAT | CTCGCGG |
| LL142_02 | Crossville, TN | 35.98003 | -85.0152 | *P. virginiana* | Yellow | AGTCAACA | CAACCACACA |
| LL144_01 | Crossville, TN | 35.98003 | -85.0152 | *P. virginiana* | Yellow | GTCCGCAC | TCAGAGAT |
| LL180_02 | Goshen, KY | 38.40235 | -85.5859 | *P. echinata* | Yellow | TGACCAAT | TATGT |
| LL181 | Goshen, KY | 38.40235 | -85.5859 | *P. echinata* | Yellow | GTGAAACG | CACCA |
| LL194_02 | Stanton, KY | 37.80602 | -83.6779 | *P. virginiana* | Yellow | TGACCAAT | TGCTT |
| LL195_02 | Stanton, KY | 37.80602 | -83.6779 | *P. virginiana* | Yellow | GTCCGCAC | CTTGA |
| RB020Db | London, KY | 37.066 | -84.159 | *P. echinata* | Yellow | GTCCGCAC | ATTAT |

| | | | | | | | |
|---|---|---|---|---|---|---|---|
| RB028_01† | Egg Harbor, NJ | 39.69028 | -74.593 | *P. rigida* | White | GTCCGCAC | CAACCACACA |
| RB076_01 | Lexington, KY | 38.014 | -84.504 | *P. virginiana* | Yellow | GTGAAACG | TGACGCCA |
| RB107_01 | Mountain Grove, VA | 38.212 | -79.719 | *P. rigida* | White | GTGAAACG | CCGAACA |
| RB108_01 | Deer Run, WV | 38.678 | -79.399 | *P. rigida* | White | GTGAAACG | CGTCGCCACT |
| RB110.01† | Browns Mill, NJ | 39.934 | -74.533 | *P. rigida* | White | AGTCAACA | ACAACCAACT |
| RB112_01 | Tuckerton, NJ | 39.621 | -74.428 | *P. rigida* | White | TGACCAAT | CCTTGCCATT |
| RB118_01 | Brandywine, WV | 38.592 | -79.172 | *P. virginiana* | White | CGATGTAT | GCAAGCCAT |
| RB119_01 | Buena Vista, VA | 37.713 | -79.367 | *P. virginiana* | White | TGACCAAT | AACGCACATT |
| RB126_01 | Lexington, KY | 38.014 | -84.504 | *P. virginiana* | Yellow | GTGAAACG | AACGTGCCT |
| RB164 | Tyron, NC | 35.28019 | -82.1175 | *P. virginiana* | Yellow | AGTCAACA | TATTCGCAT |
| RB165 | Tyron, NC | 35.28014 | -82.1179 | *P. virginiana* | Yellow | TTAGGCAT | CTCGCGG |
| RB167_01 | Chesnee, SC | 35.18269 | -81.963 | *P. virginiana* | Yellow | ATCACGAT | GGCTTA |
| RB190 | Pickens, SC | 35.01364 | -82.7156 | *P. virginiana* | Yellow | CCGTCCCG | ACGGTACT |
| RB220 | Sandy Ridge, NC | 36.49728 | -80.1035 | *P. virginiana* | Yellow | AGTCAACA | GGTGCACATT |
| RB221 | Sandy Ridge, NC | 36.49728 | -80.1035 | *P. virginiana* | Yellow | GTCCGCAC | TCACTG |
| RB222 | Sandy Ridge, NC | 36.49728 | -80.1035 | *P. virginiana* | Yellow | ATCACGAT | GCGTCCT |
| RB223 | Bassett, VA | 36.80192 | -79.9371 | *P. echinata* | Mixed | ATCACGAT | ACGGTACT |
| RB224 | Penhook, VA | 36.97347 | -79.6076 | *P. rigida* | White | ATCACGAT | AAGACGCT |
| RB225 | Penhook, VA | 36.97347 | -79.6076 | *P. rigida* | White | AGTCAACA | GCAAGCCAT |
| RB226_old | Gretna, VA | 36.93961 | -79.2893 | *P. rigida* | Mixed | ATCACGAT | ACCAGGA |
| RB227 | Amelia, VA | 37.31906 | -78.0428 | *P. rigida* | White | AGTCAACA | CTCGCGG |
| RB228_new | Amelia, VA | 37.31906 | -78.0428 | *P. rigida* | White | ATCACGAT | CTCGCGG |
| RB229 | Amelia, VA | 37.31906 | -78.0428 | *P. rigida* | White | CCGTCCCG | ATGAGCAA |
| RB286 | Clear Springs, MD | 39.63111 | -77.9631 | *P. virginiana* | Yellow | GTCCGCAC | TAGCGGAT |
| RB287 | Clear Springs, MD | 39.63111 | -77.9631 | *P. virginiana* | Yellow | AGTCAACA | AACGCACATT |
| RB288_02 | Clear Springs, MD | 39.63111 | -77.9631 | *P. virginiana* | Yellow | TGACCAAT | GGTGT |
| RB289 | Clear Springs, MD | 39.63111 | -77.9631 | *P. virginiana* | Yellow | GTGAAACG | ATAGAT |
| RB290 | Old Bridge, NJ | 40.38742 | -74.3351 | *P. rigida* | White | TGACCAAT | CTTGA |
| RB304 | Ossippee, NH | 43.81939 | -71.2052 | *P. rigida* | Yellow | GTCCGCAC | ATGAGCAA |
| RB305 | Ossippee, NH | 43.81939 | -71.2052 | *P. rigida* | Yellow | AGTCAACA | CTCTCGCAT |
| RB306 | Ossippee, NH | 43.81939 | -71.2052 | *P. rigida* | Yellow | CGATGTAT | TATTCGCAT |
| RB307 | Ossippee, NH | 43.81939 | -71.2052 | *P. rigida* | Yellow | TGACCAAT | AACGTGCCT |

| | | | | | | | |
|---|---|---|---|---|---|---|---|
| RB308 | Ossippee, NH | 43.67597 | -71.0816 | *P. rigida* | Yellow | TTAGGCAT | GCCTACCT |
| RB339 | Lexington, KY | 38.014 | -84.504 | *P. echinata* | Yellow | GTCCGCAC | CCTTGCCATT |
| RB343_01 | Lexington, KY | 38.014 | -84.504 | *P. rigida* | Yellow | ATCACGAT | GGAACGA |
| RB355 | Lexington, KY | 38.014 | -84.504 | *P. virginiana* | Yellow | AGTCAACA | GGTGT |
| RB368 | Old Bridge, NJ | 40.36814 | -74.3022 | *P. rigida* | White | TGACCAAT | GGTGCACATT |
| RB369 | Old Bridge, NJ | 40.36814 | -74.3022 | *P. rigida* | White | TTAGGCAT | TGGCAACAGA |
| RB404 | Morehead, KY | 38.18613 | -83.5568 | *P. echinata* | Yellow | GTGAAACG | TCACTG |

**Figure S1. Cluster number determined by model and non-model based methods.** A *K* of 2 is suggested using both ADMIXTURE's 5-fold CV error score (A) and *adegenet*'s BIC score (B).

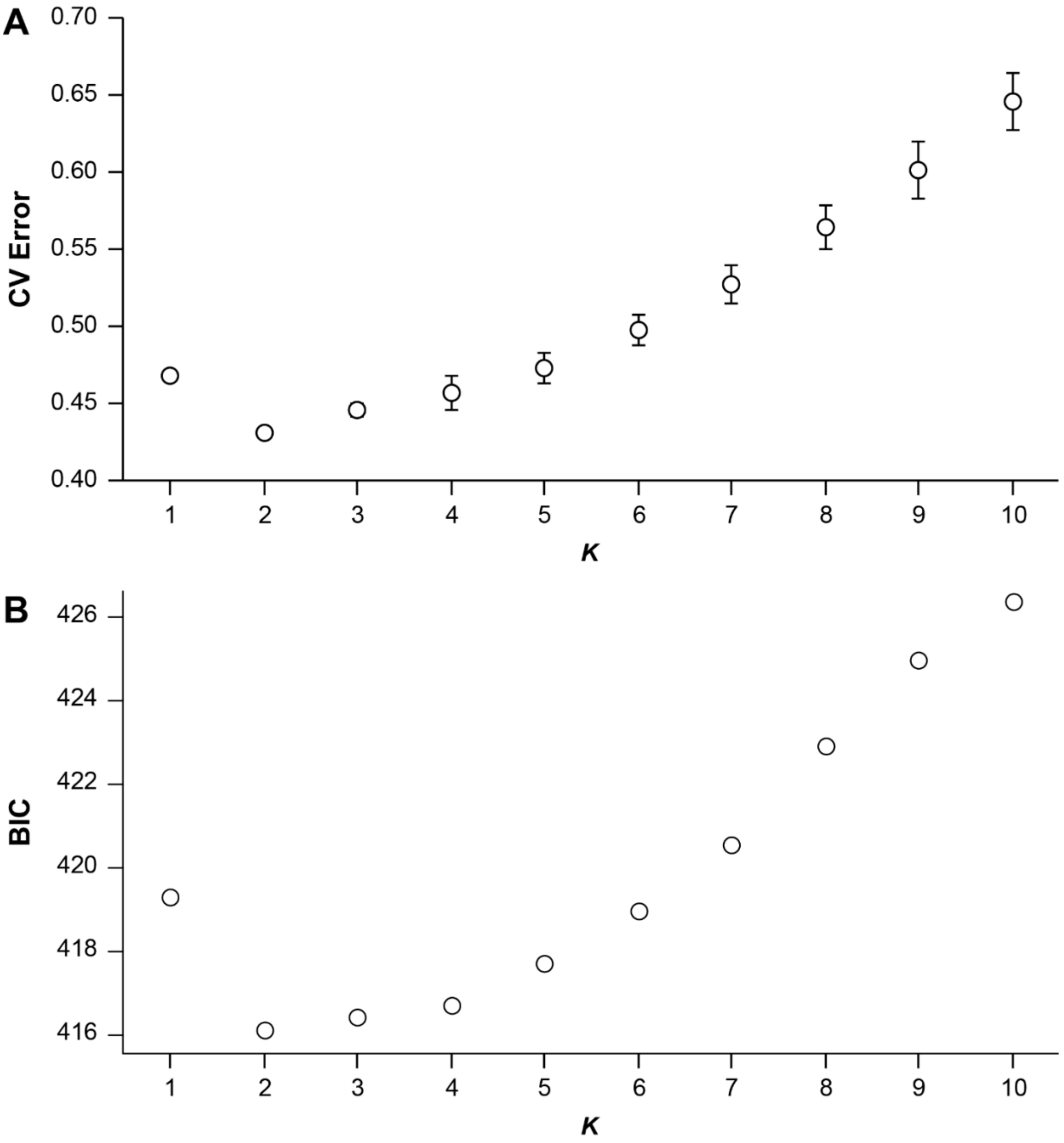

**Table S2. Random and fixed effects from the linear mixed model on *N. lecontei* and host plant (*P. virginiana* [VA], *P. echinata* [SL], and *P. rigida*) color traits.**

*Color contrasts*

Random effects

| **Source of variation** | **Name** | **σ²** | **SD** |
|---|---|---|---|
| Individual ID | Intercept | 4.038 | 4.135 |

Fixed effects

| **Source of variation** | **Estimate** | **s.e.** | **DF** | t | **P** |
|---|---|---|---|---|---|
| Intercept | 21.919 | 1.607 | 152 | 13.644 | <0.001* |
| Larval color | -1.210 | 2.272 | 18 | -0.533 | 0.601 |
| Host plant SL | 1.902 | 1.068 | 152 | 1.782 | 0.077 |
| Host plant VA | 0.884 | 1.068 | 152 | 0.828 | 0.409 |
| Plant part Old foliage | 5.122 | 1.068 | 152 | 4.797 | <0.001* |
| Plant part Young foliage | 4.728 | 1.068 | 152 | 4.428 | <0.001* |
| Larval color Y * Host plant SL | -0.139 | 1.510 | 152 | -0.092 | 0.927 |
| Larval color Y * Host plant VA | -0.214 | 1.510 | 152 | -0.142 | 0.888 |
| Larval color Y * Plant part Old foliage | -0.616 | 1.510 | 152 | -0.408 | 0.684 |
| Larval color Y * Plant part young foliage | -0.840 | 1.510 | 152 | -0.556 | 0.579 |

*Luminance contrasts*

Random effects

| **Source of variation** | **Name** | **σ²** | **SD** |
|---|---|---|---|
| Individual ID | Intercept | 5.167 | 3.227 |

Fixed effects

| **Source of variation** | **Estimate** | **s.e.** | **DF** | t | **P** |
|---|---|---|---|---|---|
| Intercept | 16.729 | 1.802 | 152 | 9.285 | <0.001* |
| Larval color | 3.240 | 2.548 | 18 | 1.271 | 0.220 |
| Host plant SL | 12.953 | 0.834 | 152 | 15.537 | <0.001* |
| Host plant VA | 11.994 | 0.833 | 152 | 14.396 | <0.001* |
| Plant part Old foliage | -0.540 | 0.837 | 152 | -0.645 | <0.520 |
| Plant part Young foliage | -11.665 | 0.827 | 152 | -14.101 | <0.001* |
| Larval color Y * Host plant SL | 0.713 | 1.179 | 152 | 0.605 | 0.546 |
| Larval color Y * Host plant VA | 0.612 | 1.178 | 152 | 0.519 | 0.605 |
| Larval color Y * Plant part Old foliage | 0.325 | 1.181 | 152 | 0.275 | 0.783 |
| Larval color Y * Plant part young foliage | -0.369 | 1.174 | 152 | -0.315 | 0.754 |

**Table S3. Human-sorted white and yellow *Neodiprion lecontei* larvae reflected underlying differences in color. Similarly color of frozen white and yellow *N. lecontei* larvae recapitulate differences between fresh larvae.** We recorded reflectance spectra from 10 frozen larvae of each color (white and yellow) and from 10 living, $CO_2$-immobilized larvae of each color. For each larva, we recorded nine spectra across the dorsal, lateral, and ventral surfaces (three spectra per surface) and computed an average S1B value (summary statistic that correlates with carotenoid content) as described in the main text. We then performed a two-way ANOVA in R with human-designated color category (white or yellow) and larval treatment (fresh or frozen) as main effects, plus an interaction term, followed by a Tukey's Honest Significant Difference test with FDR correction for multiple testing.  Significant adjusted p-values (<0.05) are bolded. A significant "Category" term indicates that there are significant differences in color between larvae that were sorted as yellow or white. Although freezing also has an impact on larval color (significant treatment effect), the larval color morphs do not respond differently to freezing (the interaction term is not significant). Also note that white and yellow larvae differ significantly in color (and in the same direction) regardless of whether both are fresh or both are frozen (Fresh yellow vs. Fresh white and Frozen yellow vs. Frozen white comparisons).

*Two-way ANOVA table*

| | DF | Sum Sq. | Mean Sq. | F-value | P-value |
|---|---|---|---|---|---|
| Category (White or Yellow) | 1 | 0.0241 | 0.0241 | 8.39 | **0.00637** |
| Treatment (Fresh or Frozen) | 1 | 0.109 | 0.109 | 38.0 | **4.16e-07** |
| Category:Treatment | 1 | 0.00452 | 0.00452 | 1.57 | 0.218 |
| Residuals | 36 | 0.103 | 0.00287 | | |

*Tukey HSD test*

| Comparison | Diff | Lwr | Upr | P. adj. |
|---|---|---|---|---|
| Frozen:White-Fresh:White | -0.0279 | -0.0924 | 0.0367 | 0.654 |
| Fresh:Yellow-Fresh:White | -0.0833 | -0.148 | -0.0187 | **0.00708** |
| Frozen:Yellow-Fresh:White | -0.154 | -0.218 | -0.0891 | **0.0000012** |
| Fresh:Yellow-Frozen:White | -0.0554 | -0.120 | 0.00914 | 0.114 |
| Frozen:Yellow-Frozen:White | -0.126 | -0.190 | -0.0612 | **0.0000405** |
| Frozen:Yellow-Fresh:Yellow | -0.0704 | -0.135 | -0.00580 | **0.0282** |

**Figure S2.** Avoidance learning rates of birds towards white and yellow *Neodiprion lecontei* larvae were compared with light green-black *Diprion pini* larvae with the similar defensive compounds (Photo Carita Lindstedt).

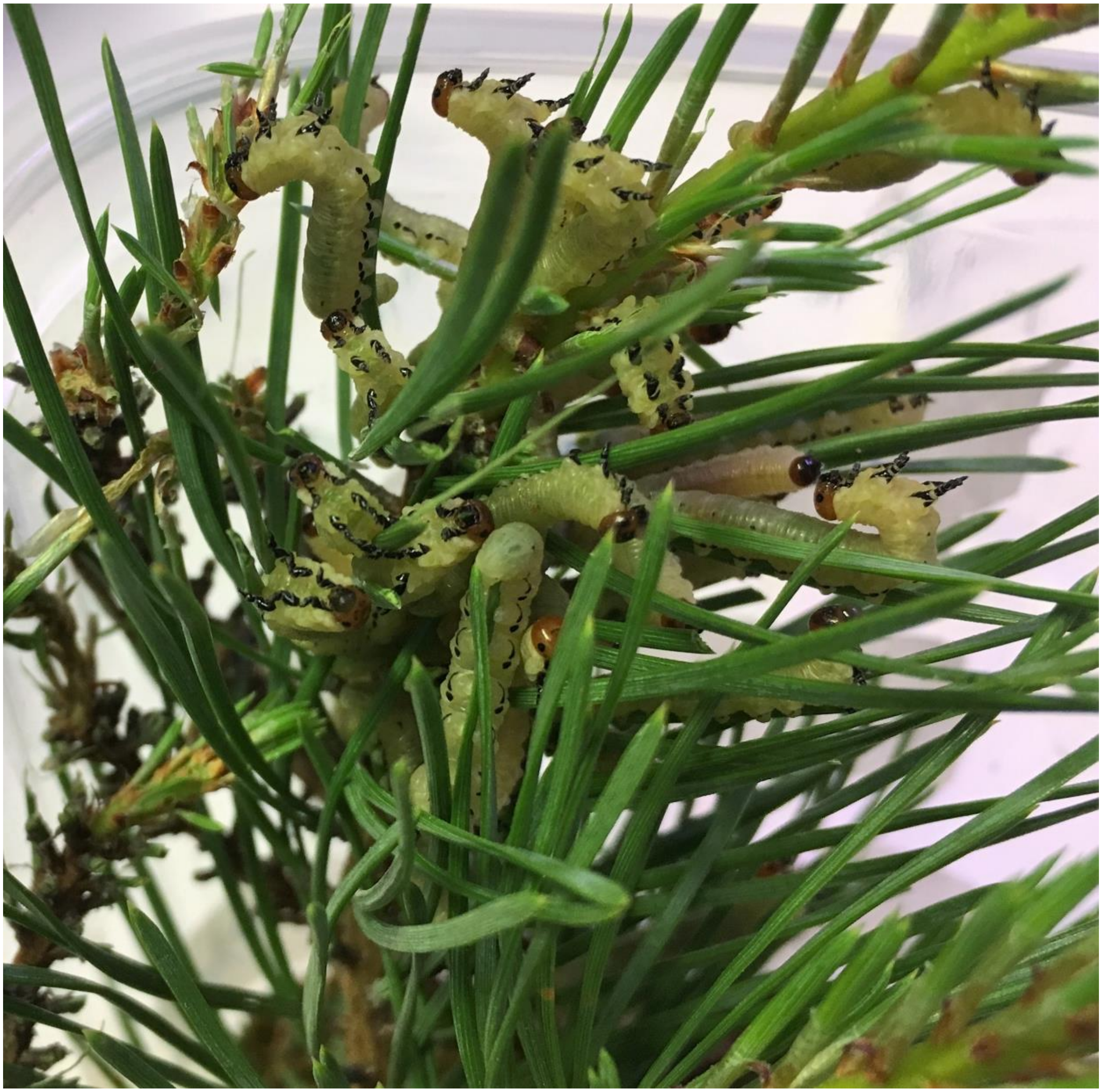

**Table S4. Collection location, collection host, collection date, body color, and genetic cluster for all *N. lecontei* collected between 2001 and 2016.**

| Specimen ID | Latitude | Longitude | Larval Color | Genetic Cluster | City, State/Province | Collection Host | Collection Date |
|---|---|---|---|---|---|---|---|
| 001-01 | 42.229 | -71.523 | YELLOW | CENTRAL | Hopkinton, MA | *P. banksiana* | 19-Jul-2001 |
| 002-01 | 42.229 | -71.523 | YELLOW | CENTRAL | Hopkinton, MA | *P. banksiana* | 19-Jul-2001 |
| 003-01 | 42.229 | -71.523 | YELLOW | CENTRAL | Hopkinton, MA | *P. banksiana* | 19-Jul-2001 |
| 017-01 | 44.544 | -73.215 | YELLOW | CENTRAL | Malletts Bay, VT | *P. resinosa* | 3-Aug-2001 |
| 018-01 | 44.544 | -73.215 | YELLOW | CENTRAL | Malletts Bay, VT | *P. resinosa* | 3-Aug-2001 |
| 025-0263 | 44.137 | -75.639 | YELLOW | NORTH | Kemptville, ON | *P. resinosa* | 20-Jul-2002 |
| 025-0309 | 44.395 | -77.205 | YELLOW | NORTH | Tweed, ON | *P. resinosa* | 7-Aug-2002 |
| 025-0312 | 44.730 | -79.169 | YELLOW | NORTH | Orillia, ON | *P. resinosa* | 20-Aug-2002 |
| 025-0335 | 46.017 | -77.450 | YELLOW | NORTH | Chalk River, ON | *P. resinosa* | 24-Aug-2002 |
| 025-0339B | 46.383 | -82.650 | YELLOW | NORTH | Elliot Lake, ON | *P. resinosa* | 15-Aug-2002 |
| 075-04 | 28.096 | -81.275 | YELLOW | SOUTH | Canoe Creek, FL | *P. elliottii* | 10-Jul-2004 |
| 076-04 | 28.096 | -81.275 | YELLOW | SOUTH | Canoe Creek, FL | *P. elliottii* | 10-Jul-2004 |
| 077-04 | 26.923 | -81.336 | YELLOW | SOUTH | Palmdale, FL | *P. elliottii* | 11-Jul-2004 |
| 078-04 | 26.923 | -81.336 | YELLOW | SOUTH | Palmdale, FL | *P. elliottii* | 11-Jul-2004 |
| 079-04 | 26.923 | -81.336 | YELLOW | SOUTH | Palmdale, FL | *P. elliottii* | 11-Jul-2004 |
| 085-04 | 29.718 | -82.457 | YELLOW | SOUTH | Gainesville, FL | *P. palustris* | 12-Jul-2004 |
| 086-04 | 29.718 | -82.457 | YELLOW | SOUTH | Gainesville, FL | *P. palustris* | 12-Jul-2004 |
| 087-04 | 29.748 | -82.477 | YELLOW | SOUTH | Gainesville, FL | *P. taeda* | 13-Jul-2004 |
| 088-04 | 29.748 | -82.477 | YELLOW | SOUTH | Gainesville, FL | *P. palustris* | 13-Jul-2004 |
| 096-04 | 31.498 | -84.593 | YELLOW | SOUTH | Morgan, GA | *P. taeda* | 14-Jul-2004 |
| 097-04 | 31.498 | -84.593 | YELLOW | SOUTH | Morgan, GA | *P. glabra* | 14-Jul-2004 |
| 102-04 | 31.555 | -83.989 | YELLOW | SOUTH | Sylvester, GA | *P. elliottii* | 15-Jul-2004 |
| 106-04 | 32.074 | -83.761 | YELLOW | SOUTH | Vienna, GA | *P. elliottii* | 17-Jul-2004 |
| 116-04 | 36.039 | -85.109 | YELLOW | CENTRAL | Crossville, TN | *P. virginiana* | 19-Jul-2004 |
| 125-02 | 43.115 | -71.100 | YELLOW | CENTRAL | Nottingham, NH | *P. sylvestris* | 21-Jul-2002 |

| | | | | | | | |
|---|---|---|---|---|---|---|---|
| 129-02 | 43.115 | -71.100 | YELLOW | CENTRAL | Nottingham, NH | *P. sylvestris* | 21-Jul-2002 |
| 132-04 | 38.716 | -76.064 | WHITE | CENTRAL | Trappe, MD | *P. virginiana* | 23-Jul-2004 |
| 133-04 | 38.716 | -76.064 | WHITE | CENTRAL | Trappe, MD | *P. taeda* | 23-Jul-2004 |
| 145-04 | 43.781 | -71.170 | YELLOW | CENTRAL | Ossippee, NH | *P. rigida* | 27-Jul-2004 |
| 164-02 | 45.073 | -77.710 | YELLOW | NORTH | Bancroft, ON | *P. resinosa* | 9-Aug-2002 |
| 168-02 | 44.856 | -77.859 | YELLOW | NORTH | Apsley, ON | *P. banksiana* | 9-Aug-2002 |
| 168-04 | 43.685 | -71.117 | YELLOW | CENTRAL | Ossippee, NH | *P. rigida* | 30-Jul-2004 |
| 171-02 | 44.730 | -79.169 | YELLOW | NORTH | Sebrite, ON | *P. resinosa* | 10-Aug-2002 |
| 173-02 | 44.730 | -79.169 | YELLOW | NORTH | Sebrite, ON | *P. resinosa* | 10-Aug-2002 |
| 174-02 | 44.730 | -79.169 | YELLOW | NORTH | Sebrite, ON | *P. resinosa* | 10-Aug-2002 |
| 174-03A | 29.680 | -83.257 | YELLOW | SOUTH | Dixie Co, FL | *P. taeda* | 23-Nov-2003 |
| 175-02 | 44.730 | -79.169 | YELLOW | NORTH | Sebrite, ON | *P. resinosa* | 10-Aug-2002 |
| 176-02 | 44.730 | -79.169 | YELLOW | NORTH | Sebrite, ON | *P. resinosa* | 10-Aug-2002 |
| 177-02 | 44.730 | -79.169 | YELLOW | NORTH | Sebrite, ON | *P. resinosa* | 10-Aug-2002 |
| 178-02 | 44.730 | -79.169 | YELLOW | NORTH | Sebrite, ON | *P. resinosa* | 10-Aug-2002 |
| 178-03 | 30.428 | -85.603 | YELLOW | SOUTH | Crystal Lake, FL | *P. palustris* | 24-Nov-2003 |
| 180-03 | 28.787 | -81.982 | YELLOW | SOUTH | Lake Co, FL | *P. taeda* | 25-Nov-2003 |
| 183-03 | 26.871 | -81.521 | YELLOW | SOUTH | Glades Co, FL | *P. palustris* | 26-Nov-2003 |
| 185-03 | 26.923 | -81.337 | YELLOW | SOUTH | Palmdale, FL | *P. elliottii* | 26-Nov-2003 |
| 188-04 | 44.759 | -91.457 | YELLOW | NORTH | Eau Claire, WI | *P. banksiana* | 15-Aug-2004 |
| 196-04 | 43.912 | -90.866 | YELLOW | NORTH | Sparta, WI | *P. banksiana* | 15-Aug-2004 |
| 207-04 | 45.975 | -90.496 | YELLOW | NORTH | Park Falls, WI | *P. banksiana* | 17-Aug-2004 |
| 339-02 | 46.348 | -79.334 | YELLOW | NORTH | North Bay, ON | *P. banksiana* | 19-Aug-2002 |
| 342-02 | 46.395 | -79.244 | YELLOW | NORTH | North Bay, ON | *P. banksiana* | 19-Aug-2002 |
| 343-02 | 46.395 | -79.244 | YELLOW | NORTH | North Bay, ON | *P. banksiana* | 19-Aug-2002 |
| 344-02 | 46.395 | -79.244 | YELLOW | NORTH | North Bay, ON | *P. resinosa* | 19-Aug-2002 |
| 345-02 | 46.395 | -79.244 | YELLOW | NORTH | North Bay, ON | *P. resinosa* | 19-Aug-2002 |
| 349-02 | 46.378 | -78.867 | YELLOW | NORTH | Mattawan, ON | *P. resinosa* | 19-Aug-2002 |
| 352-02 | 46.378 | -78.867 | YELLOW | NORTH | Mattawan, ON | *P. resinosa* | 19-Aug-2002 |

| | | | | | | | |
|---|---|---|---|---|---|---|---|
| 372-02 | 41.874 | -70.652 | WHITE | CENTRAL | Plymouth, MA | *P. rigida* | 25-Sep-2002 |
| 379-02 | 41.866 | -70.658 | WHITE | CENTRAL | Plymouth, MA | *P. rigida* | 25-Sep-2002 |
| CAN002 | 44.736 | -79.161 | YELLOW | NORTH | Lindsay, ON | *P. banksiana* | 27-Jul-2014 |
| CAN003 | 44.736 | -79.161 | YELLOW | NORTH | Lindsay, ON | *P. banksiana* | 27-Jul-2014 |
| CAN004 | 44.736 | -79.161 | YELLOW | NORTH | Lindsay, ON | *P. banksiana* | 27-Jul-2014 |
| CAN005 | 44.736 | -79.161 | YELLOW | NORTH | Lindsay, ON | *P. resinosa* | 27-Jul-2014 |
| CAN006 | 44.732 | -79.169 | YELLOW | NORTH | Lindsay, ON | *P. resinosa* | 27-Jul-2014 |
| CAN007 | 44.863 | -78.113 | YELLOW | NORTH | Harcourt, ON | *P. resinosa* | 27-Jul-2014 |
| CAN008 | 44.863 | -78.113 | YELLOW | NORTH | Harcourt, ON | *P. resinosa* | 27-Jul-2014 |
| CAN009 | 44.863 | -78.113 | YELLOW | NORTH | Harcourt, ON | *P. strobus* | 27-Jul-2014 |
| CAN011 | 45.316 | -77.764 | YELLOW | NORTH | Combermere, ON | *P. strobus* | 28-Jul-2014 |
| CAN012 | 45.316 | -77.764 | YELLOW | NORTH | Combermere, ON | *P. strobus* | 28-Jul-2014 |
| CAN013 | 45.316 | -77.764 | YELLOW | NORTH | Combermere, ON | *P. resinosa* | 28-Jul-2014 |
| CAN014 | 45.316 | -77.764 | YELLOW | NORTH | Combermere, ON | *P. resinosa* | 28-Jul-2014 |
| CAN015 | 45.316 | -77.764 | YELLOW | NORTH | Combermere, ON | *P. resinosa* | 28-Jul-2014 |
| CAN016 | 45.316 | -77.764 | YELLOW | NORTH | Combermere, ON | *P. resinosa* | 28-Jul-2014 |
| CAN017 | 45.316 | -77.764 | YELLOW | NORTH | Combermere, ON | *P. resinosa* | 28-Jul-2014 |
| CAN018 | 45.316 | -77.764 | YELLOW | NORTH | Combermere, ON | *P. resinosa* | 28-Jul-2014 |
| CAN019 | 45.316 | -77.764 | YELLOW | NORTH | Combermere, ON | *P. resinosa* | 28-Jul-2014 |
| CAN020 | 45.316 | -77.764 | YELLOW | NORTH | Combermere, ON | *P. resinosa* | 28-Jul-2014 |
| CAN021 | 45.483 | -77.675 | YELLOW | NORTH | Barry's Bay, ON | *P. resinosa* | 28-Jul-2014 |
| CAN022 | 45.483 | -77.675 | YELLOW | NORTH | Barry's Bay, ON | *P. resinosa* | 28-Jul-2014 |
| CAN023 | 45.483 | -77.675 | YELLOW | NORTH | Barry's Bay, ON | *P. resinosa* | 28-Jul-2014 |
| CAN024 | 45.483 | -77.675 | YELLOW | NORTH | Barry's Bay, ON | *P. resinosa* | 28-Jul-2014 |
| CAN025 | 45.483 | -77.675 | YELLOW | NORTH | Barry's Bay, ON | *P. resinosa* | 28-Jul-2014 |
| CAN026 | 45.486 | -77.672 | YELLOW | NORTH | Barry's Bay, ON | *P. resinosa* | 28-Jul-2014 |
| CAN027 | 45.486 | -77.672 | YELLOW | NORTH | Barry's Bay, ON | *P. resinosa* | 28-Jul-2014 |
| CAN028 | 45.486 | -77.672 | YELLOW | NORTH | Barry's Bay, ON | *P. resinosa* | 28-Jul-2014 |
| CAN029 | 45.486 | -77.672 | YELLOW | NORTH | Barry's Bay, ON | *P. resinosa* | 28-Jul-2014 |

| | | | | | | | |
|---|---|---|---|---|---|---|---|
| CAN030 | 45.486 | -77.672 | YELLOW | NORTH | Barry's Bay, ON | *P. resinosa* | 28-Jul-2014 |
| CAN031 | 45.486 | -77.672 | YELLOW | NORTH | Barry's Bay, ON | *P. resinosa* | 28-Jul-2014 |
| CAN032 | 45.486 | -77.672 | YELLOW | NORTH | Barry's Bay, ON | *P. resinosa* | 28-Jul-2014 |
| CAN033 | 45.486 | -77.672 | YELLOW | NORTH | Barry's Bay, ON | *P. resinosa* | 28-Jul-2014 |
| CAN034 | 45.486 | -77.672 | YELLOW | NORTH | Barry's Bay, ON | *P. resinosa* | 28-Jul-2014 |
| CAN035 | 45.486 | -77.672 | YELLOW | NORTH | Barry's Bay, ON | *P. resinosa* | 28-Jul-2014 |
| CAN036 | 45.798 | -77.192 | YELLOW | NORTH | Laurentian Valley, ON | *P. resinosa* | 29-Jul-2014 |
| CAN037a | 45.834 | -77.237 | YELLOW | NORTH | Laurentian Valley, ON | *P. resinosa* | 29-Jul-2014 |
| CAN038 | 45.834 | -77.237 | YELLOW | NORTH | Laurentian Valley, ON | *P. resinosa* | 29-Jul-2014 |
| CAN039 | 45.834 | -77.237 | YELLOW | NORTH | Laurentian Valley, ON | *P. resinosa* | 29-Jul-2014 |
| CAN040 | 46.288 | -78.812 | YELLOW | NORTH | Papineau-Cameron, ON | *P. resinosa* | 29-Jul-2014 |
| CAN041 | 46.370 | -81.384 | YELLOW | NORTH | Greater Sudbury, ON | *P. resinosa* | 30-Jul-2014 |
| CAN042 | 46.288 | -81.793 | YELLOW | NORTH | Baldwin, ON | *P. resinosa* | 30-Jul-2014 |
| CAN043 | 44.601 | -84.712 | YELLOW | NORTH | Grayling, MI | *P. banksiana* | 31-Jul-2014 |
| CAN044 | 44.601 | -84.712 | YELLOW | NORTH | Grayling, MI | *P. banksiana* | 31-Jul-2014 |
| CAN045 | 44.601 | -84.712 | YELLOW | NORTH | Grayling, MI | *P. banksiana* | 31-Jul-2014 |
| CAN046 | 44.601 | -84.712 | YELLOW | NORTH | Grayling, MI | *P. banksiana* | 31-Jul-2014 |
| CAN047 | 44.601 | -84.712 | YELLOW | NORTH | Grayling, MI | *P. banksiana* | 31-Jul-2014 |
| CAN048 | 44.657 | -84.696 | YELLOW | NORTH | Grayling, MI | *P. resinosa* | 1-Aug-2014 |
| CAN049 | 44.657 | -84.696 | YELLOW | NORTH | Grayling, MI | *P. resinosa* | 1-Aug-2014 |
| CAN050 | 44.657 | -84.696 | YELLOW | NORTH | Grayling, MI | *P. resinosa* | 1-Aug-2014 |
| CAN051 | 44.657 | -84.696 | YELLOW | NORTH | Grayling, MI | *P. sylvestris* | 1-Aug-2014 |
| CAN052 | 44.657 | -84.696 | YELLOW | NORTH | Grayling, MI | *P. strobus* | 1-Aug-2014 |
| CAN053 | 44.657 | -84.696 | YELLOW | NORTH | Grayling, MI | *P. strobus* | 1-Aug-2014 |
| CAN054 | 44.657 | -84.696 | YELLOW | NORTH | Grayling, MI | *P. banksiana* | 1-Aug-2014 |
| CAN055 | 44.657 | -84.696 | YELLOW | NORTH | Grayling, MI | *P. banksiana* | 1-Aug-2014 |
| CAN056 | 44.657 | -84.696 | YELLOW | NORTH | Grayling, MI | *P. banksiana* | 1-Aug-2014 |
| CAN057 | 44.122 | -85.471 | YELLOW | NORTH | Tustin, MI | *P. banksiana* | 1-Aug-2014 |
| CAN059 | 43.759 | -85.741 | YELLOW | NORTH | Bitely, MI | *P. sylvestris* | 1-Aug-2014 |

| | | | | | | | |
|---|---|---|---|---|---|---|---|
| CAN060 | 43.789 | -85.740 | YELLOW | NORTH | Bitely, MI | *P. sylvestris* | 1-Aug-2014 |
| CAN061a | 43.789 | -85.740 | YELLOW | NORTH | Bitely, MI | *P. banksiana* | 1-Aug-2014 |
| CAN062 | 43.769 | -85.741 | YELLOW | NORTH | Bitely, MI | *P. banksiana* | 1-Aug-2014 |
| CAN063 | 46.214 | -83.099 | YELLOW | NORTH | Blind River, ON | *P. resinosa* | 13-Aug-2014 |
| CAN064 | 46.220 | -83.108 | YELLOW | NORTH | Blind River, ON | *P. resinosa* | 15-Aug-2014 |
| CAN065 | 46.220 | -83.108 | YELLOW | NORTH | Blind River, ON | *P. resinosa* | 15-Aug-2014 |
| CAN066 | 46.220 | -83.108 | YELLOW | NORTH | Blind River, ON | *P. resinosa* | 15-Aug-2014 |
| CAN067 | 46.220 | -83.108 | YELLOW | NORTH | Blind River, ON | *P. resinosa* | 15-Aug-2014 |
| CAN068 | 46.220 | -83.108 | YELLOW | NORTH | Blind River, ON | *P. resinosa* | 15-Aug-2014 |
| CAN069 | 46.214 | -83.099 | YELLOW | NORTH | Blind River, ON | *P. resinosa* | 15-Aug-2014 |
| CAN070 | 46.214 | -83.099 | YELLOW | NORTH | Blind River, ON | *P. resinosa* | 15-Aug-2014 |
| CAN071 | 46.214 | -83.099 | YELLOW | NORTH | Blind River, ON | *P. resinosa* | 15-Aug-2014 |
| CAN072 | 46.214 | -83.099 | YELLOW | NORTH | Blind River, ON | *P. resinosa* | 15-Aug-2014 |
| CAN073 | 46.214 | -83.099 | YELLOW | NORTH | Blind River, ON | *P. resinosa* | 15-Aug-2014 |
| CAN074 | 46.214 | -83.099 | YELLOW | NORTH | Blind River, ON | *P. resinosa* | 15-Aug-2014 |
| CAN075 | 46.201 | -82.358 | YELLOW | NORTH | Spanish, ON | *P. resinosa* | 15-Aug-2014 |
| CAN076 | 46.201 | -82.358 | YELLOW | NORTH | Spanish, ON | *P. resinosa* | 15-Aug-2014 |
| CAN077 | 46.212 | -82.081 | YELLOW | NORTH | Sables-Spanish River, ON | *P. banksiana* | 15-Aug-2014 |
| CAN078 | 46.221 | -82.024 | YELLOW | NORTH | Sables-Spanish River, ON | *P. resinosa* | 15-Aug-2014 |
| CAN079 | 46.221 | -82.024 | YELLOW | NORTH | Sables-Spanish River, ON | *P. sylvestris* | 15-Aug-2014 |
| CAN080 | 46.569 | -81.232 | YELLOW | NORTH | Chelmsford, ON | *P. banksiana* | 16-Aug-2014 |
| CAN082 | 46.587 | -81.382 | YELLOW | NORTH | Dowling, ON | *P. resinosa* | 16-Aug-2014 |
| CAN083 | 46.732 | -81.589 | YELLOW | NORTH | Cartier, ON | *P. resinosa* | 16-Aug-2014 |
| CAN084 | 46.732 | -81.589 | YELLOW | NORTH | Cartier, ON | *P. banksiana* | 16-Aug-2014 |
| CAN090 | 47.032 | -83.152 | YELLOW | NORTH | Algmoa, Unorganized, North Part, ON | *P. banksiana* | 17-Aug-2014 |
| CAN092 | 46.719 | -83.425 | YELLOW | NORTH | Algmoa, Unorganized, ON | *P. resinosa* | 17-Aug-2014 |
| CAN093 | 46.719 | -83.425 | YELLOW | NORTH | Algmoa, Unorganized, ON | *P. resinosa* | 17-Aug-2014 |
| CAN094 | 46.719 | -83.425 | YELLOW | NORTH | Algmoa, Unorganized, ON | *P. resinosa* | 17-Aug-2014 |

| | | | | | | | |
|---|---|---|---|---|---|---|---|
| CAN095 | 46.719 | -83.425 | YELLOW | NORTH | Algmoa, Unorganized, ON | *P. strobus* | 17-Aug-2014 |
| CAN096 | 46.719 | -83.425 | YELLOW | NORTH | Algmoa, Unorganized, ON | *P. strobus* | 17-Aug-2014 |
| CAN097 | 46.719 | -83.425 | YELLOW | NORTH | Algmoa, Unorganized, ON | *P. strobus* | 17-Aug-2014 |
| CAN098 | 44.335 | -90.730 | YELLOW | NORTH | Black River Falls, WI | *P. banksiana/P. strobus* | 21-Aug-2014 |
| CAN099 | 43.799 | -85.736 | YELLOW | NORTH | Bitely, MI | *P. resinosa* | 11-Sep-2014 |
| CAN100 | 44.704 | -84.906 | YELLOW | NORTH | Grayling, MI | *P. banksiana* | 11-Sep-2014 |
| CAN101 | 45.924 | -86.302 | YELLOW | NORTH | Manistique, MI | *P. banksiana* | 12-Sep-2014 |
| CAN102 | 45.892 | -86.520 | YELLOW | NORTH | Manistique, MI | *P. banksiana* | 12-Sep-2014 |
| CAN103 | 44.348 | -90.346 | YELLOW | NORTH | City Point, WI | *P. banksiana* | 13-Sep-2014 |
| CAN104 | 44.341 | -90.411 | YELLOW | NORTH | Pittsville, WI | *P. banksiana* | 13-Sep-2014 |
| CAN105 | 44.341 | -90.411 | YELLOW | NORTH | Pittsville, WI | *P. banksiana* | 13-Sep-2014 |
| CAN106 | 44.341 | -90.411 | YELLOW | NORTH | Pittsville, WI | *P. banksiana* | 13-Sep-2014 |
| CAN107 | 44.130 | -90.393 | YELLOW | NORTH | Warrens, WI | *P. banksiana* | 13-Sep-2014 |
| CAN108 | 44.130 | -90.393 | YELLOW | NORTH | Warrens, WI | *P. banksiana* | 13-Sep-2014 |
| CAN109 | 44.199 | -90.136 | YELLOW | NORTH | Necedah, WI | *P. banksiana* | 13-Sep-2014 |
| CAN111 | 44.036 | -90.082 | YELLOW | NORTH | Necedah, WI | *P. banksiana* | 13-Sep-2014 |
| LL002 | 38.014 | -84.504 | YELLOW | CENTRAL | Lexington, KY | *P. echinata* | 13-Jun-2012 |
| LL003 | 38.014 | -84.504 | YELLOW | CENTRAL | Lexington, KY | *P. echinata* | 13-Jun-2012 |
| LL004 | 38.014 | -84.504 | YELLOW | CENTRAL | Lexington, KY | *P. echinata* | 13-Jun-2012 |
| LL005 | 38.014 | -84.504 | YELLOW | CENTRAL | Lexington, KY | *P. rigida* | 15-Jun-2012 |
| LL006 | 38.014 | -84.504 | YELLOW | CENTRAL | Lexington, KY | *P. rigida* | 15-Jun-2012 |
| LL007 | 38.014 | -84.504 | YELLOW | CENTRAL | Lexington, KY | *P. virginiana* | 15-Jun-2012 |
| LL009 | 35.929 | -84.914 | YELLOW | CENTRAL | Crossville, TN | *P. virginiana* | 11-Jul-2013 |
| LL010 | 36.928 | -84.619 | YELLOW | CENTRAL | Bronston, KY | *P. virginiana* | 14-Jul-2013 |
| LL011 | 37.071 | -84.211 | YELLOW | CENTRAL | London, KY | *P. echinata* | 16-Jul-2013 |
| LL012 | 37.071 | -84.211 | YELLOW | CENTRAL | London, KY | *P. echinata* | 16-Jul-2013 |
| LL013 | 37.071 | -84.211 | YELLOW | CENTRAL | London, KY | *P. echinata* | 16-Jul-2013 |
| LL014 | 37.071 | -84.211 | YELLOW | CENTRAL | London, KY | *P. echinata* | 16-Jul-2013 |

| | | | | | | | |
|---|---|---|---|---|---|---|---|
| LL015 | 37.071 | -84.211 | YELLOW | CENTRAL | London, KY | *P. echinata* | 16-Jul-2013 |
| LL017 | 38.014 | -84.504 | YELLOW | CENTRAL | Lexington, KY | *P. echinata* | 18-Jul-2013 |
| LL030 | 34.115 | -79.940 | YELLOW | SOUTH | Florence county, SC | *P. palustris* | 5-Aug-2013 |
| LL031 | 40.550 | -74.431 | MIXED | CENTRAL | Piscataway, NJ | *P. sylvestris* | 14-Aug-2013 |
| LL032 | 46.183 | -82.948 | YELLOW | NORTH | Blind River, ON | *P. resinosa* | 15-Aug-2013 |
| LL033 | 46.183 | -82.948 | YELLOW | NORTH | Blind River, ON | *NA* | 15-Aug-2013 |
| LL034 | 44.939 | -75.671 | YELLOW | NORTH | Oxford Mills, ON | *P. resinosa* | 16-Aug-2013 |
| LL035 | 46.296 | -83.550 | YELLOW | NORTH | Little Rapids, ON | *P. resinosa* | 18-Aug-2013 |
| LL036 | 46.422 | -83.375 | YELLOW | NORTH | Whamcliffe, ON | *P. resinosa* | 18-Aug-2013 |
| LL037 | 46.207 | -83.060 | YELLOW | NORTH | Mississaugi River, ON | *P. resinosa* | 19-Aug-2013 |
| LL038 | 46.183 | -82.948 | YELLOW | NORTH | Blind River, ON | *P. resinosa* | 15-Aug-2013 |
| LL039 | 46.791 | -84.030 | YELLOW | NORTH | Searchmont, ON | *P. banksiana* | 22-Aug-2013 |
| LL040 | 46.791 | -84.030 | YELLOW | NORTH | Searchmont, ON | *P. banksiana* | 22-Aug-2013 |
| LL041 | 46.372 | -82.606 | YELLOW | NORTH | Elliot Lake, ON | *P. resinosa* | 23-Aug-2013 |
| LL042 | 37.249 | -77.725 | WHITE | CENTRAL | Amelia, VA | *P. echinata* | 18-Sep-2013 |
| LL045 | 29.474 | -82.861 | YELLOW | SOUTH | Chiefland, FL | *P. palustris* | 7-Apr-2014 |
| LL047 | 38.014 | -84.504 | YELLOW | CENTRAL | Lexington, KY | *P. echinata* | 9-Jun-2014 |
| LL048 | 38.014 | -84.504 | YELLOW | CENTRAL | Lexington, KY | *P. echinata* | 9-Jun-2014 |
| LL049 | 38.014 | -84.504 | YELLOW | CENTRAL | Lexington, KY | *P. echinata* | 9-Jun-2014 |
| LL050 | 38.014 | -84.504 | YELLOW | CENTRAL | Lexington, KY | *P. echinata* | 9-Jun-2014 |
| LL051 | 38.014 | -84.504 | YELLOW | CENTRAL | Lexington, KY | *P. echinata* | 9-Jun-2014 |
| LL052 | 38.014 | -84.504 | YELLOW | CENTRAL | Lexington, KY | *P. virginiana* | 9-Jun-2014 |
| LL053 | 38.014 | -84.504 | YELLOW | CENTRAL | Lexington, KY | *P. virginiana* | 9-Jun-2014 |
| LL054 | 38.014 | -84.504 | YELLOW | CENTRAL | Lexington, KY | *P. virginiana* | 9-Jun-2014 |
| LL055 | 38.014 | -84.504 | YELLOW | CENTRAL | Lexington, KY | *P. virginiana* | 9-Jun-2014 |
| LL056 | 38.014 | -84.504 | YELLOW | CENTRAL | Lexington, KY | *P. rigida* | 9-Jun-2014 |
| LL057 | 38.014 | -84.504 | YELLOW | CENTRAL | Lexington, KY | *P. rigida* | 9-Jun-2014 |
| LL058 | 38.014 | -84.504 | YELLOW | CENTRAL | Lexington, KY | *P. virginiana* | 19-Jun-2014 |
| LL059 | 38.014 | -84.504 | YELLOW | CENTRAL | Lexington, KY | *P. virginiana* | 19-Jun-2014 |

| | | | | | | | |
|---|---|---|---|---|---|---|---|
| LL060 | 38.014 | -84.504 | YELLOW | CENTRAL | Lexington, KY | *P. virginiana* | 19-Jun-2014 |
| LL061 | 38.014 | -84.504 | YELLOW | CENTRAL | Lexington, KY | *P. virginiana* | 19-Jun-2014 |
| LL062 | 38.014 | -84.504 | YELLOW | CENTRAL | Lexington, KY | *P. echinata* | 19-Jun-2014 |
| LL063 | 38.014 | -84.504 | YELLOW | CENTRAL | Lexington, KY | *P. echinata* | 19-Jun-2014 |
| LL064 | 38.014 | -84.504 | YELLOW | CENTRAL | Lexington, KY | *P. rigida* | 19-Jun-2014 |
| LL065 | 38.014 | -84.504 | YELLOW | CENTRAL | Lexington, KY | *P. rigida* | 19-Jun-2014 |
| LL066 | 38.014 | -84.504 | YELLOW | CENTRAL | Lexington, KY | *P. rigida* | 19-Jun-2014 |
| LL067 | 38.014 | -84.504 | YELLOW | CENTRAL | Lexington, KY | *P. rigida* | 19-Jun-2014 |
| LL068 | 35.782 | -78.640 | YELLOW | NORTH | Raleigh, NC | *P. palustris* | 25-Jun-2014 |
| LL069 | 38.014 | -84.504 | YELLOW | CENTRAL | Lexington, KY | *P. virginiana* | 27-Jun-2014 |
| LL070 | 38.014 | -84.504 | YELLOW | CENTRAL | Lexington, KY | *P. rigida* | 27-Jun-2014 |
| LL071 | 38.014 | -84.504 | YELLOW | CENTRAL | Lexington, KY | *P. echinata* | 27-Jun-2014 |
| LL072 | 38.014 | -84.504 | YELLOW | CENTRAL | Lexington, KY | *P. virginiana* | 27-Jun-2014 |
| LL073 | 38.014 | -84.504 | YELLOW | CENTRAL | Lexington, KY | *P. echinata* | 27-Jun-2014 |
| LL074 | 38.014 | -84.504 | YELLOW | CENTRAL | Lexington, KY | *P. echinata* | 30-Jun-2014 |
| LL075 | 38.014 | -84.504 | YELLOW | CENTRAL | Lexington, KY | *P. rigida* | 30-Jun-2014 |
| LL076 | 38.014 | -84.504 | YELLOW | CENTRAL | Lexington, KY | *P. virginiana* | 30-Jun-2014 |
| LL077 | 38.014 | -84.504 | YELLOW | CENTRAL | Lexington, KY | *P. virginiana* | 30-Jun-2014 |
| LL078 | 38.014 | -84.504 | YELLOW | CENTRAL | Lexington, KY | *P. rigida* | 30-Jun-2014 |
| LL081 | 38.014 | -84.504 | YELLOW | CENTRAL | Lexington, KY | *P. rigida* | 24-Jul-2014 |
| LL082 | 41.645 | -70.230 | WHITE | CENTRAL | West Yarmouth, MA | *P. rigida* | 5-Aug-2014 |
| LL083 | 41.645 | -70.230 | WHITE | CENTRAL | West Yarmouth, MA | *P. rigida* | 5-Aug-2014 |
| LL084 | 41.645 | -70.230 | WHITE | CENTRAL | West Yarmouth, MA | *P. rigida* | 5-Aug-2014 |
| LL086 | 38.014 | -84.504 | YELLOW | CENTRAL | Lexington, KY | *P. echinata* | 28-Aug-2014 |
| LL087 | 38.014 | -84.504 | YELLOW | CENTRAL | Lexington, KY | *P. echinata* | 28-Aug-2014 |
| LL088 | 38.014 | -84.504 | YELLOW | CENTRAL | Lexington, KY | *P. echinata* | 28-Aug-2014 |
| LL089 | 38.014 | -84.504 | YELLOW | CENTRAL | Lexington, KY | *P. echinata* | 28-Aug-2014 |
| LL090 | 38.014 | -84.504 | YELLOW | CENTRAL | Lexington, KY | *P. echinata* | 28-Aug-2014 |
| LL091 | 38.014 | -84.504 | YELLOW | CENTRAL | Lexington, KY | *P. echinata* | 28-Aug-2014 |

| | | | | | | | |
|---|---|---|---|---|---|---|---|
| LL092 | 38.014 | -84.504 | YELLOW | CENTRAL | Lexington, KY | *P. virginiana* | 28-Aug-2014 |
| LL093 | 38.014 | -84.504 | YELLOW | CENTRAL | Lexington, KY | *P. virginiana* | 28-Aug-2014 |
| LL094 | 38.014 | -84.504 | YELLOW | CENTRAL | Lexington, KY | *P. virginiana* | 28-Aug-2014 |
| LL095 | 38.014 | -84.504 | YELLOW | CENTRAL | Lexington, KY | *P. virginiana* | 28-Aug-2014 |
| LL096 | 38.014 | -84.504 | YELLOW | CENTRAL | Lexington, KY | *P. virginiana* | 28-Aug-2014 |
| LL097 | 38.044 | -84.497 | YELLOW | CENTRAL | Lexington, KY | *P. mugho* | 3-Sep-2014 |
| LL098 | 38.044 | -84.497 | YELLOW | CENTRAL | Lexington, KY | *P. mugho* | 3-Sep-2014 |
| LL099 | 38.044 | -84.497 | YELLOW | CENTRAL | Lexington, KY | *P. mugho* | 3-Sep-2014 |
| LL100 | 38.023 | -84.494 | YELLOW | CENTRAL | Lexington, KY | *P. nigra* | 3-Sep-2014 |
| LL101 | 38.014 | -84.504 | YELLOW | CENTRAL | Lexington, KY | *P. rigida* | 4-Sep-2014 |
| LL102 | 38.014 | -84.504 | YELLOW | CENTRAL | Lexington, KY | *P. rigida* | 4-Sep-2014 |
| LL103 | 38.014 | -84.504 | YELLOW | CENTRAL | Lexington, KY | *P. echinata* | 4-Sep-2014 |
| LL104 | 38.014 | -84.504 | YELLOW | CENTRAL | Lexington, KY | *P. echinata* | 4-Sep-2014 |
| LL105 | 38.014 | -84.504 | YELLOW | CENTRAL | Lexington, KY | *P. echinata* | 4-Sep-2014 |
| LL106 | 38.014 | -84.504 | YELLOW | CENTRAL | Lexington, KY | *P. virginiana* | 4-Sep-2014 |
| LL107 | 38.014 | -84.504 | YELLOW | CENTRAL | Lexington, KY | *P. virginiana* | 4-Sep-2014 |
| LL108 | 38.014 | -84.504 | YELLOW | CENTRAL | Lexington, KY | *P. virginiana* | 4-Sep-2014 |
| LL109 | 38.024 | -84.532 | YELLOW | CENTRAL | Lexington, KY | *P. mugho* | 21-Sep-2014 |
| LL110 | 38.024 | -84.532 | YELLOW | CENTRAL | Lexington, KY | *P. mugho* | 21-Sep-2014 |
| LL111 | 38.024 | -84.532 | YELLOW | CENTRAL | Lexington, KY | *P. mugho* | 21-Sep-2014 |
| LL112 | 38.024 | -84.532 | YELLOW | CENTRAL | Lexington, KY | *P. mugho* | 21-Sep-2014 |
| LL113 | 38.024 | -84.532 | YELLOW | CENTRAL | Lexington, KY | *P. mugho* | 21-Sep-2014 |
| LL116 | 38.014 | -84.504 | YELLOW | CENTRAL | Lexington, KY | *P. rigida* | 2-Jun-2015 |
| LL117 | 37.984 | -84.418 | YELLOW | CENTRAL | Lexington, KY | *P. taeda* | 4-Jun-2015 |
| LL121 | 38.014 | -84.504 | YELLOW | CENTRAL | Lexington, KY | *P. rigida* | 17-Jun-2015 |
| LL122 | 38.014 | -84.504 | YELLOW | CENTRAL | Lexington, KY | *P. rigida* | 17-Jun-2015 |
| LL123 | 38.033 | -84.507 | YELLOW | CENTRAL | Lexington, KY | *P. mugho* | 18-Jun-2015 |
| LL124 | 38.033 | -84.507 | YELLOW | CENTRAL | Lexington, KY | *P. mugho* | 18-Jun-2015 |
| LL125 | 37.984 | -84.418 | YELLOW | CENTRAL | Lexington, KY | *P. taeda* | 18-Jun-2015 |

| | | | | | | | |
|---|---|---|---|---|---|---|---|
| LL126 | 37.984 | -84.418 | YELLOW | CENTRAL | Lexington, KY | *P. taeda* | 18-Jun-2015 |
| LL127 | 37.984 | -84.418 | YELLOW | CENTRAL | Lexington, KY | *P. taeda* | 18-Jun-2015 |
| LL128 | 38.044 | -84.497 | YELLOW | CENTRAL | Lexington, KY | *P. mugho* | 18-Jun-2015 |
| LL129 | 38.044 | -84.497 | YELLOW | CENTRAL | Lexington, KY | *P. mugho* | 18-Jun-2015 |
| LL130 | 38.044 | -84.497 | YELLOW | CENTRAL | Lexington, KY | *P. mugho* | 18-Jun-2015 |
| LL131 | 38.044 | -84.497 | YELLOW | CENTRAL | Lexington, KY | *P. mugho* | 18-Jun-2015 |
| LL132 | 38.024 | -84.532 | YELLOW | CENTRAL | Lexington, KY | *P. mugho* | 19-Jun-2015 |
| LL133 | 38.024 | -84.532 | YELLOW | CENTRAL | Lexington, KY | *P. mugho* | 19-Jun-2015 |
| LL134 | 38.024 | -84.532 | YELLOW | CENTRAL | Lexington, KY | *P. mugho* | 19-Jun-2015 |
| LL135 | 38.024 | -84.532 | YELLOW | CENTRAL | Lexington, KY | *P. mugho* | 19-Jun-2015 |
| LL136 | 38.014 | -84.504 | YELLOW | CENTRAL | Lexington, KY | *P. virginiana* | 19-Jun-2015 |
| LL137 | 38.014 | -84.504 | YELLOW | CENTRAL | Lexington, KY | *P. virginiana* | 19-Jun-2015 |
| LL138 | 38.014 | -84.504 | YELLOW | CENTRAL | Lexington, KY | *P. echinata* | 19-Jun-2015 |
| LL139 | 38.014 | -84.504 | YELLOW | CENTRAL | Lexington, KY | *P. echinata* | 19-Jun-2015 |
| LL140 | 38.014 | -84.504 | YELLOW | CENTRAL | Lexington, KY | *P. virginiana* | 19-Jun-2015 |
| LL142 | 35.980 | -85.015 | YELLOW | CENTRAL | Crossville, TN | *P. virginiana* | 23-Jun-2015 |
| LL143 | 35.980 | -85.015 | YELLOW | CENTRAL | Crossville, TN | *P. virginiana* | 23-Jun-2015 |
| LL144 | 35.980 | -85.015 | YELLOW | CENTRAL | Crossville, TN | *P. virginiana* | 23-Jun-2015 |
| LL145 | 35.980 | -85.015 | YELLOW | CENTRAL | Crossville, TN | *P. virginiana* | 23-Jun-2015 |
| LL146 | 35.980 | -85.015 | YELLOW | CENTRAL | Crossville, TN | *P. virginiana* | 23-Jun-2015 |
| LL147 | 35.980 | -85.015 | YELLOW | CENTRAL | Crossville, TN | *P. virginiana* | 23-Jun-2015 |
| LL148 | 38.015 | -84.501 | YELLOW | CENTRAL | Lexington, KY | *P. virginiana* | 25-Jun-2015 |
| LL149 | 38.015 | -84.501 | YELLOW | CENTRAL | Lexington, KY | *P. virginiana* | 25-Jun-2015 |
| LL150 | 38.015 | -84.501 | YELLOW | CENTRAL | Lexington, KY | *P. virginiana* | 25-Jun-2015 |
| LL151 | 38.015 | -84.501 | YELLOW | CENTRAL | Lexington, KY | *P. virginiana* | 25-Jun-2015 |
| LL152 | 38.015 | -84.501 | YELLOW | CENTRAL | Lexington, KY | *P. virginiana* | 25-Jun-2015 |
| LL153 | 38.015 | -84.501 | YELLOW | CENTRAL | Lexington, KY | *P. virginiana* | 25-Jun-2015 |
| LL154 | 38.015 | -84.501 | YELLOW | CENTRAL | Lexington, KY | *P. virginiana* | 25-Jun-2015 |
| LL155 | 38.015 | -84.501 | YELLOW | CENTRAL | Lexington, KY | *P. virginiana* | 25-Jun-2015 |

| | | | | | | | |
|---|---|---|---|---|---|---|---|
| LL156 | 38.015 | -84.501 | YELLOW | CENTRAL | Lexington, KY | *P. virginiana* | 25-Jun-2015 |
| LL157 | 38.015 | -84.501 | YELLOW | CENTRAL | Lexington, KY | *P. virginiana* | 25-Jun-2015 |
| LL158 | 38.015 | -84.501 | YELLOW | CENTRAL | Lexington, KY | *P. virginiana* | 25-Jun-2015 |
| LL159 | 38.015 | -84.501 | YELLOW | CENTRAL | Lexington, KY | *P. virginiana* | 25-Jun-2015 |
| LL160 | 38.015 | -84.501 | YELLOW | CENTRAL | Lexington, KY | *P. virginiana* | 25-Jun-2015 |
| LL161 | 38.015 | -84.501 | YELLOW | CENTRAL | Lexington, KY | *P. virginiana* | 25-Jun-2015 |
| LL162 | 38.015 | -84.501 | YELLOW | CENTRAL | Lexington, KY | *P. virginiana* | 25-Jun-2015 |
| LL163 | 38.015 | -84.501 | YELLOW | CENTRAL | Lexington, KY | *P. virginiana* | 25-Jun-2015 |
| LL164 | 38.015 | -84.501 | YELLOW | CENTRAL | Lexington, KY | *P. virginiana* | 25-Jun-2015 |
| LL165 | 38.015 | -84.501 | YELLOW | CENTRAL | Lexington, KY | *P. virginiana* | 25-Jun-2015 |
| LL166 | 38.015 | -84.501 | YELLOW | CENTRAL | Lexington, KY | *P. virginiana* | 25-Jun-2015 |
| LL167 | 38.015 | -84.501 | YELLOW | CENTRAL | Lexington, KY | *P. virginiana* | 25-Jun-2015 |
| LL168 | 38.015 | -84.501 | YELLOW | CENTRAL | Lexington, KY | *P. virginiana* | 25-Jun-2015 |
| LL169 | 38.015 | -84.501 | YELLOW | CENTRAL | Lexington, KY | *P. virginiana* | 29-Jun-2015 |
| LL170 | 38.015 | -84.501 | YELLOW | CENTRAL | Lexington, KY | *P. virginiana* | 29-Jun-2015 |
| LL171 | 38.015 | -84.501 | YELLOW | CENTRAL | Lexington, KY | *P. virginiana* | 29-Jun-2015 |
| LL172 | 38.015 | -84.501 | YELLOW | CENTRAL | Lexington, KY | *P. virginiana* | 29-Jun-2015 |
| LL173 | 38.015 | -84.501 | YELLOW | CENTRAL | Lexington, KY | *P. virginiana* | 29-Jun-2015 |
| LL174 | 38.015 | -84.501 | YELLOW | CENTRAL | Lexington, KY | *P. virginiana* | 29-Jun-2015 |
| LL175 | 38.015 | -84.501 | YELLOW | CENTRAL | Lexington, KY | *P. virginiana* | 29-Jun-2015 |
| LL176 | 38.015 | -84.501 | YELLOW | CENTRAL | Lexington, KY | *P. virginiana* | 29-Jun-2015 |
| LL177 | 38.015 | -84.501 | YELLOW | CENTRAL | Lexington, KY | *P. virginiana* | 29-Jun-2015 |
| LL178 | 38.014 | -84.504 | YELLOW | CENTRAL | Lexington, KY | *P. virginiana* | 30-Jun-2015 |
| LL179 | 38.014 | -84.504 | YELLOW | CENTRAL | Lexington, KY | *P. echinata* | 3-Jul-2015 |
| LL180 | 38.402 | -85.586 | YELLOW | CENTRAL | Goshen, KY | *P. echinata* | 4-Jul-2015 |
| LL181 | 38.402 | -85.586 | YELLOW | CENTRAL | Goshen, KY | *P. echinata* | 4-Jul-2015 |
| LL184 | 38.014 | -84.504 | YELLOW | CENTRAL | Lexington, KY | *P. rigida* | 8-Jul-2015 |
| LL191 | 38.014 | -84.504 | YELLOW | CENTRAL | Lexington, KY | *P. echinata* | 20-Jul-2015 |
| LL193 | 37.806 | -83.678 | YELLOW | CENTRAL | Stanton, KY | *P. virginiana* | 5-Aug-2015 |

| | | | | | | | |
|---|---|---|---|---|---|---|---|
| LL194 | 37.806 | -83.678 | YELLOW | CENTRAL | Stanton, KY | *P. virginiana* | 5-Aug-2015 |
| LL195 | 37.806 | -83.678 | YELLOW | CENTRAL | Stanton, KY | *P. virginiana* | 5-Aug-2015 |
| LL196 | 37.805 | -83.656 | YELLOW | CENTRAL | Stanton, KY | *P. virginiana* | 5-Aug-2015 |
| LL200 | 38.014 | -84.504 | YELLOW | CENTRAL | Lexington, KY | *P. virginiana* | 13-Aug-2015 |
| LL204 | 38.014 | -84.504 | YELLOW | CENTRAL | Lexington, KY | *P. virginiana* | 20-Aug-2015 |
| LL205 | 38.014 | -84.504 | YELLOW | CENTRAL | Lexington, KY | *P. virginiana* | 20-Aug-2015 |
| LL206 | 38.014 | -84.504 | YELLOW | CENTRAL | Lexington, KY | *P. virginiana* | 20-Aug-2015 |
| LL207 | 38.014 | -84.504 | YELLOW | CENTRAL | Lexington, KY | *P. virginiana* | 20-Aug-2015 |
| LL208 | 38.033 | -84.507 | YELLOW | CENTRAL | Lexington, KY | *P. mugho* | 24-Aug-2015 |
| LL209 | 38.014 | -84.504 | YELLOW | CENTRAL | Lexington, KY | *P. virginiana* | 28-Aug-2015 |
| LL210 | 38.014 | -84.504 | YELLOW | CENTRAL | Lexington, KY | *P. virginiana* | 28-Aug-2015 |
| LL211 | 38.014 | -84.504 | YELLOW | CENTRAL | Lexington, KY | *P. virginiana* | 28-Aug-2015 |
| LL212 | 38.014 | -84.504 | YELLOW | CENTRAL | Lexington, KY | *P. virginiana* | 28-Aug-2015 |
| LL213 | 38.014 | -84.504 | YELLOW | CENTRAL | Lexington, KY | *P. virginiana* | 28-Aug-2015 |
| LL214 | 38.014 | -84.504 | YELLOW | CENTRAL | Lexington, KY | *P. virginiana* | 28-Aug-2015 |
| LL215 | 38.014 | -84.504 | YELLOW | CENTRAL | Lexington, KY | *P. virginiana* | 28-Aug-2015 |
| LL216 | 38.014 | -84.504 | YELLOW | CENTRAL | Lexington, KY | *P. echinata* | 28-Aug-2015 |
| LL218 | 38.014 | -84.504 | YELLOW | CENTRAL | Lexington, KY | *P. virginiana* | 10-Sep-2015 |
| LL219 | 38.014 | -84.504 | YELLOW | CENTRAL | Lexington, KY | *P. virginiana* | 10-Sep-2015 |
| LL220 | 38.014 | -84.504 | YELLOW | CENTRAL | Lexington, KY | *P. virginiana* | 10-Sep-2015 |
| LL221 | 38.014 | -84.504 | YELLOW | CENTRAL | Lexington, KY | *P. virginiana* | 10-Sep-2015 |
| LL222 | 38.014 | -84.504 | YELLOW | CENTRAL | Lexington, KY | *P. virginiana* | 10-Sep-2015 |
| LL223 | 38.014 | -84.504 | YELLOW | CENTRAL | Lexington, KY | *P. virginiana* | 10-Sep-2015 |
| LL224 | 38.024 | -84.532 | YELLOW | CENTRAL | Lexington, KY | *P. mugho* | 12-Sep-2015 |
| LL225 | 38.014 | -84.504 | YELLOW | CENTRAL | Lexington, KY | *P. virginiana* | 14-Sep-2015 |
| LL226 | 38.014 | -84.504 | YELLOW | CENTRAL | Lexington, KY | *P. virginiana* | 14-Sep-2015 |
| LL227 | 38.014 | -84.504 | YELLOW | CENTRAL | Lexington, KY | *P. virginiana* | 14-Sep-2015 |
| LL228 | 38.014 | -84.504 | YELLOW | CENTRAL | Lexington, KY | *P. virginiana* | 14-Sep-2015 |
| LL229 | 38.014 | -84.504 | YELLOW | CENTRAL | Lexington, KY | *P. virginiana* | 14-Sep-2015 |

| | | | | | | | |
|---|---|---|---|---|---|---|---|
| LL230 | 38.014 | -84.504 | YELLOW | CENTRAL | Lexington, KY | *P. virginiana* | 14-Sep-2015 |
| LL242 | 37.984 | -84.418 | YELLOW | CENTRAL | Lexington, KY | *P. taeda* | 7-Jun-2016 |
| LL243 | 37.984 | -84.418 | YELLOW | CENTRAL | Lexington, KY | *P. taeda* | 7-Jun-2016 |
| LL245 | 38.014 | -84.504 | YELLOW | CENTRAL | Lexington, KY | *P. virginiana* | 9-Jun-2016 |
| LL246 | 37.984 | -84.418 | YELLOW | CENTRAL | Lexington, KY | *P. taeda* | 13-Jun-2016 |
| LL247 | 38.014 | -84.504 | YELLOW | CENTRAL | Lexington, KY | *P. virginiana* | 16-Jun-2016 |
| LL248 | 38.014 | -84.504 | YELLOW | CENTRAL | Lexington, KY | *P. virginiana* | 16-Jun-2016 |
| LL249 | 38.014 | -84.504 | YELLOW | CENTRAL | Lexington, KY | *P. virginiana* | 16-Jun-2016 |
| LL250 | 38.014 | -84.504 | YELLOW | CENTRAL | Lexington, KY | *P. virginiana* | 16-Jun-2016 |
| LL251 | 38.014 | -84.504 | YELLOW | CENTRAL | Lexington, KY | *P. virginiana* | 16-Jun-2016 |
| LL252 | 38.044 | -84.497 | YELLOW | CENTRAL | Lexington, KY | *P. mugho* | 23-Jun-2016 |
| LL253 | 38.044 | -84.497 | YELLOW | CENTRAL | Lexington, KY | *P. mugho* | 23-Jun-2016 |
| LL254 | 38.014 | -84.504 | YELLOW | CENTRAL | Lexington, KY | *P. rigida* | 24-Jun-2016 |
| LL255 | 38.014 | -84.504 | YELLOW | CENTRAL | Lexington, KY | *P. virginiana* | 24-Jun-2016 |
| LL256 | 38.014 | -84.504 | YELLOW | CENTRAL | Lexington, KY | *P. virginiana* | 24-Jun-2016 |
| LL257 | 38.014 | -84.504 | YELLOW | CENTRAL | Lexington, KY | *P. virginiana* | 24-Jun-2016 |
| LL258 | 38.014 | -84.504 | YELLOW | CENTRAL | Lexington, KY | *P. virginiana* | 24-Jun-2016 |
| LL259 | 38.014 | -84.504 | YELLOW | CENTRAL | Lexington, KY | *P. virginiana* | 24-Jun-2016 |
| LL260 | 38.014 | -84.504 | YELLOW | CENTRAL | Lexington, KY | *P. virginiana* | 24-Jun-2016 |
| LL261 | 38.044 | -84.497 | YELLOW | CENTRAL | Lexington, KY | *P. mugho* | 24-Jun-2016 |
| LL262 | 38.044 | -84.497 | YELLOW | CENTRAL | Lexington, KY | *P. mugho* | 24-Jun-2016 |
| LL263 | 38.044 | -84.497 | YELLOW | CENTRAL | Lexington, KY | *P. mugho* | 24-Jun-2016 |
| LL264 | 38.044 | -84.497 | YELLOW | CENTRAL | Lexington, KY | *P. mugho* | 24-Jun-2016 |
| LL265 | 38.044 | -84.497 | YELLOW | CENTRAL | Lexington, KY | *P. mugho* | 24-Jun-2016 |
| LL266 | 38.044 | -84.497 | YELLOW | CENTRAL | Lexington, KY | *P. mugho* | 24-Jun-2016 |
| LL267 | 38.044 | -84.497 | YELLOW | CENTRAL | Lexington, KY | *P. mugho* | 24-Jun-2016 |
| LL268 | 38.044 | -84.497 | YELLOW | CENTRAL | Lexington, KY | *P. mugho* | 24-Jun-2016 |
| LL269 | 38.044 | -84.497 | YELLOW | CENTRAL | Lexington, KY | *P. mugho* | 24-Jun-2016 |
| LL270 | 38.044 | -84.497 | YELLOW | CENTRAL | Lexington, KY | *P. mugho* | 24-Jun-2016 |

| | | | | | | | |
|---|---|---|---|---|---|---|---|
| LL271 | 38.044 | -84.497 | YELLOW | CENTRAL | Lexington, KY | *P. mugho* | 24-Jun-2016 |
| LL272 | 38.044 | -84.497 | YELLOW | CENTRAL | Lexington, KY | *P. mugho* | 24-Jun-2016 |
| LL273 | 38.044 | -84.497 | YELLOW | CENTRAL | Lexington, KY | *P. mugho* | 24-Jun-2016 |
| LL274 | 38.044 | -84.497 | YELLOW | CENTRAL | Lexington, KY | *P. mugho* | 24-Jun-2016 |
| LL275 | 38.044 | -84.497 | YELLOW | CENTRAL | Lexington, KY | *P. mugho* | 24-Jun-2016 |
| LL276 | 38.044 | -84.497 | YELLOW | CENTRAL | Lexington, KY | *P. mugho* | 24-Jun-2016 |
| LL277 | 35.980 | -85.015 | YELLOW | CENTRAL | Crossville, TN | *P. virginiana* | 25-Jun-2016 |
| LL278 | 35.980 | -85.015 | YELLOW | CENTRAL | Crossville, TN | *P. virginiana* | 25-Jun-2016 |
| LL279 | 38.014 | -84.504 | YELLOW | CENTRAL | Lexington, KY | *P. virginiana* | 24-Aug-2016 |
| LL280 | 38.014 | -84.504 | YELLOW | CENTRAL | Lexington, KY | *P. virginiana* | 24-Aug-2016 |
| LL281 | 38.014 | -84.504 | YELLOW | CENTRAL | Lexington, KY | *P. virginiana* | 24-Aug-2016 |
| LL282 | 38.014 | -84.504 | YELLOW | CENTRAL | Lexington, KY | *P. virginiana* | 24-Aug-2016 |
| LL283 | 38.014 | -84.504 | YELLOW | CENTRAL | Lexington, KY | *P. virginiana* | 24-Aug-2016 |
| LL284 | 38.014 | -84.504 | YELLOW | CENTRAL | Lexington, KY | *P. virginiana* | 24-Aug-2016 |
| LL285 | 38.014 | -84.504 | YELLOW | CENTRAL | Lexington, KY | *P. virginiana* | 24-Aug-2016 |
| LL286 | 38.014 | -84.504 | YELLOW | CENTRAL | Lexington, KY | *P. virginiana* | 24-Aug-2016 |
| LL287 | 38.014 | -84.504 | YELLOW | CENTRAL | Lexington, KY | *P. virginiana* | 24-Aug-2016 |
| LL288 | 38.014 | -84.504 | YELLOW | CENTRAL | Lexington, KY | *P. virginiana* | 24-Aug-2016 |
| LL289 | 38.014 | -84.504 | YELLOW | CENTRAL | Lexington, KY | *P. virginiana* | 2-Sep-2016 |
| LL290 | 38.014 | -84.504 | YELLOW | CENTRAL | Lexington, KY | *P. virginiana* | 2-Sep-2016 |
| LL291 | 38.014 | -84.504 | YELLOW | CENTRAL | Lexington, KY | *P. virginiana* | 2-Sep-2016 |
| LL292 | 38.014 | -84.504 | YELLOW | CENTRAL | Lexington, KY | *P. virginiana* | 2-Sep-2016 |
| LL293 | 38.014 | -84.504 | YELLOW | CENTRAL | Lexington, KY | *P. virginiana* | 2-Sep-2016 |
| LL294 | 38.014 | -84.504 | YELLOW | CENTRAL | Lexington, KY | *P. virginiana* | 2-Sep-2016 |
| LL295 | 38.014 | -84.504 | YELLOW | CENTRAL | Lexington, KY | *P. virginiana* | 2-Sep-2016 |
| LL296 | 38.014 | -84.504 | YELLOW | CENTRAL | Lexington, KY | *P. virginiana* | 2-Sep-2016 |
| LL297 | 38.014 | -84.504 | YELLOW | CENTRAL | Lexington, KY | *P. virginiana* | 2-Sep-2016 |
| LL298 | 38.014 | -84.504 | YELLOW | CENTRAL | Lexington, KY | *P. virginiana* | 2-Sep-2016 |
| LL299 | 38.014 | -84.504 | YELLOW | CENTRAL | Lexington, KY | *P. virginiana* | 2-Sep-2016 |

| | | | | | | | |
|---|---|---|---|---|---|---|---|
| LL300 | 38.014 | -84.504 | YELLOW | CENTRAL | Lexington, KY | *P. virginiana* | 2-Sep-2016 |
| LL301 | 38.014 | -84.504 | YELLOW | CENTRAL | Lexington, KY | *P. virginiana* | 2-Sep-2016 |
| LL302 | 38.014 | -84.504 | YELLOW | CENTRAL | Lexington, KY | *P. virginiana* | 2-Sep-2016 |
| LL303 | 38.014 | -84.504 | YELLOW | CENTRAL | Lexington, KY | *P. virginiana* | 2-Sep-2016 |
| LL304 | 38.014 | -84.504 | YELLOW | CENTRAL | Lexington, KY | *P. virginiana* | 2-Sep-2016 |
| LL305 | 38.014 | -84.504 | YELLOW | CENTRAL | Lexington, KY | *P. virginiana* | 2-Sep-2016 |
| RB001 | 40.680 | -74.234 | YELLOW | CENTRAL | Union, NJ | *P. strobus* | 5-Sep-2009 |
| RB002 | 38.171 | -83.556 | YELLOW | CENTRAL | Morehead, KY | *P. rigida* | 25-Jul-2011 |
| RB003 | 40.603 | -74.475 | MIXED | CENTRAL | Greenbrook, NJ | *P. mugho* | 1-Jul-2011 |
| RB004 | 44.350 | -89.822 | YELLOW | NORTH | Grand Rapids, WI | *P. banksiana* | 28-Jul-2011 |
| RB006 | 37.997 | -84.672 | YELLOW | CENTRAL | London, KY | *P. echinata* | 2-Aug-2011 |
| RB008 | 32.138 | -82.969 | YELLOW | SOUTH | Helena, GA | *P. elliottii* | 4-Aug-2011 |
| RB009 | 32.523 | -83.496 | YELLOW | SOUTH | Dry Branch, GA | *P. taeda* | 7-Aug-2011 |
| RB010 | 32.523 | -83.496 | YELLOW | SOUTH | Dry Branch, GA | *P. echinata* | 7-Aug-2011 |
| RB011 | 32.523 | -83.496 | YELLOW | SOUTH | Dry Branch, GA | *P. echinata* | 7-Aug-2011 |
| RB012 | 32.523 | -83.496 | YELLOW | SOUTH | Dry Branch, GA | *P. echinata* | 7-Aug-2011 |
| RB015 | 44.461 | -85.992 | YELLOW | NORTH | Springdale Township, MI | *P. banksiana* | 4-Aug-2011 |
| RB016 | 37.066 | -84.159 | YELLOW | CENTRAL | Laurel Co, KY | *P. echinata* | 17-Aug-2011 |
| RB017 | 37.984 | -84.511 | YELLOW | CENTRAL | Lexingon, KY | *P. mugho* | 18-Aug-2011 |
| RB018 | 43.797 | -71.915 | YELLOW | CENTRAL | Dorchester, NH | *P. resinosa* | 5-Aug-2011 |
| RB019 | 37.066 | -84.159 | YELLOW | CENTRAL | Laurel Co, KY | *P. echinata* | 6-Sep-2011 |
| RB020 | 37.066 | -84.159 | YELLOW | CENTRAL | Laurel Co, KY | *P. echinata* | 6-Sep-2011 |
| RB021 | 37.066 | -84.159 | YELLOW | CENTRAL | Laurel Co, KY | *P. echinata* | 6-Sep-2011 |
| RB022 | 38.024 | -84.494 | YELLOW | CENTRAL | Lexington, KY | *P. nigra* | 7-Sep-2011 |
| RB024 | 44.936 | -75.668 | YELLOW | NORTH | Pattersons Corners, ON | *young pinus resinosa* | 31-Aug-2011 |
| RB025 | 41.268 | -78.280 | MIXED | CENTRAL | Jay Township, PN | *P. mugho* | 17-Oct-2011 |
| RB027 | 33.990 | -83.796 | YELLOW | CENTRAL/SOUTH | Auburn, GA | *P. echinata* | 25-Oct-2011 |
| RB028 | 39.690 | -74.593 | WHITE | CENTRAL | Washington, NJ | *P. rigida* | 9-Sep-2010 |

| | | | | | | | |
|---|---|---|---|---|---|---|---|
| RB029 | 38.209 | -84.390 | YELLOW | CENTRAL | Scotch Plains, NJ | *P. sylvestris* | 9/2010 |
| RB039 | 37.071 | -84.211 | YELLOW | CENTRAL | London, KY | *P. echinata* | 29-May-2012 |
| RB040 | 38.209 | -84.390 | YELLOW | CENTRAL | Lexington, KY | *P. sylvestris* | 29-May-2012 |
| RB042 | 38.024 | -84.494 | YELLOW | CENTRAL | Lexington, KY | *P. nigra* | 30-May-2012 |
| RB044 | 27.692 | -82.420 | YELLOW | SOUTH | Ruskin, FL | *P. elliottii* | 5-Jun-2012 |
| RB046 | 27.618 | -81.815 | YELLOW | SOUTH | Bowling Green, FL | *P. palustris* | 8-Jun-2012 |
| RB047 | 27.618 | -81.815 | YELLOW | SOUTH | Bowling Green, FL | *P. palustris* | 8-Jun-2012 |
| RB048 | 27.618 | -81.815 | YELLOW | SOUTH | Bowling Green, FL | *P. palustris* | 8-Jun-2012 |
| RB049 | 29.507 | -81.860 | YELLOW | SOUTH | Interlachen, FL | *P. palustris* | 9-Jun-2012 |
| RB050 | 29.507 | -81.860 | YELLOW | SOUTH | Interlachen, FL | *P. palustris* | 9-Jun-2012 |
| RB051 | 29.507 | -81.860 | YELLOW | SOUTH | Interlachen, FL | *P. palustris* | 9-Jun-2012 |
| RB052 | 29.507 | -81.860 | YELLOW | SOUTH | Interlachen, FL | *P. palustris* | 9-Jun-2012 |
| RB053 | 29.507 | -81.860 | YELLOW | SOUTH | Interlachen, FL | *P. palustris* | 9-Jun-2012 |
| RB054 | 29.507 | -81.860 | YELLOW | SOUTH | Interlachen, FL | *P. palustris* | 9-Jun-2012 |
| RB055 | 29.507 | -81.860 | YELLOW | SOUTH | Interlachen, FL | *P. palustris* | 9-Jun-2012 |
| RB056 | 29.507 | -81.860 | YELLOW | SOUTH | Interlachen, FL | *P. palustris* | 9-Jun-2012 |
| RB057 | 29.320 | -81.727 | YELLOW | SOUTH | Salt Springs, FL | *P. palustris* | 10-Jun-2012 |
| RB058 | 29.320 | -81.727 | YELLOW | SOUTH | Salt Springs, FL | *P. palustris* | 10-Jun-2012 |
| RB059 | 29.320 | -81.727 | YELLOW | SOUTH | Salt Springs, FL | *P. palustris* | 10-Jun-2012 |
| RB060 | 29.320 | -81.727 | YELLOW | SOUTH | Salt Springs, FL | *P. palustris* | 10-Jun-2012 |
| RB061 | 29.320 | -81.727 | YELLOW | SOUTH | Salt Springs, FL | *P. palustris* | 10-Jun-2012 |
| RB062 | 29.320 | -81.727 | YELLOW | SOUTH | Salt Springs, FL | *P. palustris* | 10-Jun-2012 |
| RB063 | 29.320 | -81.727 | YELLOW | SOUTH | Salt Springs, FL | *P. palustris* | 10-Jun-2012 |
| RB064 | 29.507 | -82.960 | YELLOW | SOUTH | Fanning Springs, FL | *P. palustris* | 11-Jun-2012 |
| RB065 | 29.507 | -82.960 | YELLOW | SOUTH | Fanning Springs, FL | *P. palustris* | 11-Jun-2012 |
| RB066 | 29.508 | -82.958 | YELLOW | SOUTH | Fanning Springs, FL | *P. palustris* | 11-Jun-2012 |
| RB067 | 29.508 | -82.958 | YELLOW | SOUTH | Fanning Springs, FL | *P. palustris* | 11-Jun-2012 |
| RB068 | 30.191 | -84.370 | YELLOW | SOUTH | Crawfordville, FL | *P. palustris* | 12-Jun-2012 |
| RB069 | 30.191 | -84.370 | YELLOW | SOUTH | Crawfordville, FL | *P. palustris* | 12-Jun-2012 |

| | | | | | | | |
|---|---|---|---|---|---|---|---|
| RB071 | 32.843 | -87.952 | YELLOW | CENTRAL/SOUTH | Eutaw, AL | *P. echinata* | 16-Jun-2012 |
| RB073 | 38.014 | -84.504 | YELLOW | CENTRAL | Lexington, KY | *P. rigida* | 21-Jun-2012 |
| RB074 | 37.984 | -84.418 | YELLOW | CENTRAL | Lexington, KY | *P. taeda* | 27-Jun-2012 |
| RB075 | 32.239 | -80.859 | YELLOW | SOUTH | Bluffton, SC | *P. palustris* | 25-Jun-2012 |
| RB076 | 38.014 | -84.504 | YELLOW | CENTRAL | Lexington, KY | *P. virginiana* | 5-Jul-2012 |
| RB077 | 43.759 | -85.741 | YELLOW | NORTH | Brohman, MI | *P. banksiana* | 20-Jul-2012 |
| RB078 | 43.759 | -85.741 | YELLOW | NORTH | Brohman, MI | *P. banksiana* | 20-Jul-2012 |
| RB079 | 43.759 | -85.741 | YELLOW | NORTH | Brohman, MI | *P. banksiana* | 20-Jul-2012 |
| RB080 | 43.796 | -85.740 | YELLOW | NORTH | Bitely, MI | *P. banksiana* | 20-Jul-2012 |
| RB081 | 43.796 | -85.740 | YELLOW | NORTH | Bitely, MI | *P. banksiana* | 20-Jul-2012 |
| RB082 | 43.796 | -85.740 | YELLOW | NORTH | Bitely, MI | *P. banksiana* | 20-Jul-2012 |
| RB083 | 43.796 | -85.740 | YELLOW | NORTH | Bitely, MI | *P. banksiana* | 20-Jul-2012 |
| RB084 | 43.796 | -85.740 | YELLOW | NORTH | Bitely, MI | *P. banksiana* | 20-Jul-2012 |
| RB085 | 43.796 | -85.740 | YELLOW | NORTH | Bitely, MI | *P. banksiana* | 20-Jul-2012 |
| RB086 | 43.796 | -85.740 | YELLOW | NORTH | Bitely, MI | *P. banksiana* | 20-Jul-2012 |
| RB087 | 43.796 | -85.740 | YELLOW | NORTH | Bitely, MI | *P. banksiana* | 20-Jul-2012 |
| RB088 | 43.796 | -85.740 | YELLOW | NORTH | Bitely, MI | *P. banksiana* | 20-Jul-2012 |
| RB089 | 43.796 | -85.740 | YELLOW | NORTH | Bitely, MI | *P. banksiana* | 20-Jul-2012 |
| RB090 | 44.657 | -84.414 | YELLOW | NORTH | Grayling, MI | *P. sylvestris* | 21-Jul-2012 |
| RB091 | 44.657 | -84.414 | YELLOW | NORTH | Grayling, MI | *P. banksiana* | 21-Jul-2012 |
| RB092 | 44.657 | -84.414 | YELLOW | NORTH | Grayling, MI | *P. banksiana* | 21-Jul-2012 |
| RB093 | 44.657 | -84.414 | YELLOW | NORTH | Grayling, MI | *P. banksiana* | 21-Jul-2012 |
| RB094 | 45.504 | -84.615 | YELLOW | NORTH | Glaque Beach, MI | *P. banksiana* | 21-Jul-2012 |
| RB095 | 46.094 | -85.339 | YELLOW | NORTH | Naubinway, MI | *P. banksiana* | 22-Jul-2012 |
| RB096 | 46.096 | -85.394 | YELLOW | NORTH | Naubinway, MI | *P. banksiana* | 22-Jul-2012 |
| RB097 | 46.096 | -85.394 | YELLOW | NORTH | Naubinway, MI | *P. banksiana* | 22-Jul-2012 |
| RB098 | 46.096 | -85.394 | YELLOW | NORTH | Naubinway, MI | *P. banksiana* | 22-Jul-2012 |
| RB099 | 45.924 | -86.303 | YELLOW | NORTH | Manistique, MI | *P. banksiana* | 22-Jul-2012 |
| RB100 | 46.354 | -89.179 | YELLOW | NORTH | Bruce Crossing, MI | *P. banksiana* | 23-Jul-2012 |

| | | | | | | | |
|---|---|---|---|---|---|---|---|
| RB101 | 44.985 | -88.449 | YELLOW | NORTH | How, WI | *P. resinosa* | 24-Jul-2012 |
| RB102 | 44.985 | -88.449 | YELLOW | NORTH | How, WI | *P. resinosa* | 24-Jul-2012 |
| RB103 | 44.985 | -88.449 | YELLOW | NORTH | How, WI | *P. resinosa* | 24-Jul-2012 |
| RB104 | 44.985 | -88.449 | YELLOW | NORTH | How, WI | *P. resinosa* | 24-Jul-2012 |
| RB105 | 44.985 | -88.449 | YELLOW | NORTH | How, WI | *P. resinosa* | 24-Jul-2012 |
| RB106 | 37.913 | -79.896 | WHITE | CENTRAL | Valley Springs, VA | *P. virginiana* | 2-Aug-2012 |
| RB107 | 38.212 | -79.719 | WHITE | CENTRAL | Mountain Grove, VA | *P. rigida* | 2-Aug-2012 |
| RB108 | 38.678 | -79.399 | WHITE | CENTRAL | Deer Run, WV | *P. rigida* | 2-Aug-2012 |
| RB110 | 39.934 | -74.533 | WHITE | CENTRAL | Browns Mills, NJ | *P. rigida* | 5-Aug-2012 |
| RB112 | 39.621 | -74.428 | WHITE | CENTRAL | Tuckerton, NJ | *P. rigida* | 6-Aug-2012 |
| RB118 | 38.592 | -79.172 | WHITE | CENTRAL | Eastern, WV | *P. virginiana* | 9-Aug-2012 |
| RB119 | 37.713 | -79.367 | WHITE | CENTRAL | Buena Vista, VA | *P. virginiana* | 10-Aug-2012 |
| RB120 | 38.209 | -84.390 | YELLOW | CENTRAL | Lexington, KY | *P. sylvestris* | 20-Aug-2012 |
| RB121 | 38.209 | -84.390 | YELLOW | CENTRAL | Lexington, KY | *P. sylvestris* | 20-Aug-2012 |
| RB122 | 38.010 | -84.302 | YELLOW | CENTRAL | Lexington, KY | *P. echinata* | 20-Aug-2012 |
| RB124 | 38.209 | -84.390 | YELLOW | CENTRAL | Scotch Plains, NJ | *P. sylvestris* | 26-Aug-2012 |
| RB125 | 39.717 | -78.280 | YELLOW | CENTRAL | Sideling Hill, MD | *P. mugho* | 31-Aug-2012 |
| RB126 | 38.014 | -84.504 | YELLOW | CENTRAL | Lexington, KY | *P. virginiana* | 31-Aug-2012 |
| RB127 | 38.014 | -84.504 | YELLOW | CENTRAL | Lexington, KY | *P. virginiana* | 31-Aug-2012 |
| RB128 | 38.014 | -84.504 | YELLOW | CENTRAL | Lexington, KY | *P. virginiana* | 31-Aug-2012 |
| RB129 | 38.014 | -84.504 | YELLOW | CENTRAL | Lexington, KY | *P. echinata* | 31-Aug-2012 |
| RB130 | 32.277 | -80.983 | YELLOW | SOUTH | Bluffton SC | *P. palustris* | 30-Aug-2012 |
| RB131 | 32.277 | -80.983 | YELLOW | SOUTH | Bluffton SC | *P. taeda* | 30-Aug-2012 |
| RB132 | 38.014 | -84.504 | YELLOW | CENTRAL | Lexington, KY | *P. rigida* | 14-Sep-2012 |
| RB133 | 38.014 | -84.504 | YELLOW | CENTRAL | Lexington, KY | *P. virginiana* | 14-Sep-2012 |
| RB134 | 38.014 | -84.504 | YELLOW | CENTRAL | Lexington, KY | *P. virginiana* | 14-Sep-2012 |
| RB135 | 38.014 | -84.504 | YELLOW | CENTRAL | Lexington, KY | *P. virginiana* | 14-Sep-2012 |
| RB136 | 43.759 | -85.741 | YELLOW | NORTH | Bitely, MI | *P. resinosa* | 23-Oct-2012 |
| RB137 | 27.618 | -81.815 | YELLOW | SOUTH | Bowling Green, FL | *P. elliottii* | 23-Oct-2012 |

| | | | | | | | |
|---|---|---|---|---|---|---|---|
| RB138 | 27.618 | -81.815 | YELLOW | SOUTH | Bowling Green, FL | *P. elliottii* | 23-Oct-2012 |
| RB139 | 27.618 | -81.815 | YELLOW | SOUTH | Bowling Green, FL | *P. elliottii* | 23-Oct-2012 |
| RB141 | 38.014 | -84.504 | MIXED | CENTRAL | Lexington, KY | *P. virginiana* | 26-Jun-2013 |
| RB142 | 38.209 | -84.390 | YELLOW | CENTRAL | Lexington, KY | *P. sylvestris* | 26-Jun-2013 |
| RB143 | 38.209 | -84.390 | YELLOW | CENTRAL | Lexington, KY | *P. sylvestris* | 26-Jun-2013 |
| RB144 | 38.209 | -84.390 | YELLOW | CENTRAL | Lexington, KY | *P. sylvestris* | 26-Jun-2013 |
| RB145 | 38.209 | -84.390 | YELLOW | CENTRAL | Lexington, KY | *P. sylvestris* | 26-Jun-2013 |
| RB146 | 38.209 | -84.390 | YELLOW | CENTRAL | Lexington, KY | *P. sylvestris* | 26-Jun-2013 |
| RB147 | 38.209 | -84.390 | YELLOW | CENTRAL | Lexington, KY | *P. sylvestris* | 26-Jun-2013 |
| RB148 | 38.209 | -84.390 | YELLOW | CENTRAL | Lexington, KY | *P. sylvestris* | 26-Jun-2013 |
| RB149 | 38.209 | -84.390 | YELLOW | CENTRAL | Lexington, KY | *P. sylvestris* | 26-Jun-2013 |
| RB150 | 38.209 | -84.390 | YELLOW | CENTRAL | Lexington, KY | *P. sylvestris* | 26-Jun-2013 |
| RB151 | 38.209 | -84.390 | YELLOW | CENTRAL | Lexington, KY | *P. sylvestris* | 26-Jun-2013 |
| RB152 | 38.209 | -84.390 | YELLOW | CENTRAL | Lexington, KY | *P. sylvestris* | 26-Jun-2013 |
| RB153 | 38.209 | -84.390 | YELLOW | CENTRAL | Lexington, KY | *P. sylvestris* | 26-Jun-2013 |
| RB154 | 38.209 | -84.390 | YELLOW | CENTRAL | Lexington, KY | *P. sylvestris* | 26-Jun-2013 |
| RB155 | 38.209 | -84.390 | YELLOW | CENTRAL | Lexington, KY | *P. sylvestris* | 26-Jun-2013 |
| RB156 | 38.209 | -84.390 | YELLOW | CENTRAL | Lexington, KY | *P. sylvestris* | 26-Jun-2013 |
| RB157 | 38.209 | -84.390 | YELLOW | CENTRAL | Lexington, KY | *P. sylvestris* | 26-Jun-2013 |
| RB159 | 37.984 | -84.418 | YELLOW | CENTRAL | Lexington, KY | *P. taeda* | 27-Jun-2013 |
| RB160 | 37.984 | -84.418 | YELLOW | CENTRAL | Lexington, KY | *P. taeda* | 27-Jun-2013 |
| RB161 | 37.984 | -84.418 | YELLOW | CENTRAL | Lexington, KY | *P. taeda* | 27-Jun-2013 |
| RB162 | 38.024 | -84.494 | YELLOW | CENTRAL | Lexington, KY | *P. nigra* | 2-Jul-2013 |
| RB163 | 35.280 | -82.117 | YELLOW | NORTH | Tryon, NC | *P. taeda* | 3-Jul-2013 |
| RB164 | 35.280 | -82.117 | YELLOW | NORTH | Tryon, NC | *P. virginiana* | 3-Jul-2013 |
| RB165 | 35.280 | -82.118 | YELLOW | NORTH | Tryon, NC | *P. virginiana* | 3-Jul-2013 |
| RB167 | 35.183 | -81.963 | YELLOW | SOUTH | Chesnee,SC | *P. virginiana* | 4-Jul-2013 |
| RB168 | 35.183 | -81.963 | YELLOW | SOUTH | Chesnee,SC | *P. virginiana* | 4-Jul-2013 |
| RB171 | 35.183 | -81.963 | YELLOW | SOUTH | Chesnee,SC | *P. taeda* | 4-Jul-2013 |

| | | | | | | | |
|---|---|---|---|---|---|---|---|
| RB172 | 35.183 | -81.963 | YELLOW | SOUTH | Chesnee,SC | *P. taeda* | 4-Jul-2013 |
| RB173 | 35.183 | -81.963 | YELLOW | SOUTH | Chesnee,SC | *P. taeda* | 4-Jul-2013 |
| RB174 | 35.183 | -81.963 | YELLOW | SOUTH | Chesnee,SC | *P. taeda* | 4-Jul-2013 |
| RB175 | 35.183 | -81.963 | YELLOW | SOUTH | Chesnee,SC | *P. taeda* | 4-Jul-2013 |
| RB176 | 35.183 | -81.963 | YELLOW | SOUTH | Chesnee,SC | *P. taeda* | 4-Jul-2013 |
| RB177 | 35.183 | -81.963 | YELLOW | SOUTH | Chesnee,SC | *P. taeda* | 4-Jul-2013 |
| RB178 | 35.183 | -81.963 | YELLOW | SOUTH | Chesnee,SC | *P. taeda* | 4-Jul-2013 |
| RB179 | 35.183 | -81.963 | YELLOW | SOUTH | Chesnee,SC | *P. taeda* | 4-Jul-2013 |
| RB180 | 35.183 | -81.963 | YELLOW | SOUTH | Chesnee,SC | *P. taeda* | 4-Jul-2013 |
| RB181 | 35.183 | -81.963 | YELLOW | SOUTH | Chesnee,SC | *P. taeda* | 4-Jul-2013 |
| RB182 | 35.183 | -81.963 | YELLOW | SOUTH | Chesnee,SC | *P. taeda* | 4-Jul-2013 |
| RB183 | 35.183 | -81.963 | YELLOW | SOUTH | Chesnee,SC | *P. taeda* | 4-Jul-2013 |
| RB184 | 35.183 | -81.963 | YELLOW | SOUTH | Chesnee,SC | *P. taeda* | 4-Jul-2013 |
| RB185 | 35.183 | -81.963 | YELLOW | SOUTH | Chesnee,SC | *P. taeda* | 4-Jul-2013 |
| RB186 | 35.183 | -81.963 | YELLOW | SOUTH | Chesnee,SC | *P. taeda* | 4-Jul-2013 |
| RB187 | 35.183 | -81.963 | YELLOW | SOUTH | Chesnee,SC | *P. taeda* | 4-Jul-2013 |
| RB188 | 35.183 | -81.963 | YELLOW | SOUTH | Chesnee,SC | *P. taeda* | 4-Jul-2013 |
| RB189 | 35.183 | -81.963 | YELLOW | SOUTH | Chesnee,SC | *P. taeda* | 4-Jul-2013 |
| RB190 | 35.014 | -82.716 | YELLOW | SOUTH | Pickens, SC | *P. virginiana* | 4-Jul-2013 |
| RB197 | 34.409 | -81.399 | YELLOW | SOUTH | Pomaria, SC | *P. taeda* | 5-Jul-2013 |
| RB198 | 34.409 | -81.399 | YELLOW | SOUTH | Pomaria, SC | *P. taeda* | 5-Jul-2013 |
| RB199 | 34.409 | -81.399 | YELLOW | SOUTH | Pomaria, SC | *P. taeda* | 5-Jul-2013 |
| RB200 | 34.396 | -81.402 | YELLOW | SOUTH | Blair, SC | *P. taeda* | 5-Jul-2013 |
| RB201 | 34.396 | -81.402 | YELLOW | SOUTH | Blair, SC | *P. taeda* | 5-Jul-2013 |
| RB202 | 34.396 | -81.402 | YELLOW | SOUTH | Blair, SC | *P. taeda* | 5-Jul-2013 |
| RB204 | 33.765 | -80.920 | YELLOW | SOUTH | Columbia, SC | *P. taeda* | 6-Jul-2013 |
| RB205 | 33.765 | -80.920 | YELLOW | SOUTH | Columbia, SC | *P. taeda* | 6-Jul-2013 |
| RB211 | 34.018 | -78.949 | YELLOW | SOUTH | Loris, SC | *P. taeda (?)* | 6-Jul-2013 |
| RB212 | 34.034 | -78.924 | YELLOW | SOUTH | Loris, SC | *P. taeda* | 6-Jul-2013 |

| | | | | | | | |
|---|---|---|---|---|---|---|---|
| RB214 | 34.527 | -78.745 | YELLOW | NORTH | Bladenboro, NC | *P. taeda* | 7-Jul-2013 |
| RB215 | 35.184 | -79.717 | YELLOW | NORTH | Norman, NC | *P. taeda* | 7-Jul-2013 |
| RB217 | 35.515 | -79.779 | YELLOW | NORTH | Seagrove, NC | *P. taeda* | 7-Jul-2013 |
| RB220 | 36.497 | -80.104 | YELLOW | NORTH | Sandy Ridge, NC | *P. virginiana* | 8-Jul-2013 |
| RB221 | 36.497 | -80.104 | YELLOW | NORTH | Sandy Ridge, NC | *P. virginiana* | 8-Jul-2013 |
| RB222 | 36.497 | -80.104 | YELLOW | NORTH | Sandy Ridge, NC | *P. virginiana* | 8-Jul-2013 |
| RB223 | 36.802 | -79.937 | WHITE | CENTRAL | Bassett, VA | *P. echinata* | 8-Jul-2013 |
| RB224 | 36.973 | -79.608 | WHITE | CENTRAL | Penhook, VA | *P. rigida* | 8-Jul-2013 |
| RB225 | 36.973 | -79.608 | WHITE | CENTRAL | Penhook, VA | *P. rigida* | 8-Jul-2013 |
| RB226 | 36.940 | -79.289 | MIXED | CENTRAL | Gretna, VA | *P. rigida* | 8-Jul-2013 |
| RB227 | 37.319 | -78.043 | WHITE | CENTRAL | Amelia, VA | *P. rigida* | 8-Jul-2013 |
| RB228 | 37.319 | -78.043 | WHITE | CENTRAL | Amelia, VA | *P. rigida* | 8-Jul-2013 |
| RB229 | 37.319 | -78.043 | WHITE | CENTRAL | Amelia, VA | *P. rigida* | 8-Jul-2013 |
| RB230 | 37.356 | -77.862 | MIXED | CENTRAL | Amelia, VA | *P. rigida* | 8-Jul-2013 |
| RB233 | 43.759 | -85.741 | YELLOW | NORTH | Bitely, MI | *P. banksiana* | 16-Jul-2013 |
| RB234 | 43.759 | -85.741 | YELLOW | NORTH | Bitely, MI | *P. banksiana* | 16-Jul-2013 |
| RB235 | 43.759 | -85.741 | YELLOW | NORTH | Bitely, MI | *P. banksiana* | 16-Jul-2013 |
| RB236 | 43.759 | -85.741 | YELLOW | NORTH | Bitely, MI | *P. banksiana* | 16-Jul-2013 |
| RB237 | 43.786 | -85.741 | YELLOW | NORTH | Bitely, MI | *P. banksiana* | 16-Jul-2013 |
| RB238 | 43.791 | -85.740 | YELLOW | NORTH | Bitely, MI | *P. banksiana* | 16-Jul-2013 |
| RB239 | 43.791 | -85.740 | YELLOW | NORTH | Bitely, MI | *P. banksiana* | 16-Jul-2013 |
| RB240 | 43.793 | -85.740 | YELLOW | NORTH | Bitely, MI | *P. banksiana* | 16-Jul-2013 |
| RB241 | 43.793 | -85.740 | YELLOW | NORTH | Bitely, MI | *P. banksiana* | 16-Jul-2013 |
| RB242 | 43.793 | -85.740 | YELLOW | NORTH | Bitely, MI | *P. banksiana* | 16-Jul-2013 |
| RB243 | 43.793 | -85.740 | YELLOW | NORTH | Bitely, MI | *P. banksiana* | 16-Jul-2013 |
| RB244 | 43.793 | -85.740 | YELLOW | NORTH | Bitely, MI | *P. banksiana* | 16-Jul-2013 |
| RB245 | 44.600 | -84.713 | YELLOW | NORTH | Grayling, MI | *P. banksiana* | 16-Jul-2013 |
| RB246 | 44.600 | -84.713 | YELLOW | NORTH | Grayling, MI | *P. banksiana* | 16-Jul-2013 |
| RB247 | 44.600 | -84.713 | YELLOW | NORTH | Grayling, MI | *P. banksiana* | 16-Jul-2013 |

| | | | | | | | |
|---|---|---|---|---|---|---|---|
| RB248 | 44.583 | -84.700 | YELLOW | NORTH | Grayling, MI | *P. banksiana* | 17-Jul-2013 |
| RB249 | 44.658 | -84.695 | YELLOW | NORTH | Grayling, MI | *P. resinosa* | 17-Jul-2013 |
| RB250 | 44.658 | -84.695 | YELLOW | NORTH | Grayling, MI | *P. resinosa* | 17-Jul-2013 |
| RB251 | 44.658 | -84.695 | YELLOW | NORTH | Grayling, MI | *P. resinosa* | 17-Jul-2013 |
| RB252 | 44.658 | -84.695 | YELLOW | NORTH | Grayling, MI | *P. resinosa* | 17-Jul-2013 |
| RB253 | 44.658 | -84.695 | YELLOW | NORTH | Grayling, MI | *P. resinosa* | 17-Jul-2013 |
| RB254 | 44.658 | -84.695 | YELLOW | NORTH | Grayling, MI | *P. banksiana* | 17-Jul-2013 |
| RB255 | 44.658 | -84.695 | YELLOW | NORTH | Grayling, MI | *P. sylvestris* | 17-Jul-2013 |
| RB256 | 44.658 | -84.695 | YELLOW | NORTH | Grayling, MI | *P. banksiana* | 17-Jul-2013 |
| RB257 | 44.658 | -84.695 | YELLOW | NORTH | Grayling, MI | *P. banksiana* | 17-Jul-2013 |
| RB258 | 44.658 | -84.695 | YELLOW | NORTH | Grayling, MI | *P. banksiana* | 17-Jul-2013 |
| RB259 | 44.657 | -84.696 | YELLOW | NORTH | Grayling, MI | *P. banksiana* | 17-Jul-2013 |
| RB260 | 44.657 | -84.696 | YELLOW | NORTH | Grayling, MI | *P. banksiana* | 17-Jul-2013 |
| RB261 | 44.657 | -84.696 | YELLOW | NORTH | Grayling, MI | *P. banksiana* | 17-Jul-2013 |
| RB266 | 44.985 | -88.449 | YELLOW | NORTH | Suring, WI | *P. resinosa* | 19-Jul-2013 |
| RB267 | 44.985 | -88.449 | YELLOW | NORTH | Suring, WI | *P. resinosa* | 19-Jul-2013 |
| RB268 | 44.985 | -88.449 | YELLOW | NORTH | Suring, WI | *P. resinosa* | 19-Jul-2013 |
| RB269 | 44.985 | -88.449 | YELLOW | NORTH | Suring, WI | *P. resinosa* | 19-Jul-2013 |
| RB270 | 44.985 | -88.449 | YELLOW | NORTH | Suring, WI | *P. resinosa* | 19-Jul-2013 |
| RB271 | 44.985 | -88.449 | YELLOW | NORTH | Suring, WI | *P. resinosa* | 19-Jul-2013 |
| RB272 | 44.985 | -88.449 | YELLOW | NORTH | Suring, WI | *P. resinosa* | 19-Jul-2013 |
| RB273 | 44.985 | -88.449 | YELLOW | NORTH | Suring, WI | *P. resinosa* | 19-Jul-2013 |
| RB274 | 44.985 | -88.449 | YELLOW | NORTH | Suring, WI | *P. resinosa* | 19-Jul-2013 |
| RB275 | 44.054 | -89.806 | YELLOW | NORTH | Arkdale, WI | *P. banksiana* | 20-Jul-2013 |
| RB278 | 44.112 | -90.117 | YELLOW | NORTH | Necedah, WI | *P. banksiana* | 20-Jul-2013 |
| RB279 | 44.115 | -90.118 | YELLOW | NORTH | Necedah, WI | *P. banksiana* | 20-Jul-2013 |
| RB280 | 44.115 | -90.118 | YELLOW | NORTH | Necedah, WI | *P. banksiana* | 20-Jul-2013 |
| RB281 | 44.115 | -90.118 | YELLOW | NORTH | Necedah, WI | *P. banksiana* | 20-Jul-2013 |
| RB282 | 44.115 | -90.118 | YELLOW | NORTH | Necedah, WI | *P. banksiana* | 20-Jul-2013 |

| | | | | | | | |
|---|---|---|---|---|---|---|---|
| RB283 | 44.136 | -90.128 | YELLOW | NORTH | Necedah, WI | *P. banksiana* | 20-Jul-2013 |
| RB284 | 44.207 | -90.136 | YELLOW | NORTH | Necedah, WI | *P. banksiana* | 20-Jul-2013 |
| RB285 | 44.207 | -90.136 | YELLOW | NORTH | Necedah, WI | *P. banksiana* | 20-Jul-2013 |
| RB286 | 39.631 | -77.963 | YELLOW | CENTRAL | Clear Spring, MD | *P. virginiana* | 25-Jul-2013 |
| RB287 | 39.631 | -77.963 | YELLOW | CENTRAL | Clear Spring, MD | *P. virginiana* | 25-Jul-2013 |
| RB288 | 39.631 | -77.963 | YELLOW | CENTRAL | Clear Spring, MD | *P. virginiana* | 25-Jul-2013 |
| RB289 | 39.631 | -77.963 | YELLOW | CENTRAL | Clear Spring, MD | *P. virginiana* | 25-Jul-2013 |
| RB290 | 40.387 | -74.335 | WHITE | CENTRAL | Old Bridge, NJ | *P. rigida* | 26-Jul-2013 |
| RB291 | 40.427 | -74.303 | MIXED | CENTRAL | Old Bridge NJ | *P. sylvestris* | 26-Jul-2013 |
| RB292 | 40.427 | -74.303 | WHITE | CENTRAL | Old Bridge, NJ | *P. sylvestris* | 26-Jul-2013 |
| RB293 | 43.559 | -73.733 | YELLOW | CENTRAL/NORTH | Warrensburg, NY | *P. sylvestris* | 28-Jul-2013 |
| RB294 | 43.559 | -73.733 | YELLOW | CENTRAL/NORTH | Warrensburg, NY | *P. sylvestris* | 28-Jul-2013 |
| RB295 | 43.559 | -73.733 | YELLOW | CENTRAL/NORTH | Warrensburg, NY | *P. sylvestris* | 28-Jul-2013 |
| RB296 | 43.559 | -73.733 | YELLOW | CENTRAL/NORTH | Warrensburg, NY | *P. sylvestris* | 28-Jul-2013 |
| RB297 | 43.559 | -73.733 | YELLOW | CENTRAL/NORTH | Warrensburg, NY | *P. sylvestris* | 28-Jul-2013 |
| RB298 | 43.559 | -73.733 | YELLOW | CENTRAL/NORTH | Warrensburg, NY | *P. strobus* | 28-Jul-2013 |
| RB299 | 44.000 | -73.718 | YELLOW | CENTRAL/NORTH | Warrensburg, NY | *P. strobus* | 28-Jul-2013 |
| RB300 | 44.000 | -73.718 | YELLOW | CENTRAL/NORTH | Warrensburg, NY | *P. strobus* | 28-Jul-2013 |
| RB301 | 44.000 | -73.718 | YELLOW | CENTRAL/NORTH | Warrensburg, NY | *P. resinosa* | 28-Jul-2013 |
| RB302 | 44.000 | -73.718 | YELLOW | CENTRAL/NORTH | Warrensburg, NY | *P. resinosa* | 28-Jul-2013 |
| RB303 | 44.016 | -73.704 | YELLOW | CENTRAL/NORTH | North Hudson, NY | *P. resinosa* | 28-Jul-2013 |
| RB304 | 43.819 | -71.205 | YELLOW | CENTRAL | Ossippee, NH | *P. rigida* | 29-Jul-2013 |
| RB305 | 43.819 | -71.205 | YELLOW | CENTRAL | Ossippee, NH | *P. rigida* | 29-Jul-2013 |
| RB306 | 43.819 | -71.205 | YELLOW | CENTRAL | Ossippee, NH | *P. rigida* | 29-Jul-2013 |
| RB307 | 43.819 | -71.205 | YELLOW | CENTRAL | Ossippee, NH | *P. rigida* | 29-Jul-2013 |
| RB308 | 43.676 | -71.082 | YELLOW | CENTRAL | Ossippee, NH | *P. rigida* | 29-Jul-2013 |
| RB309 | 30.239 | -82.299 | YELLOW | SOUTH | Sanderson, FL | *P. ellotti* | 6-Aug-2013 |
| RB310 | 29.508 | -81.860 | YELLOW | SOUTH | Fort McCoy, FL | *P. palustris* | 7-Aug-2013 |
| RB311 | 29.508 | -81.860 | YELLOW | SOUTH | Fort McCoy, FL | *P. palustris* | 7-Aug-2013 |

| | | | | | | | |
|---|---|---|---|---|---|---|---|
| RB312 | 29.508 | -81.860 | YELLOW | SOUTH | Fort McCoy, FL | *P. palustris* | 7-Aug-2013 |
| RB313 | 29.508 | -81.860 | YELLOW | SOUTH | Fort McCoy, FL | *P. palustris* | 7-Aug-2013 |
| RB314 | 29.508 | -81.860 | YELLOW | SOUTH | Fort McCoy, FL | *P. palustris* | 7-Aug-2013 |
| RB315 | 29.508 | -81.860 | YELLOW | SOUTH | Fort McCoy, FL | *P. palustris* | 7-Aug-2013 |
| RB316 | 29.508 | -81.860 | YELLOW | SOUTH | Fort McCoy, FL | *P. palustris* | 7-Aug-2013 |
| RB317 | 29.591 | -82.362 | YELLOW | SOUTH | Arrendondo, FL | *P. palustris* | 7-Aug-2013 |
| RB318 | 30.255 | -84.685 | YELLOW | SOUTH | Sopchoppy,FL | *P. palustris* | 8-Aug-2013 |
| RB319 | 30.255 | -84.685 | YELLOW | SOUTH | Sopchoppy,FL | *P. palustris* | 8-Aug-2013 |
| RB320 | 30.255 | -84.685 | YELLOW | SOUTH | Sopchoppy,FL | *P. palustris* | 8-Aug-2013 |
| RB321 | 30.255 | -84.685 | YELLOW | SOUTH | Sopchoppy,FL | *P. palustris* | 8-Aug-2013 |
| RB322 | 30.255 | -84.685 | YELLOW | SOUTH | Sopchoppy,FL | *P. palustris* | 8-Aug-2013 |
| RB323 | 30.255 | -84.685 | YELLOW | SOUTH | Sopchoppy,FL | *P. palustris* | 8-Aug-2013 |
| RB325 | 30.269 | -84.362 | YELLOW | SOUTH | Crawfordville, FL | *P. elliottii* | 8-Aug-2013 |
| RB326 | 30.255 | -84.362 | YELLOW | SOUTH | Crawfordville, FL | *P. elliottii* | 8-Aug-2013 |
| RB327 | 30.255 | -84.362 | YELLOW | SOUTH | Crawfordville, FL | *P. elliottii* | 8-Aug-2013 |
| RB328 | 30.255 | -84.362 | YELLOW | SOUTH | Crawfordville, FL | *P. elliottii* | 8-Aug-2013 |
| RB329 | 30.255 | -84.362 | YELLOW | SOUTH | Crawfordville, FL | *P. elliottii* | 8-Aug-2013 |
| RB330 | 30.315 | -84.340 | YELLOW | SOUTH | Tallahassee, FL | *P. palustris* | 8-Aug-2013 |
| RB331 | 30.333 | -84.323 | YELLOW | SOUTH | Woodville, FL | *P. palustris* | 8-Aug-2013 |
| RB332 | 30.333 | -84.323 | YELLOW | SOUTH | Woodville, FL | *P. palustris* | 8-Aug-2013 |
| RB333 | 31.090 | -86.567 | YELLOW | SOUTH | Dixie, AL | *P. elliottii* | 9-Aug-2013 |
| RB334 | 31.090 | -86.567 | YELLOW | SOUTH | Dixie, AL | *P. elliottii* | 9-Aug-2013 |
| RB335 | 38.014 | -84.504 | YELLOW | CENTRAL | Lexington KY | *P. echinata* | 22-Aug-2013 |
| RB336 | 38.014 | -84.504 | YELLOW | CENTRAL | Lexington KY | *P. echinata* | 22-Aug-2013 |
| RB337 | 38.014 | -84.504 | YELLOW | CENTRAL | Lexington KY | *P. virginiana* | 22-Aug-2013 |
| RB338 | 38.014 | -84.504 | YELLOW | CENTRAL | Lexington KY | *P. echinata* | 22-Aug-2013 |
| RB339 | 38.014 | -84.504 | YELLOW | CENTRAL | Lexington KY | *P. echinata* | 22-Aug-2013 |
| RB340 | 38.014 | -84.504 | YELLOW | CENTRAL | Lexington KY | *P. echinata* | 22-Aug-2013 |
| RB341 | 38.014 | -84.504 | YELLOW | CENTRAL | Lexington KY | *P. virginiana* | 22-Aug-2013 |

| | | | | | | | |
|---|---|---|---|---|---|---|---|
| RB342 | 38.014 | -84.504 | YELLOW | CENTRAL | Lexington KY | *P. echinata* | 9-Sep-2013 |
| RB343 | 38.014 | -84.504 | YELLOW | CENTRAL | Lexington KY | *P. rigida* | 9-Sep-2013 |
| RB344 | 38.014 | -84.504 | YELLOW | CENTRAL | Lexington KY | *P. rigida* | 9-Sep-2013 |
| RB345 | 38.014 | -84.504 | YELLOW | CENTRAL | Lexington KY | *P. rigida* | 9-Sep-2013 |
| RB346 | 38.014 | -84.504 | YELLOW | CENTRAL | Lexington KY | *P. virginiana* | 9-Sep-2013 |
| RB347 | 38.014 | -84.504 | YELLOW | CENTRAL | Lexington KY | *P. virginiana* | 9-Sep-2013 |
| RB348 | 38.014 | -84.504 | YELLOW | CENTRAL | Lexington KY | *P. virginiana* | 9-Sep-2013 |
| RB349 | 38.014 | -84.504 | YELLOW | CENTRAL | Lexington KY | *P. echinata* | 9-Sep-2013 |
| RB350 | 38.014 | -84.504 | YELLOW | CENTRAL | Lexington KY | *P. echinata* | 9-Sep-2013 |
| RB351 | 38.014 | -84.504 | YELLOW | CENTRAL | Lexington KY | *P. echinata* | 9-Sep-2013 |
| RB352 | 38.014 | -84.504 | YELLOW | CENTRAL | Lexington KY | *P. echinata* | 9-Sep-2013 |
| RB353 | 38.014 | -84.504 | YELLOW | CENTRAL | Lexington KY | *P. echinata* | 9-Sep-2013 |
| RB354 | 38.014 | -84.504 | YELLOW | CENTRAL | Lexington KY | *P. virginiana* | 9-Sep-2013 |
| RB355 | 38.014 | -84.504 | YELLOW | CENTRAL | Lexington KY | *P. virginiana* | 9-Sep-2013 |
| RB356 | 38.014 | -84.504 | YELLOW | CENTRAL | Lexington KY | *P. virginiana* | 9-Sep-2013 |
| RB357 | 38.014 | -84.504 | YELLOW | CENTRAL | Lexington KY | *P. virginiana* | 9-Sep-2013 |
| RB358 | 38.014 | -84.504 | YELLOW | CENTRAL | Lexington KY | *P. virginiana* | 9-Sep-2013 |
| RB359 | 38.014 | -84.504 | YELLOW | CENTRAL | Lexington KY | *P. virginiana* | 9-Sep-2013 |
| RB360 | 38.014 | -84.504 | YELLOW | CENTRAL | Lexington KY | *P. echinata* | 9-Sep-2013 |
| RB361 | 37.984 | -84.418 | YELLOW | CENTRAL | Lexington KY | *P. taeda* | 10-Sep-2013 |
| RB368 | 40.368 | -74.302 | WHITE | CENTRAL | Old Bridge, NJ | *P. rigida* | 20-Jul-2014 |
| RB369 | 40.368 | -74.302 | WHITE | CENTRAL | Old Bridge, NJ | *P. rigida* | 20-Jul-2014 |
| RB370 | 38.033 | -84.507 | YELLOW | CENTRAL | Lexington KY | *P. mugho* | 3-Sep-2014 |
| RB371 | 45.493 | -77.597 | YELLOW | NORTH | Wilno, ON | *P. resinosa* | 19-Aug-2014 |
| RB372 | 45.512 | -77.447 | YELLOW | NORTH | Killaloe, ON | *P. resinosa* | 19-Aug-2014 |
| RB373 | 46.471 | -82.663 | YELLOW | NORTH | Elliot Lake, ON | *P. resinosa* | 22-Aug-2014 |
| RB374 | 46.471 | -82.663 | YELLOW | NORTH | Elliot Lake, ON | *P. resinosa* | 22-Aug-2014 |
| RB375 | 46.471 | -82.663 | YELLOW | NORTH | Elliot Lake, ON | *P. resinosa* | 22-Aug-2014 |
| RB376 | 46.471 | -82.663 | YELLOW | NORTH | Elliot Lake, ON | *P. resinosa* | 22-Aug-2014 |

| | | | | | | | |
|---|---|---|---|---|---|---|---|
| RB377 | 46.471 | -82.663 | YELLOW | NORTH | Elliot Lake, ON | *P. resinosa* | 22-Aug-2014 |
| RB378 | 46.471 | -82.663 | YELLOW | NORTH | Elliot Lake, ON | *P. resinosa* | 22-Aug-2014 |
| RB379 | 46.439 | -83.227 | YELLOW | NORTH | Iron Bridge, ON | *P. resinosa* | 22-Aug-2014 |
| RB380 | 43.768 | -85.740 | YELLOW | NORTH | Brohman, MI | *P. banksiana* | 15-Jul-2015 |
| RB381 | 43.769 | -85.741 | YELLOW | NORTH | Brohman, MI | *P. banksiana* | 15-Jul-2015 |
| RB382 | 43.770 | -85.742 | YELLOW | NORTH | Bitely, MI | *P. banksiana* | 15-Jul-2015 |
| RB383 | 43.770 | -85.742 | YELLOW | NORTH | Bitely, MI | *P. banksiana* | 15-Jul-2015 |
| RB384 | 43.770 | -85.742 | YELLOW | NORTH | Bitely, MI | *P. banksiana* | 15-Jul-2015 |
| RB385 | 43.770 | -85.742 | YELLOW | NORTH | Bitely, MI | *P. banksiana* | 15-Jul-2015 |
| RB386 | 44.657 | -84.696 | YELLOW | NORTH | Grayling, MI | *P. banksiana* | 16-Jul-2015 |
| RB387 | 44.657 | -84.696 | YELLOW | NORTH | Grayling, MI | *P. banksiana* | 16-Jul-2015 |
| RB388 | 44.657 | -84.696 | YELLOW | NORTH | Grayling, MI | *P. banksiana* | 16-Jul-2015 |
| RB389 | 44.983 | -88.448 | YELLOW | NORTH | Suring, WI | *P. resinosa* | 17-Jul-2015 |
| RB390 | 44.983 | -88.448 | YELLOW | NORTH | Suring, WI | *P. resinosa* | 17-Jul-2015 |
| RB391 | 44.983 | -88.448 | YELLOW | NORTH | Suring, WI | *P. resinosa* | 17-Jul-2015 |
| RB392 | 44.983 | -88.448 | YELLOW | NORTH | Suring, WI | *P. resinosa* | 17-Jul-2015 |
| RB393 | 44.983 | -88.448 | YELLOW | NORTH | Suring, WI | *P. resinosa* | 17-Jul-2015 |
| RB394 | 44.983 | -88.448 | YELLOW | NORTH | Suring, WI | *P. resinosa* | 17-Jul-2015 |
| RB395 | 44.983 | -88.448 | YELLOW | NORTH | Suring, WI | *P. resinosa* | 17-Jul-2015 |
| RB396 | 44.156 | -90.132 | YELLOW | NORTH | Necedah, WI | *P. banksiana* | 17-Jul-2015 |
| RB397 | 44.156 | -90.132 | YELLOW | NORTH | Necedah, WI | *P. banksiana* | 17-Jul-2015 |
| RB398 | 44.156 | -90.132 | YELLOW | NORTH | Necedah, WI | *P. banksiana* | 17-Jul-2015 |
| RB399 | 44.156 | -90.132 | YELLOW | NORTH | Necedah, WI | *P. banksiana* | 17-Jul-2015 |
| RB400 | 44.156 | -90.132 | YELLOW | NORTH | Necedah, WI | *P. banksiana* | 17-Jul-2015 |
| RB401 | 44.036 | -90.082 | YELLOW | NORTH | Necedah, WI | *P. banksiana* | 17-Jul-2015 |
| RB402 | 44.036 | -90.082 | YELLOW | NORTH | Necedah, WI | *P. banksiana* | 17-Jul-2015 |
| RB403 | 44.154 | -90.132 | YELLOW | NORTH | Necedah, WI | *P. banksiana* | 17-Jul-2015 |
| RB404 | 38.186 | -83.557 | YELLOW | CENTRAL | Morehead, KY | *P. echinata* | 10-Aug-2015 |
| RB405 | 45.949 | -86.261 | YELLOW | NORTH | Manistique, MI | *P. resinosa* | 16-Aug-2015 |

| | | | | | | | |
|---|---|---|---|---|---|---|---|
| RB406 | 45.949 | -86.261 | YELLOW | NORTH | Manistique, MI | *P. resinosa* | 16-Aug-2015 |
| RB407 | 45.918 | -86.313 | YELLOW | NORTH | Thompson Township, MI | *P. banksiana* | 16-Aug-2015 |
| RB408 | 45.918 | -86.313 | YELLOW | NORTH | Thompson Township, MI | *P. banksiana* | 16-Aug-2015 |
| RB409 | 45.926 | -86.294 | YELLOW | NORTH | Thompson Township, MI | *P. resinosa* | 16-Aug-2015 |
| RB410 | 45.926 | -86.294 | YELLOW | NORTH | Thompson Township, MI | *P. resinosa* | 16-Aug-2015 |
| RB411 | 45.926 | -86.294 | YELLOW | NORTH | Thompson Township, MI | *P. resinosa* | 16-Aug-2015 |
| RB412 | 45.926 | -86.294 | YELLOW | NORTH | Thompson Township, MI | *P. resinosa* | 16-Aug-2015 |
| RB413 | 45.926 | -86.294 | YELLOW | NORTH | Thompson Township, MI | *P. resinosa* | 16-Aug-2015 |
| RB414 | 45.926 | -86.294 | YELLOW | NORTH | Thompson Township, MI | *P. resinosa* | 16-Aug-2015 |
| RB415 | 45.926 | -86.294 | YELLOW | NORTH | Thompson Township, MI | *P. resinosa* | 16-Aug-2015 |
| RB416 | 44.983 | -88.448 | YELLOW | NORTH | Suring, WI | *P. resinosa* | 17-Aug-2015 |
| RB417 | 44.983 | -88.448 | YELLOW | NORTH | Suring, WI | *P. resinosa* | 17-Aug-2015 |
| RB418 | 44.983 | -88.448 | YELLOW | NORTH | Suring, WI | *P. resinosa* | 17-Aug-2015 |
| RB419 | 44.983 | -88.448 | YELLOW | NORTH | Suring, WI | *P. resinosa* | 17-Aug-2015 |
| RB420 | 44.983 | -88.448 | YELLOW | NORTH | Suring, WI | *P. resinosa* | 17-Aug-2015 |
| RB421 | 44.983 | -88.448 | YELLOW | NORTH | Suring, WI | *P. resinosa* | 17-Aug-2015 |
| RB425 | 44.862 | -89.637 | YELLOW | NORTH | Rothschild, WI | *P. banksiana* | 9-Jul-2016 |
| RB426 | 44.862 | -89.637 | YELLOW | NORTH | Rothschild, WI | *P. banksiana* | 9-Jul-2016 |
| RB427 | 44.862 | -89.637 | YELLOW | NORTH | Rothschild, WI | *P. banksiana* | 9-Jul-2016 |
| RB428 | 44.862 | -89.637 | YELLOW | NORTH | Rothschild, WI | *P. banksiana* | 9-Jul-2016 |
| RB429 | 44.862 | -89.637 | YELLOW | NORTH | Rothschild, WI | *P. banksiana* | 9-Jul-2016 |
| RB430 | 44.862 | -89.637 | YELLOW | NORTH | Rothschild, WI | *P. banksiana* | 9-Jul-2016 |
| RB431 | 44.844 | -89.691 | YELLOW | NORTH | Mosinee, WI | *P. resinosa* | 9-Jul-2016 |
| RB432 | 44.844 | -89.691 | YELLOW | NORTH | Mosinee, WI | *P. resinosa* | 9-Jul-2016 |
| RB433 | 44.844 | -89.691 | YELLOW | NORTH | Mosinee, WI | *P. banksiana* | 9-Jul-2016 |
| RB434 | 44.844 | -89.691 | YELLOW | NORTH | Mosinee, WI | *P. banksiana* | 9-Jul-2016 |
| RB435 | 44.844 | -89.691 | YELLOW | NORTH | Mosinee, WI | *P. banksiana* | 9-Jul-2016 |
| RB437 | 44.026 | -89.705 | YELLOW | NORTH | Friendship, WI | *P. banksiana* | 9-Jul-2016 |
| RB438 | 44.026 | -89.705 | YELLOW | NORTH | Friendship, WI | *P. banksiana* | 9-Jul-2016 |

| | | | | | | | |
|---|---|---|---|---|---|---|---|
| RB439 | 44.026 | -89.705 | YELLOW | NORTH | Friendship, WI | *P. banksiana* | 9-Jul-2016 |
| RB440 | 44.026 | -89.705 | YELLOW | NORTH | Friendship, WI | *P. banksiana* | 9-Jul-2016 |
| RB441 | 44.036 | -90.082 | YELLOW | NORTH | Necedah, WI | *P. banksiana* | 9-Jul-2016 |
| RB442 | 44.036 | -90.082 | YELLOW | NORTH | Necedah, WI | *P. banksiana* | 9-Jul-2016 |
| RB443 | 44.036 | -90.082 | YELLOW | NORTH | Necedah, WI | *P. banksiana* | 9-Jul-2016 |
| RB444 | 44.731 | -84.749 | YELLOW | NORTH | Frederic, MI | *P. resinosa* | 23-Jul-2016 |
| RB445 | 44.731 | -84.749 | YELLOW | NORTH | Frederic, MI | *P. resinosa* | 23-Jul-2016 |
| RB446 | 44.731 | -84.749 | YELLOW | NORTH | Frederic, MI | *P. resinosa* | 23-Jul-2016 |
| RB447 | 44.731 | -84.749 | YELLOW | NORTH | Frederic, MI | *P. resinosa* | 23-Jul-2016 |
| RB448 | 44.731 | -84.749 | YELLOW | NORTH | Frederic, MI | *P. resinosa* | 23-Jul-2016 |
| RB449 | 44.731 | -84.749 | YELLOW | NORTH | Frederic, MI | *P. resinosa* | 23-Jul-2016 |
| RB450 | 44.731 | -84.749 | YELLOW | NORTH | Frederic, MI | *P. resinosa* | 23-Jul-2016 |
| RB451 | 44.720 | -84.745 | YELLOW | NORTH | Frederic, MI | *P. resinosa* | 23-Jul-2016 |
| RB452 | 44.720 | -84.745 | YELLOW | NORTH | Frederic, MI | *P. resinosa* | 23-Jul-2016 |
| RB453 | 44.720 | -84.745 | YELLOW | NORTH | Frederic, MI | *P. resinosa* | 23-Jul-2016 |
| RB454 | 44.731 | -84.749 | YELLOW | NORTH | Frederic, MI | *P. resinosa* | 23-Jul-2016 |
| RB455 | 44.731 | -84.749 | YELLOW | NORTH | Frederic, MI | *P. resinosa* | 23-Jul-2016 |
| RB456 | 44.731 | -84.749 | YELLOW | NORTH | Frederic, MI | *P. resinosa* | 23-Jul-2016 |
| RB457 | 44.731 | -84.749 | YELLOW | NORTH | Frederic, MI | *P. resinosa* | 23-Jul-2016 |
| RB458 | 44.731 | -84.749 | YELLOW | NORTH | Frederic, MI | *P. resinosa* | 23-Jul-2016 |
| RB459 | 44.731 | -84.749 | YELLOW | NORTH | Frederic, MI | *P. resinosa* | 23-Jul-2016 |
| RB460 | 44.731 | -84.749 | YELLOW | NORTH | Frederic, MI | *P. resinosa* | 23-Jul-2016 |
| RB461 | 44.731 | -84.749 | YELLOW | NORTH | Frederic, MI | *P. resinosa* | 23-Jul-2016 |
| RB462 | 44.731 | -84.749 | YELLOW | NORTH | Frederic, MI | *P. banksiana* | 23-Jul-2016 |
| RB463 | 44.731 | -84.749 | YELLOW | NORTH | Frederic, MI | *P. banksiana* | 23-Jul-2016 |
| RB464 | 44.731 | -84.749 | YELLOW | NORTH | Frederic, MI | *P. banksiana* | 23-Jul-2016 |
| RB465 | 44.731 | -84.749 | YELLOW | NORTH | Frederic, MI | *P. banksiana* | 23-Jul-2016 |
| RB466 | 44.731 | -84.749 | YELLOW | NORTH | Frederic, MI | *P. banksiana* | 23-Jul-2016 |
| RB467 | 44.731 | -84.749 | YELLOW | NORTH | Frederic, MI | *P. banksiana* | 23-Jul-2016 |

| | | | | | | | |
|---|---|---|---|---|---|---|---|
| RB468 | 44.731 | -84.749 | YELLOW | NORTH | Frederic, MI | *P. banksiana* | 23-Jul-2016 |
| RB469 | 44.731 | -84.749 | YELLOW | NORTH | Frederic, MI | *P. banksiana* | 23-Jul-2016 |
| RB470 | 44.731 | -84.749 | YELLOW | NORTH | Frederic, MI | *P. banksiana* | 23-Jul-2016 |
| RB471 | 44.731 | -84.749 | YELLOW | NORTH | Frederic, MI | *P. banksiana* | 23-Jul-2016 |
| RB472 | 44.731 | -84.749 | YELLOW | NORTH | Frederic, MI | *P. banksiana* | 23-Jul-2016 |
| RB473 | 44.731 | -84.749 | YELLOW | NORTH | Frederic, MI | *P. banksiana* | 23-Jul-2016 |
| RB474 | 44.731 | -84.749 | YELLOW | NORTH | Frederic, MI | *P. banksiana* | 23-Jul-2016 |
| RB475 | 44.731 | -84.749 | YELLOW | NORTH | Frederic, MI | *P. banksiana* | 23-Jul-2016 |
| RB476 | 44.731 | -84.749 | YELLOW | NORTH | Frederic, MI | *P. banksiana* | 23-Jul-2016 |
| RB477 | 44.731 | -84.749 | YELLOW | NORTH | Frederic, MI | *P. banksiana* | 23-Jul-2016 |
| RB478 | 44.731 | -84.749 | YELLOW | NORTH | Frederic, MI | *P. banksiana* | 23-Jul-2016 |
| RB479 | 44.731 | -84.749 | YELLOW | NORTH | Frederic, MI | *P. banksiana* | 23-Jul-2016 |
| RB480 | 46.096 | -85.394 | YELLOW | NORTH | Naubinway, MI | *P. resinosa* | 23-Jul-2016 |
| RB481 | 46.096 | -85.394 | YELLOW | NORTH | Naubinway, MI | *P. banksiana* | 23-Jul-2016 |
| RB482 | 46.096 | -85.394 | YELLOW | NORTH | Naubinway, MI | *P. banksiana* | 23-Jul-2016 |
| RB483 | 46.096 | -85.394 | YELLOW | NORTH | Naubinway, MI | *P. banksiana* | 23-Jul-2016 |
| RB484 | 44.731 | -84.749 | YELLOW | NORTH | Frederic, MI | *P. resinosa* | 23-Jul-2016 |
| RB485 | 44.731 | -84.749 | YELLOW | NORTH | Frederic, MI | *P. resinosa* | 23-Jul-2016 |

**Table S5 – Sequences for adapters containing variable-length barcodes from Burford Reiskind *et al.* (2016).**

| Barcode | P1.1 sequence | P1.2 sequence |
|---|---|---|
| ATTAT | ACACTCTTTCCCTACACGACGCTCTTCCGATCTATTATCATG | /5Phos/ATAATAGATCGGAAGAGCGTCGTGTAGGGAAAGAGTGT |
| CACCA | ACACTCTTTCCCTACACGACGCTCTTCCGATCTCACCACATG | /5Phos/TGGTGAGATCGGAAGAGCGTCGTGTAGGGAAAGAGTGT |
| CCTCG | ACACTCTTTCCCTACACGACGCTCTTCCGATCTCCTCGCATG | /5Phos/CGAGGAGATCGGAAGAGCGTCGTGTAGGGAAAGAGTGT |
| CTTGA | ACACTCTTTCCCTACACGACGCTCTTCCGATCTCTTGACATG | /5Phos/TCAAGAGATCGGAAGAGCGTCGTGTAGGGAAAGAGTGT |
| GGATA | ACACTCTTTCCCTACACGACGCTCTTCCGATCTGGATACATG | /5Phos/TATCCAGATCGGAAGAGCGTCGTGTAGGGAAAGAGTGT |
| GGTGT | ACACTCTTTCCCTACACGACGCTCTTCCGATCTGGTGTCATG | /5Phos/ACACCAGATCGGAAGAGCGTCGTGTAGGGAAAGAGTGT |
| TATGT | ACACTCTTTCCCTACACGACGCTCTTCCGATCTTATGTCATG | /5Phos/ACATAAGATCGGAAGAGCGTCGTGTAGGGAAAGAGTGT |
| TGCTT | ACACTCTTTCCCTACACGACGCTCTTCCGATCTTGCTTCATG | /5Phos/AAGCAAGATCGGAAGAGCGTCGTGTAGGGAAAGAGTGT |
| AACTGG | ACACTCTTTCCCTACACGACGCTCTTCCGATCTAACTGGCATG | /5Phos/CCAGTTAGATCGGAAGAGCGTCGTGTAGGGAAAGAGTGT |
| ACAACT | ACACTCTTTCCCTACACGACGCTCTTCCGATCTACAACTCATG | /5Phos/AGTTGTAGATCGGAAGAGCGTCGTGTAGGGAAAGAGTGT |
| ATAGAT | ACACTCTTTCCCTACACGACGCTCTTCCGATCTATAGATCATG | /5Phos/ATCTATAGATCGGAAGAGCGTCGTGTAGGGAAAGAGTGT |
| CAGATA | ACACTCTTTCCCTACACGACGCTCTTCCGATCTCAGATACATG | /5Phos/TATCTGAGATCGGAAGAGCGTCGTGTAGGGAAAGAGTGT |
| GAAGTG | ACACTCTTTCCCTACACGACGCTCTTCCGATCTGAAGTGCATG | /5Phos/CACTTCAGATCGGAAGAGCGTCGTGTAGGGAAAGAGTGT |
| GGCTTA | ACACTCTTTCCCTACACGACGCTCTTCCGATCTGGCTTACATG | /5Phos/TAAGCCAGATCGGAAGAGCGTCGTGTAGGGAAAGAGTGT |
| TCTTGG | ACACTCTTTCCCTACACGACGCTCTTCCGATCTTCTTGGCATG | /5Phos/CCAAGAAGATCGGAAGAGCGTCGTGTAGGGAAAGAGTGT |
| TCACTG | ACACTCTTTCCCTACACGACGCTCTTCCGATCTTCACTGCATG | /5Phos/CAGTGAAGATCGGAAGAGCGTCGTGTAGGGAAAGAGTGT |
| ACCAGGA | ACACTCTTTCCCTACACGACGCTCTTCCGATCTACCAGGACATG | /5Phos/TCCTGGTAGATCGGAAGAGCGTCGTGTAGGGAAAGAGTGT |
| CCACTCA | ACACTCTTTCCCTACACGACGCTCTTCCGATCTCCACTCACATG | /5Phos/TGAGTGGAGATCGGAAGAGCGTCGTGTAGGGAAAGAGTGT |
| CCGAACA | ACACTCTTTCCCTACACGACGCTCTTCCGATCTCCGAACACATG | /5Phos/TGTTCGGAGATCGGAAGAGCGTCGTGTAGGGAAAGAGTGT |
| CTAAGCA | ACACTCTTTCCCTACACGACGCTCTTCCGATCTCTAAGCACATG | /5Phos/TGCTTAGAGATCGGAAGAGCGTCGTGTAGGGAAAGAGTGT |
| CTCGCGG | ACACTCTTTCCCTACACGACGCTCTTCCGATCTCTCGCGGCATG | /5Phos/CCGCGAGAGATCGGAAGAGCGTCGTGTAGGGAAAGAGTGT |
| GCGTCCT | ACACTCTTTCCCTACACGACGCTCTTCCGATCTGCGTCCTCATG | /5Phos/AGGACGCAGATCGGAAGAGCGTCGTGTAGGGAAAGAGTGT |
| GGAACGA | ACACTCTTTCCCTACACGACGCTCTTCCGATCTGGAACGACATG | /5Phos/TCGTTCCAGATCGGAAGAGCGTCGTGTAGGGAAAGAGTGT |
| TAGCCAA | ACACTCTTTCCCTACACGACGCTCTTCCGATCTTAGCCAACATG | /5Phos/TTGGCTAAGATCGGAAGAGCGTCGTGTAGGGAAAGAGTGT |

| | | |
|---|---|---|
| ACTGCGAT | ACACTCTTTCCCTACACGACGCTCTTCCGATCTACTGCGATCATG | /5Phos/ATCGCAGTAGATCGGAAGAGCGTCGTGTAGGGAAAGAGTGT |
| ATGAGCAA | ACACTCTTTCCCTACACGACGCTCTTCCGATCTATGAGCAACATG | /5Phos/TTGCTCATAGATCGGAAGAGCGTCGTGTAGGGAAAGAGTGT |
| GCCTACCT | ACACTCTTTCCCTACACGACGCTCTTCCGATCTGCCTACCTCATG | /5Phos/AGGTAGGCAGATCGGAAGAGCGTCGTGTAGGGAAAGAGTGT |
| TAGCGGAT | ACACTCTTTCCCTACACGACGCTCTTCCGATCTTAGCGGATCATG | /5Phos/ATCCGCTAAGATCGGAAGAGCGTCGTGTAGGGAAAGAGTGT |
| TGACGCCA | ACACTCTTTCCCTACACGACGCTCTTCCGATCTTGACGCCACATG | /5Phos/TGGCGTCAAGATCGGAAGAGCGTCGTGTAGGGAAAGAGTGT |
| ACGGTACT | ACACTCTTTCCCTACACGACGCTCTTCCGATCTACGGTACTCATG | /5Phos/AGTACCGTAGATCGGAAGAGCGTCGTGTAGGGAAAGAGTGT |
| AAGACGCT | ACACTCTTTCCCTACACGACGCTCTTCCGATCTAAGACGCTCATG | /5Phos/AGCGTCTTAGATCGGAAGAGCGTCGTGTAGGGAAAGAGTGT |
| TCAGAGAT | ACACTCTTTCCCTACACGACGCTCTTCCGATCTTCAGAGATCATG | /5Phos/ATCTCTGAAGATCGGAAGAGCGTCGTGTAGGGAAAGAGTGT |
| ATATCGCCA | ACACTCTTTCCCTACACGACGCTCTTCCGATCTATATCGCCACATG | /5Phos/TGGCGATATAGATCGGAAGAGCGTCGTGTAGGGAAAGAGTGT |
| GAGCGACAT | ACACTCTTTCCCTACACGACGCTCTTCCGATCTGAGCGACATCATG | /5Phos/ATGTCGCTCAGATCGGAAGAGCGTCGTGTAGGGAAAGAGTGT |
| GCAAGCCAT | ACACTCTTTCCCTACACGACGCTCTTCCGATCTGCAAGCCATCATG | /5Phos/ATGGCTTGCAGATCGGAAGAGCGTCGTGTAGGGAAAGAGTGT |
| AACGTGCCT | ACACTCTTTCCCTACACGACGCTCTTCCGATCTAACGTGCCTCATG | /5Phos/AGGCACGTTAGATCGGAAGAGCGTCGTGTAGGGAAAGAGTGT |
| TATTCGCAT | ACACTCTTTCCCTACACGACGCTCTTCCGATCTTATTCGCATCATG | /5Phos/ATGCGAATAAGATCGGAAGAGCGTCGTGTAGGGAAAGAGTGT |
| TCACGGAAG | ACACTCTTTCCCTACACGACGCTCTTCCGATCTTCACGGAAGCATG | /5Phos/CTTCCGTGAAGATCGGAAGAGCGTCGTGTAGGGAAAGAGTGT |
| TGGCACAGA | ACACTCTTTCCCTACACGACGCTCTTCCGATCTTGGCACAGACATG | /5Phos/TCTGTGCCAAGATCGGAAGAGCGTCGTGTAGGGAAAGAGTGT |
| CTCTCGCAT | ACACTCTTTCCCTACACGACGCTCTTCCGATCTCTCTCGCATCATG | /5Phos/ATGCGAGAGAGATCGGAAGAGCGTCGTGTAGGGAAAGAGTGT |
| AACGCACATT | ACACTCTTTCCCTACACGACGCTCTTCCGATCTAACGCACATTCATG | /5Phos/AATGTGCGTTAGATCGGAAGAGCGTCGTGTAGGGAAAGAGTGT |
| CCTTGCCATT | ACACTCTTTCCCTACACGACGCTCTTCCGATCTCCTTGCCATTCATG | /5Phos/AATGGCAAGGAGATCGGAAGAGCGTCGTGTAGGGAAAGAGTGT |
| CGTCGCCACT | ACACTCTTTCCCTACACGACGCTCTTCCGATCTCGTCGCCACTCATG | /5Phos/AGTGGCGACGAGATCGGAAGAGCGTCGTGTAGGGAAAGAGTGT |
| CGTGGACAGT | ACACTCTTTCCCTACACGACGCTCTTCCGATCTCGTGGACAGTCATG | /5Phos/ACTGTCCACGAGATCGGAAGAGCGTCGTGTAGGGAAAGAGTGT |
| GGTGCACATT | ACACTCTTTCCCTACACGACGCTCTTCCGATCTGGTGCACATTCATG | /5Phos/AATGTGCACCAGATCGGAAGAGCGTCGTGTAGGGAAAGAGTGT |
| TGGCAACAGA | ACACTCTTTCCCTACACGACGCTCTTCCGATCTTGGCAACAGACATG | /5Phos/TCTGTTGCCAAGATCGGAAGAGCGTCGTGTAGGGAAAGAGTGT |
| ACAACCAACT | ACACTCTTTCCCTACACGACGCTCTTCCGATCTACAACCAACTCATG | /5Phos/AGTTGGTTGTAGATCGGAAGAGCGTCGTGTAGGGAAAGAGTGT |
| CAACCACACA | ACACTCTTTCCCTACACGACGCTCTTCCGATCTCAACCACACACATG | /5Phos/TGTGTGGTTGAGATCGGAAGAGCGTCGTGTAGGGAAAGAGTGT |

**Table S6 – Sequences for PCR primers, including degenerate bases.** PCR1 is the universal primer used for all reactions, and the PCR2 primers contain their respective Illumina index sequences in their "Name". When ordering, degenerate bases should be hand-mixed to ensure equal base composition.

| **Name** | **Sequence** |
|---|---|
| PCR1 | AATGATACGGCGACCACCGAGATCTACACTCTTTCCCTACACGACG |
| PCR2_ATCACGAT | CAAGCAGAAGACGGCATACGAGATNNNNATCGTGATGTGACTGGAGTTCAGACGTGTGC |
| PCR2_CGATGTAT | CAAGCAGAAGACGGCATACGAGATNNNNATACATCGGTGACTGGAGTTCAGACGTGTGC |
| PCR2_TTAGGCAT | CAAGCAGAAGACGGCATACGAGATNNNNATGCCTAAGTGACTGGAGTTCAGACGTGTGC |
| PCR2_TGACCAAT | CAAGCAGAAGACGGCATACGAGATNNNNATTGGTCAGTGACTGGAGTTCAGACGTGTGC |
| PCR2_ACAGTGAT | CAAGCAGAAGACGGCATACGAGATNNNNATCACTGTGTGACTGGAGTTCAGACGTGTGC |
| PCR2_GGCTACAT | CAAGCAGAAGACGGCATACGAGATNNNNATGTAGCCGTGACTGGAGTTCAGACGTGTGC |
| PCR2_AGTCAACA | CAAGCAGAAGACGGCATACGAGATNNNNTGTTGACTGTGACTGGAGTTCAGACGTGTGC |
| PCR2_CCGTCCCG | CAAGCAGAAGACGGCATACGAGATNNNNCGGGACGGGTGACTGGAGTTCAGACGTGTGC |
| PCR2_GTCCGCAC | CAAGCAGAAGACGGCATACGAGATNNNNGTGCGGACGTGACTGGAGTTCAGACGTGTGC |
| PCR2_GTGAAACG | CAAGCAGAAGACGGCATACGAGATNNNNCGTTTCACGTGACTGGAGTTCAGACGTGTGC |
| PCR2_GTGGCCTT | CAAGCAGAAGACGGCATACGAGATNNNNAAGGCCACGTGACTGGAGTTCAGACGTGTGC |
| PCR2_GTTTCGGA | CAAGCAGAAGACGGCATACGAGATNNNNTCCGAAACGTGACTGGAGTTCAGACGTGTGC |

**Figure S3. α score plot for DAPC.** The optimal number of PCs to retain (1) is indicated by a red circle.

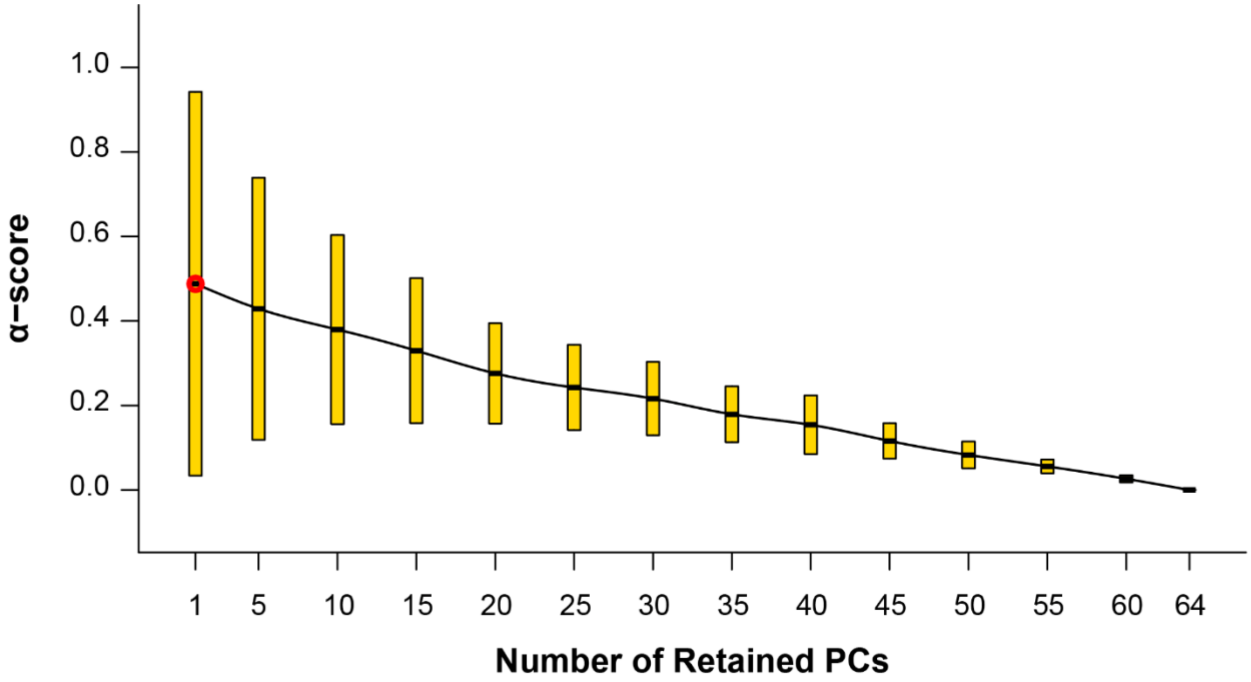

**Figure S4. Variance explained across PCs.**

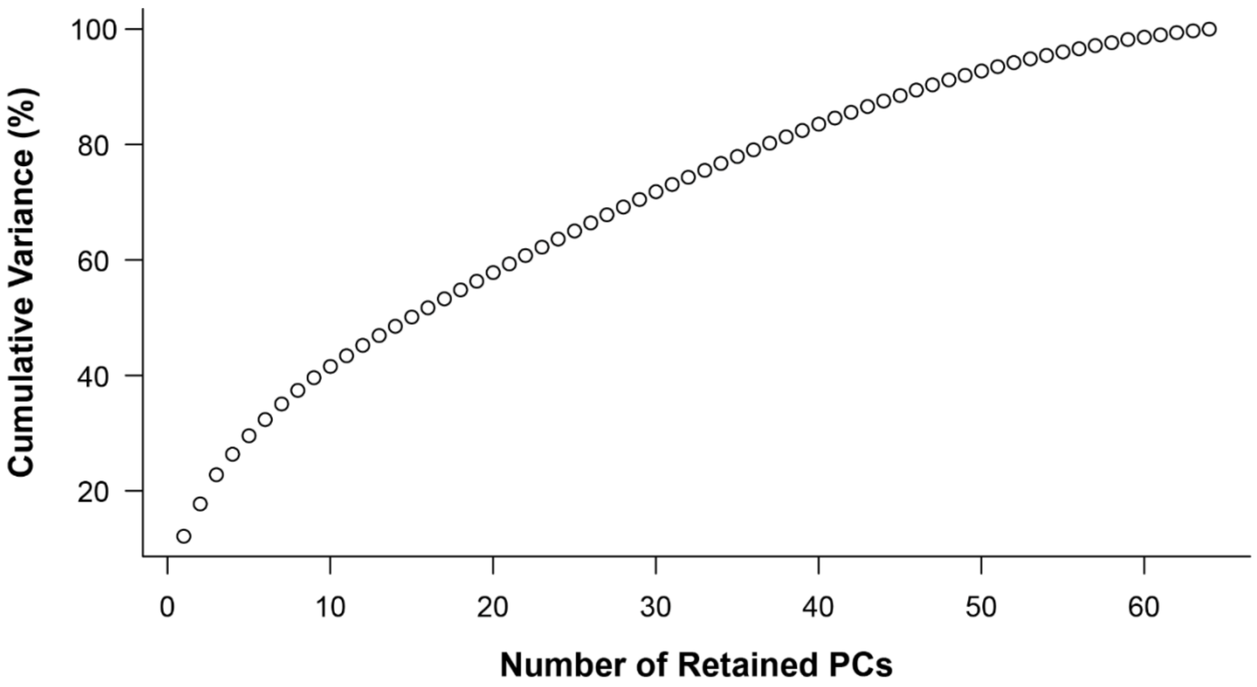

**Table S7. Population assignments for *K*=2.** For each individual, sampling location, colony body color, ancestry proportions (averaged across 100 ADMIXTURE runs), and assignment probabilities (DAPC, 1 PC) are given.

| | | | ADMIXTURE | | 1-PC DAPC | |
|---|---|---|---|---|---|---|
| **ID** | **Place** | **Color** | **East** | **West** | **East** | **West** |
| LL004_01 | Lexington, KY | Yellow | 0.000 | 1.000 | 0.000 | 1.000 |
| LL005_01 | Lexington, KY | Yellow | 0.000 | 1.000 | 0.000 | 1.000 |
| LL010 | Bronston, KY | Yellow | 0.230 | 0.770 | 0.005 | 0.995 |
| LL011 | London, KY | Yellow | 0.191 | 0.809 | 0.000 | 1.000 |
| LL013 | London, KY | Yellow | 0.227 | 0.773 | 0.001 | 0.999 |
| LL014 | London, KY | Yellow | 0.199 | 0.801 | 0.035 | 0.965 |
| LL015_01 | London, KY | Yellow | 0.208 | 0.792 | 0.001 | 0.999 |
| LL042 | Ford, VA | White | 0.953 | 0.047 | 1.000 | 0.000 |
| LL058_2 | Lexington, KY | Yellow | 0.000 | 1.000 | 0.000 | 1.000 |
| LL062_1 | Lexington, KY | Yellow | 0.000 | 1.000 | 0.000 | 1.000 |
| LL064 | Lexington, KY | Yellow | 0.000 | 1.000 | 0.000 | 1.000 |
| LL070_1 | Lexington, KY | Yellow | 0.000 | 1.000 | 0.000 | 1.000 |
| LL073_2R | Lexington, KY | Yellow | 0.000 | 1.000 | 0.000 | 1.000 |
| LL074_5 | Lexington, KY | Yellow | 0.000 | 1.000 | 0.000 | 1.000 |
| LL084 | West Yarmouth, MA | White | 1.000 | 0.000 | 1.000 | 0.000 |
| LL084_02 | West Yarmouth, MA | White | 1.000 | 0.000 | 1.000 | 0.000 |
| LL084_03 | West Yarmouth, MA | White | 1.000 | 0.000 | 1.000 | 0.000 |
| LL102 | Lexington, KY | Yellow | 0.000 | 1.000 | 0.000 | 1.000 |
| LL137 | Lexington, KY | Yellow | 0.000 | 1.000 | 0.000 | 1.000 |
| LL142_02 | Crossville, TN | Yellow | 0.497 | 0.503 | 0.944 | 0.056 |
| LL144_01 | Crossville, TN | Yellow | 0.519 | 0.481 | 0.917 | 0.083 |
| LL180_02 | Goshen, KY | Yellow | 0.033 | 0.967 | 0.155 | 0.845 |
| LL181 | Goshen, KY | Yellow | 0.150 | 0.850 | 0.001 | 0.999 |
| LL194_02 | Stanton, KY | Yellow | 0.100 | 0.900 | 0.000 | 1.000 |
| LL195_02 | Stanton, KY | Yellow | 0.090 | 0.910 | 0.000 | 1.000 |
| RB020Db | London, KY | Yellow | 0.193 | 0.807 | 0.001 | 0.999 |
| RB028_01 | Egg Harbor, NJ | White | 1.000 | 0.000 | 1.000 | 0.000 |
| RB076_01 | Lexington, KY | Yellow | 0.000 | 1.000 | 0.000 | 1.000 |
| RB107_01 | Mountain Grove, VA | White | 0.652 | 0.348 | 0.986 | 0.014 |
| RB108_01 | Deer Run, WV | White | 0.711 | 0.289 | 0.998 | 0.002 |
| RB110.01 | Browns Mill, NJ | White | 1.000 | 0.000 | 1.000 | 0.000 |
| RB112_01 | Tuckerton, NJ | White | 1.000 | 0.000 | 1.000 | 0.000 |
| RB118_01 | Brandywine, WV | White | 0.756 | 0.244 | 0.997 | 0.003 |
| RB119_01 | Buena Vista, VA | White | 0.813 | 0.187 | 1.000 | 0.000 |
| RB126_01 | Lexington, KY | Yellow | 0.000 | 1.000 | 0.000 | 1.000 |
| RB164 | Tyron, NC | Yellow | 0.873 | 0.127 | 1.000 | 0.000 |
| RB165 | Tyron, NC | Yellow | 0.867 | 0.133 | 1.000 | 0.000 |

| | | | | | | |
|---|---|---|---|---|---|---|
| RB167_01 | Chesnee, SC | Yellow | 0.859 | 0.141 | 1.000 | 0.000 |
| RB190 | Pickens, SC | Yellow | 0.809 | 0.191 | 1.000 | 0.000 |
| RB220 | Sandy Ridge, NC | Yellow | 0.859 | 0.141 | 1.000 | 0.000 |
| RB221 | Sandy Ridge, NC | Yellow | 0.849 | 0.151 | 1.000 | 0.000 |
| RB222 | Sandy Ridge, NC | Yellow | 0.839 | 0.161 | 1.000 | 0.000 |
| RB223 | Bassett, VA | Mixed | 0.824 | 0.176 | 1.000 | 0.000 |
| RB224 | Penhook, VA | White | 0.925 | 0.075 | 1.000 | 0.000 |
| RB225 | Penhook, VA | White | 0.927 | 0.073 | 1.000 | 0.000 |
| RB226_old | Gretna, VA | Mixed | 0.919 | 0.081 | 1.000 | 0.000 |
| RB227 | Amelia, VA | White | 0.952 | 0.048 | 1.000 | 0.000 |
| RB228_new | Amelia, VA | White | 1.000 | 0.000 | 1.000 | 0.000 |
| RB229 | Amelia, VA | White | 0.956 | 0.044 | 1.000 | 0.000 |
| RB286 | Clear Springs, MD | Yellow | 0.882 | 0.118 | 1.000 | 0.000 |
| RB287 | Clear Springs, MD | Yellow | 1.000 | 0.000 | 1.000 | 0.000 |
| RB288_02 | Clear Springs, MD | Yellow | 1.000 | 0.000 | 1.000 | 0.000 |
| RB289 | Clear Springs, MD | Yellow | 1.000 | 0.000 | 1.000 | 0.000 |
| RB290 | Old Bridge, NJ | White | 1.000 | 0.000 | 1.000 | 0.000 |
| RB304 | Ossippee, NH | Yellow | 1.000 | 0.000 | 1.000 | 0.000 |
| RB305 | Ossippee, NH | Yellow | 1.000 | 0.000 | 1.000 | 0.000 |
| RB306 | Ossippee, NH | Yellow | 1.000 | 0.000 | 1.000 | 0.000 |
| RB307 | Ossippee, NH | Yellow | 1.000 | 0.000 | 1.000 | 0.000 |
| RB308 | Ossippee, NH | Yellow | 1.000 | 0.000 | 1.000 | 0.000 |
| RB339 | Lexington, KY | Yellow | 0.000 | 1.000 | 0.000 | 1.000 |
| RB343_01 | Lexington, KY | Yellow | 0.000 | 1.000 | 0.000 | 1.000 |
| RB355 | Lexington, KY | Yellow | 0.000 | 1.000 | 0.000 | 1.000 |
| RB368 | Old Bridge, NJ | White | 1.000 | 0.000 | 1.000 | 0.000 |
| RB369 | Old Bridge, NJ | White | 1.000 | 0.000 | 1.000 | 0.000 |
| RB404 | Morehead, KY | Yellow | 0.188 | 0.812 | 0.331 | 0.669 |
